# CCM signaling complex (CSC) coupling both classic and non-classic progesterone receptor signaling

**DOI:** 10.1101/2020.05.24.112847

**Authors:** Johnathan Abou-Fadel, Xiaoting Jiang, Brian Grajeda, Akhil Padarti, Cameron C. Ellis, Jun Zhang

## Abstract

Excessive progesterone (PRG) may increase breast cancer risk under hormone therapy during postmenopause or hormonal contraception. As a sex steroid hormone, PRG exerts its cellular responses through signaling cascades involving classic (genomic), non-classic (non-genomic), or both combined responses by binding to either classic nuclear PRG receptors or non-classic membrane PRG receptors. Currently, the intricate balance and switch mechanisms between these two signaling cascades remain elusive. Three genes, KRIT1 (CCM1), MGC4607 (CCM2), and PDCD10 (CCM3), have been demonstrated to form a CCM signaling complex (CSC). In this report, we discover that the CSC plays an essential role in coupling both classic and non-classic PRG signaling pathways by mediating crosstalk between them. The coupled signaling pathways were detailed through high throughput omics.

**One Sentence Summary:** We discover a novel signaling network among the CCM signaling complex (CSC), classic and non-classic progesterone receptors, and their ligands-progesterone/mifepristone, is dynamically modulated and fine-tuned with a series of feedback regulations; perturbation of this intricate balance, such as hormone therapy in the postmenopause or hormonal contraception regimen, or perturbed CSC signaling could result in increased risks in breast cancer or compromising tumor therapy.

---

Excessive progesterone (PRG) may increase breast cancer risk under hormone therapy during postmenopause or hormonal contraception. As a sex steroid hormone, PRG exerts its cellular responses through signaling cascades involving classic (genomic), non-classic (non-genomic), or both combined responses by binding to either classic nuclear PRG receptors or non-classic membrane PRG receptors (*1*). It has been well-defined that PRG commonly binds to its nuclear receptor (PR) as transcriptional factors in classic genomic actions (*2*). PR has two major isoforms (PR1/2) which are alternative transcripts with quite different cellular functions (*2*). PRG can evoke both genomic and non-genomic actions of the classical PR1/2. In non-genomic responses, PRG binds PR1/2, leading to rapid changes in intracellular calcium/chloride levels in a rapid non-classical mechanism (*3*), which is defined as membrane-initiated (non-genomic) actions. Recently, two new groups of membrane-bound PRG receptors which are unrelated to PR1/2 have been identified: 1). 5 members of membrane progestin receptors (mPRs)/the Class II progestin and adipoQ receptor (PAQRs); 2). 4 members of the b5-like heme/steroid-binding protein family known as sigma2 receptor (S2R)/progesterone receptor membrane components (PGRMCs) (*4, 5*). It has been shown that mPRs are highly expressed in reproductive tissues and function by coupling to G proteins (*6*) to execute their rapid non-classical actions. Homozygous mutant fish of all mPR genes were generated with no phenotype, suggesting redundancy shared by mPR genes (*7*).

Three CCM genes, KRIT1 (CCM1), MGC4607 (CCM2), and PDCD10 (CCM3), form the CCM signaling complex (CSC) that mediates multiple signaling (*8–10*). Three CCM genes are differentially expressed among various tissues/cells (*9, 11*); and this differential expression of CCM proteins becomes more dramatic in various cancers. Nearly half of CCMs gene expression changes (44%) were found in reproductive tumors (*9*), suggesting that the CSC might play a significant role in reproductive tumorigenesis (*9–13*). Among all reproductive cancers with altered expression of CCM proteins, breast tumors seemed to have the most drastic changes (*12*). In this report, we discovered that the CSC plays an essential role in coupling both classic and non-classic PRG receptors by mediating crosstalk between them. Depleting any of three CCMs genes results in the disruption of either classic or non-classic PRG receptors-mediated signaling; similarly, silencing any of classic or non-classic PRG receptors leads to disruption of the CSC complex. Furthermore, utilizing high throughput omic approaches, we have elucidated alterations in signaling mechanisms under hormone therapy or disrupted CSC conditions affecting numerous essential signaling cascades. These signaling pathways include P53 signaling, P13K-AKT signaling, cell cycle, apoptosis, WNT, MAPK, various pathways in cancer as well as steroid hormone biosynthesis signaling pathways, solidifying the CSC’s novel role in tumorigenesis.

## Results

### Upregulation of CCM proteins suggest their potential involvement in breast tumorigenesis

#### Significantly increased expression levels of CCM proteins in breast cancer tissues

To validate our findings that CCM2 expression is most frequently perturbed in breast tumors (*9, 12*), we investigated additional breast cancer tissue-pairs using different panels with a larger sample size. Increased CCM2 expression was observed in breast tumor tissue compared to normal tissue (Left and middle panels, Fig. 1A), with statistical significance in the entire collection (right panel, Fig. 1A). We next examined the protein co-expression levels of CCM1/3 proteins in identified human breast cancers using immunofluorescence (IF) imaging. The coordinated significant increase of both CCM1/3 proteins was observed in breast tumors compared to normal tissues (Fig. 1B). Western blots further validated our IF imaging data (Fig. 1C). Furthermore, Increased PAQR7 expression was observed in breast tumor tissue compared to normal tissue (Left and middle panels, Fig. S1A), with statistical significance in the entire collection (right panel Fig. S1A), supporting previous observations of increased PAQR7 RNA expression in breast cancer (*14*). Overall, expression levels of all three CCM proteins are upregulated in breast tumors compared to normal tissues, suggesting a novel role for the CSC in tumorigenesis (*12*).

**Fig. 1.**
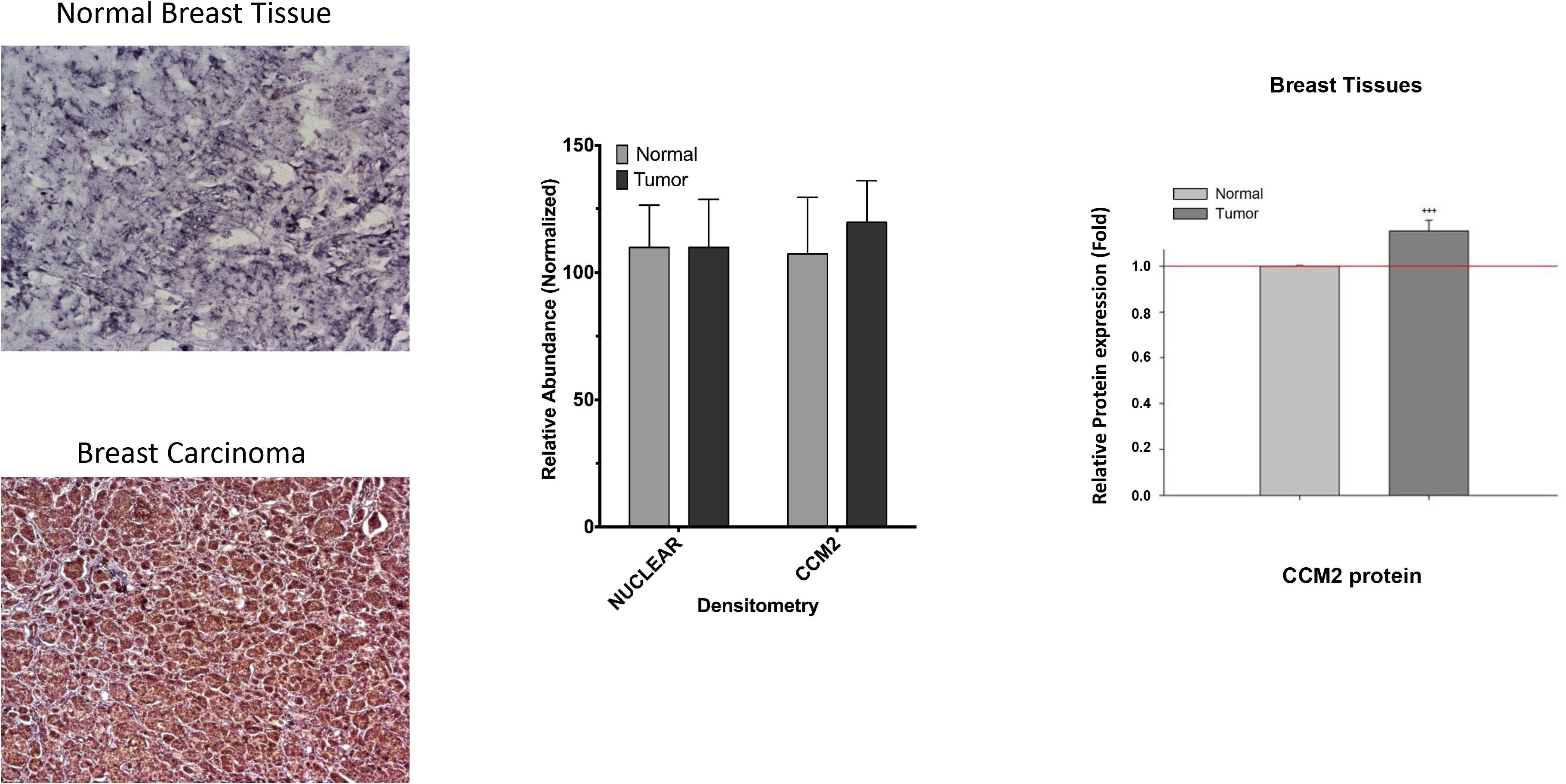
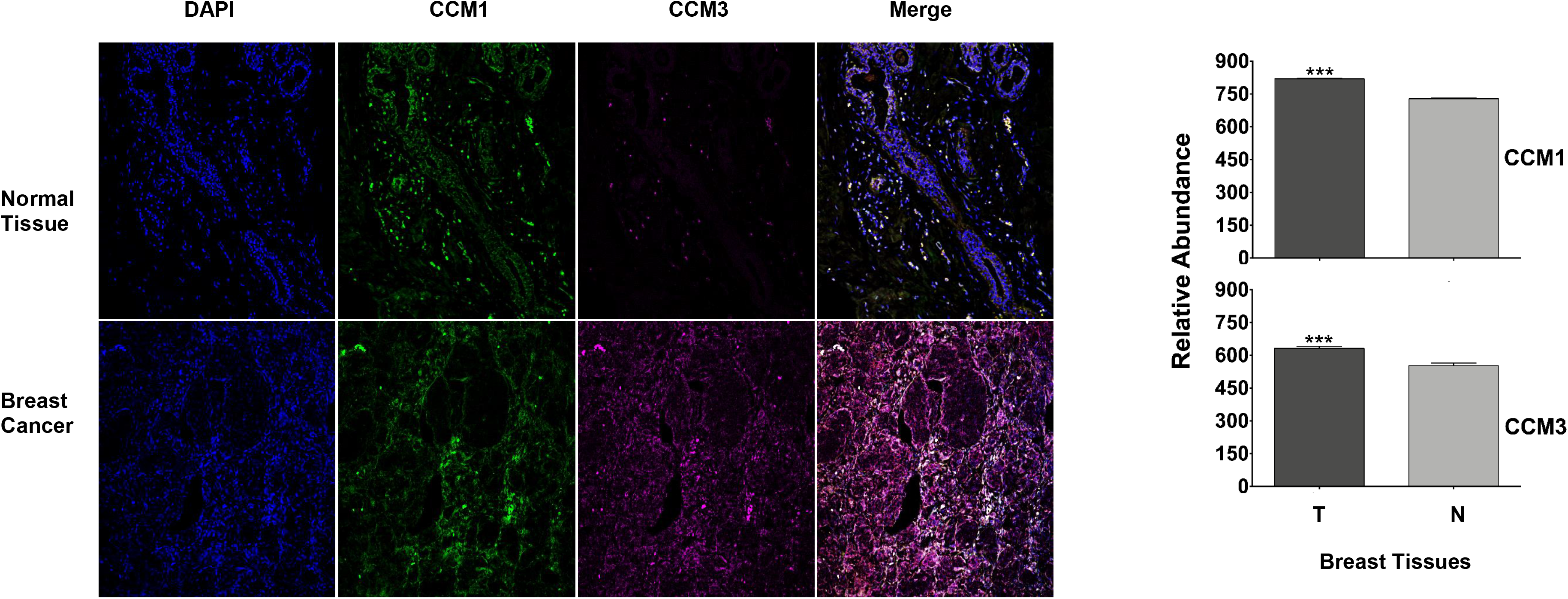
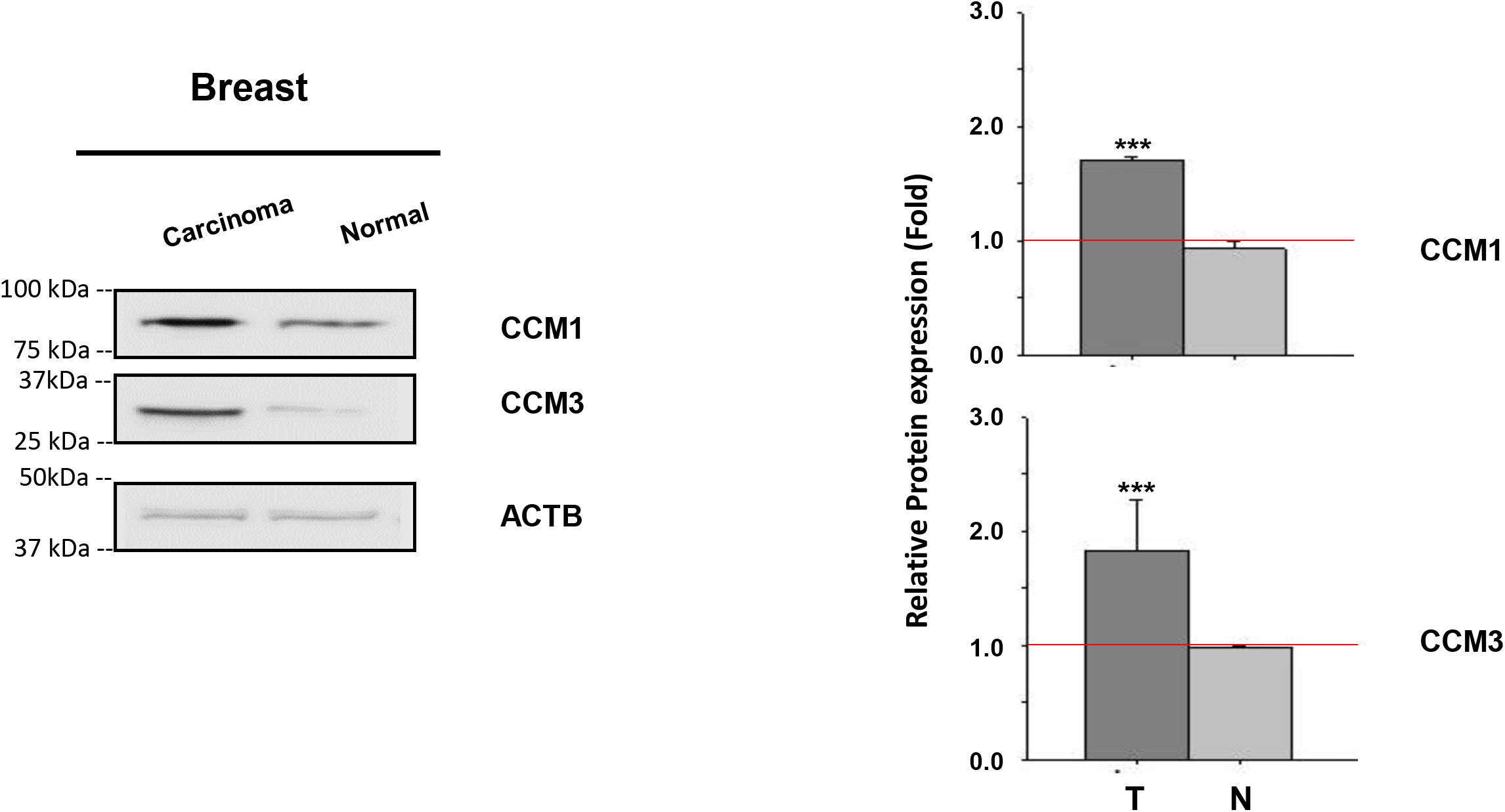
Up-regulation of CCM proteins in breast cancers. Relative expression of total CCM proteins in paired-tissue breast cancer samples **A**). IHC approaches utilizing HRP/DAB staining revealed an increase in the relative intensity of CCM2 staining between representative breast tumor tissue compared to normal breast tissues (left panel); Representative quantification is presented (ROI 2,398-8,417/section, middle panel). Statistically significant increase of CCM2 proteins were found from the entire collection of paired samples (n=11, ROI range=2,027-10,281/sample, right panel) **B**). Significant increased protein expressions of CCM1 and CCM3 proteins were visualized (left panel) in breast tumor (T), compared to normal tissue (N) samples using immunofluorescence imaging and CCM1 and CCM3 quantification displayed (ROI=159-2211, n=3). **C**. Significant increased protein expression of CCM1 and CCM3 proteins was found in paired breast protein lysates from tumor (T), compared to normal tissue (N) samples (n=4) (More detailed in Figure S1C).

#### Highly expressed CCM proteins in breast cancer cell lines

To elucidate mechanisms of CSC signaling involved in breast tumorigenesis, we screened candidate model cell lines and identified breast cancer cell line, T47D, as having higher endogenous expression, and a richer composition, of CCM2 isoforms (Fig. S1B). As a well-differentiated breast cancer cell line derived from an invasive ductal carcinoma, T47D is a luminal-A subtype of breast cancer cells (ER-positive, PR-positive, HER2-negative) (*15*). PR in T47D cells, with equally expressed isoforms (PR1/2), are estrogen insensitive (*16*), but markedly susceptible to progesterone (*17*), making it an ideal model for dissecting progesterone-responses in breast cancers (*15*).

### CSC coupling both classic and non-classic progesterone signaling in PR(+) T47D cells

#### Progesterone and mifepristone work synergistically, disrupting the CSC

To elucidate the underlying mechanisms, we systematically tested the effects of major female reproductive hormones; estrogen (EST), progesterone (PRG), and its antagonist, mifepristone (MIF), on the expression levels of three CCMs (1, 2, 3) proteins in T47D cells, using Western blots. Our data showed that PRG and its antagonist, MIF, but not EST, have significant negative impacts on the expression of all three CCM proteins (Fig. 2A). Surprisingly, we discovered that PRG works together with MIF, to further inhibit the protein expression of CCM1 and CCM3 (Fig. 2B), as well as CCM2 (Fig. 2C) in T47D cells, suggesting synergistic actions on disrupting the CSC. No synergistic actions on CCMs protein expression were observed between EST and PRG (Figure 2B), suggesting that this newly defined signaling pathway between PRG+MIF and the CSC likely bypasses classic PR1/2. Next, we found that PRG and MIF can suppress the protein expression of CCM1/3 in both time-dependent (Fig. 2D, left panel) and dose-dependent manners within the range of non-cytotoxic concentrations (*18*) (Fig. 2D, right panel), validating our previous observation that both PRG and MIF can independently inhibit the expressions of all three CCM proteins (Figs. 2A-C). Time-course data also indicated that this PRG/MIF-modulated inhibition of CCM proteins utilizes a rapid non-genomic (non-classic) mechanism (Fig. 2D, left panel). Real-time quantitative PCR (RT-qPCR) experiments showed that combined steroid (PRG+MIF) treatments can only inhibit RNA expression of CCM2 but has no effect on the RNA expression of either *CCM1/3* (Fig. 2E), suggesting that *CCM2* might be the direct target for steroid actions at the transcriptional level, and identifies the central role of CCM2 in this steroid action on the CSC. Down-regulation of PR1/2 has been observed at the transcriptional and translational levels with the treatment of PRG, and its antagonists, MIF, separately (*17*). Although MIF had long been observed as a “partial agonist” to PRG actions with unknown mechanisms (*19*), this “agonist” role of MIF was thought only to occur in the absence of PRG (*19*). Our data represent the first demonstration that either PRG or MIF can work independently, and synergistically to decrease the expression levels of CCMs proteins, in a non-classic mechanism, at both the transcriptional and translational levels in T47D cells.

**Fig. 2.**
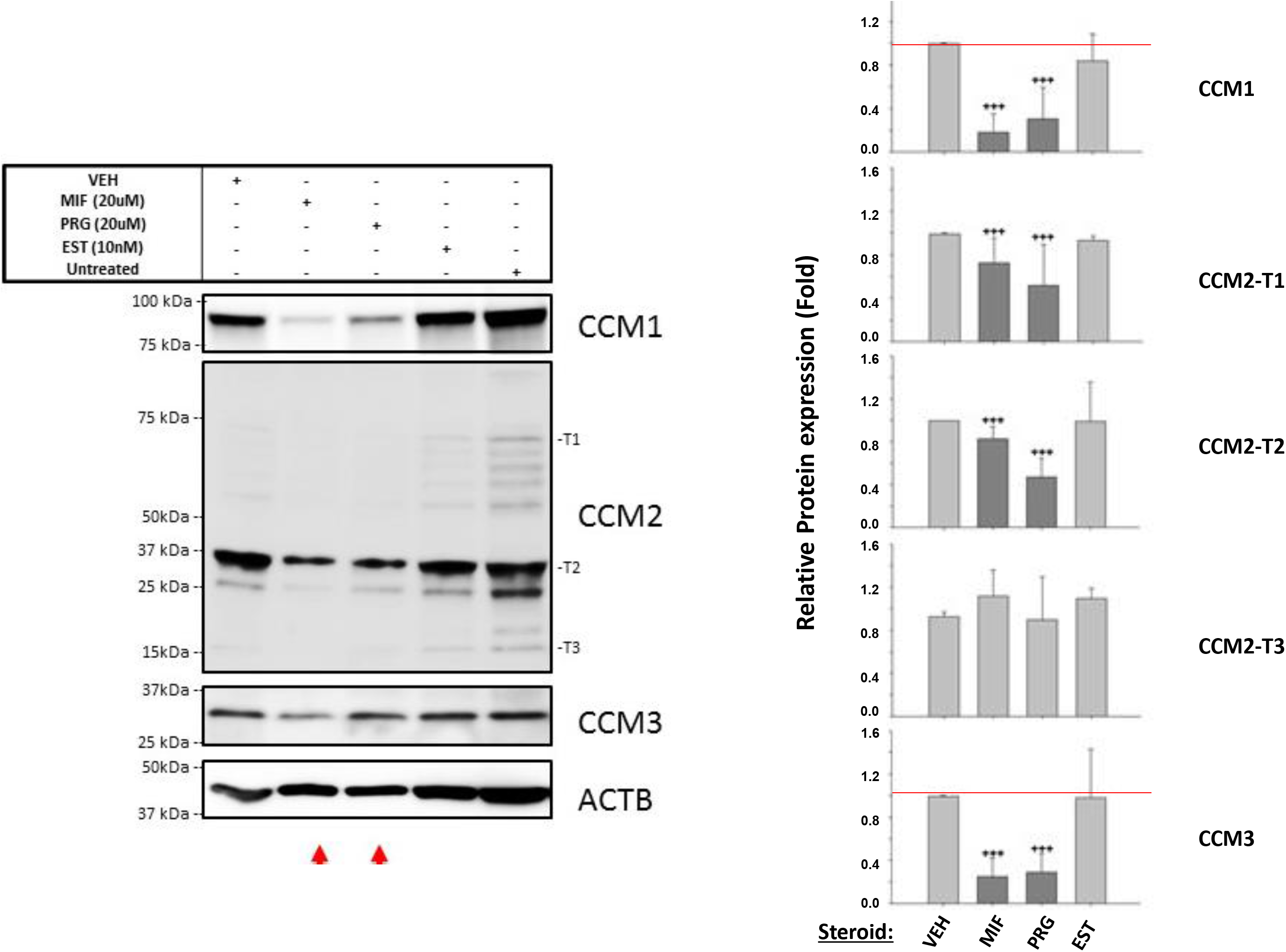
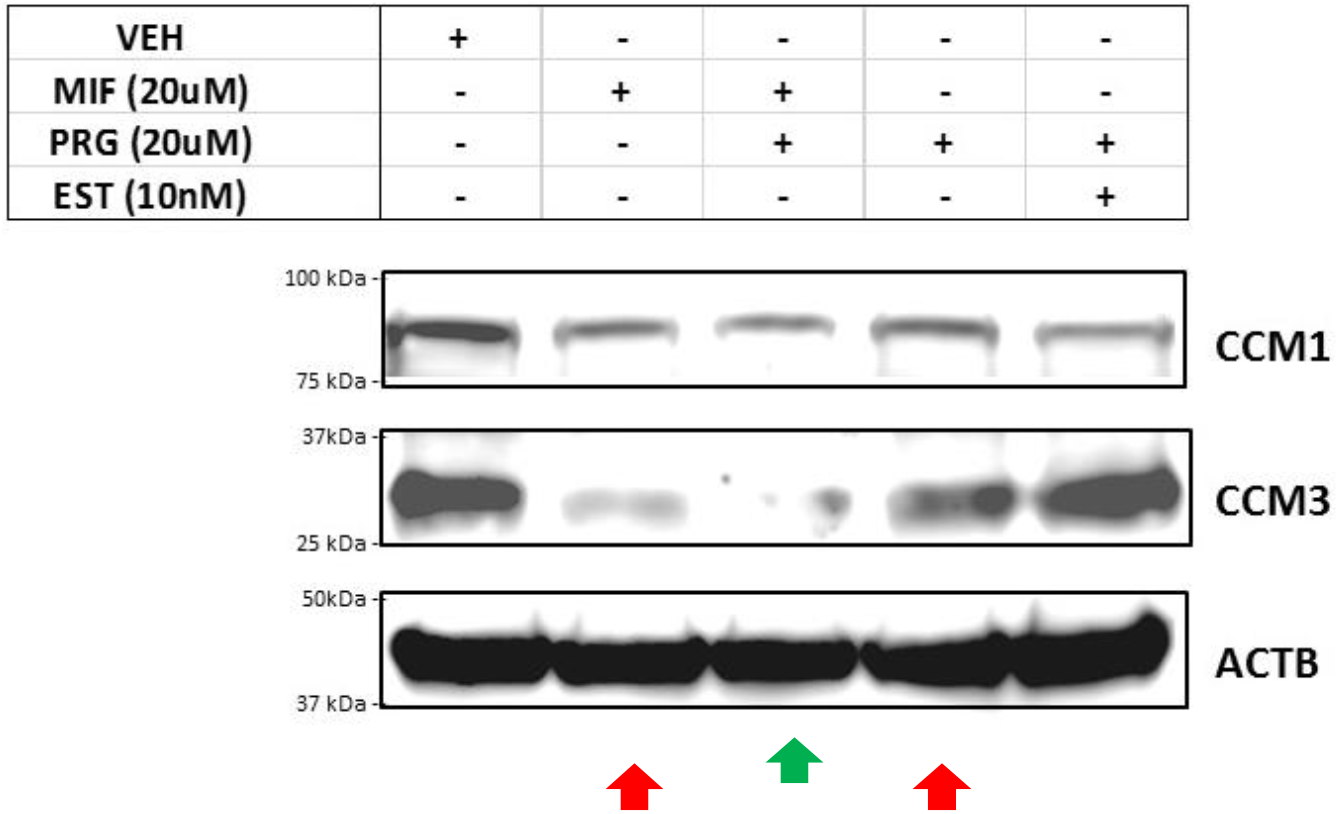
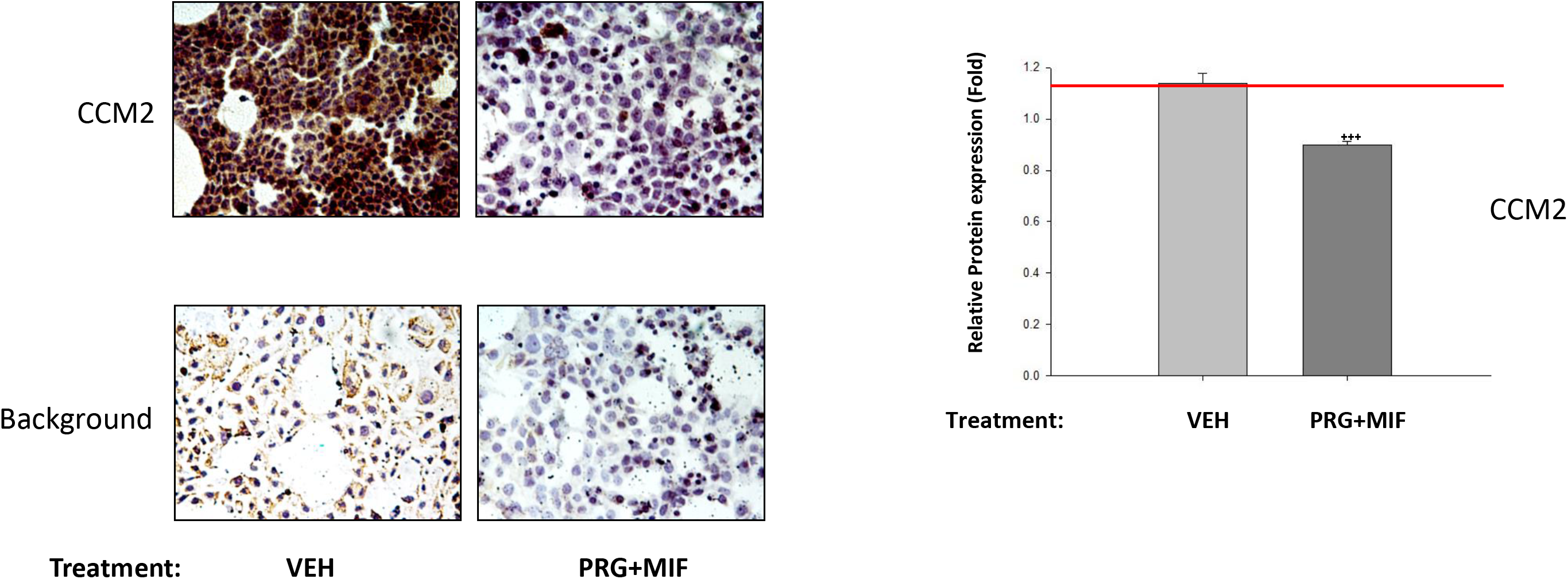
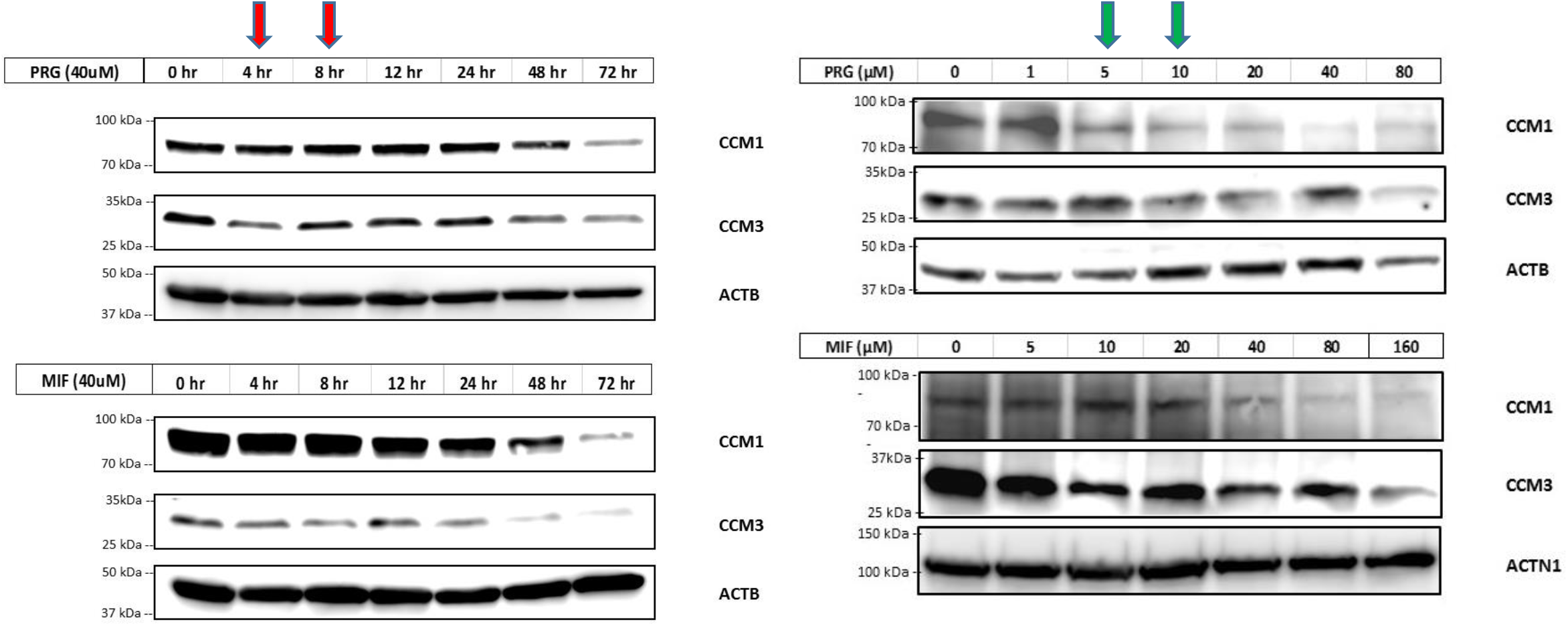
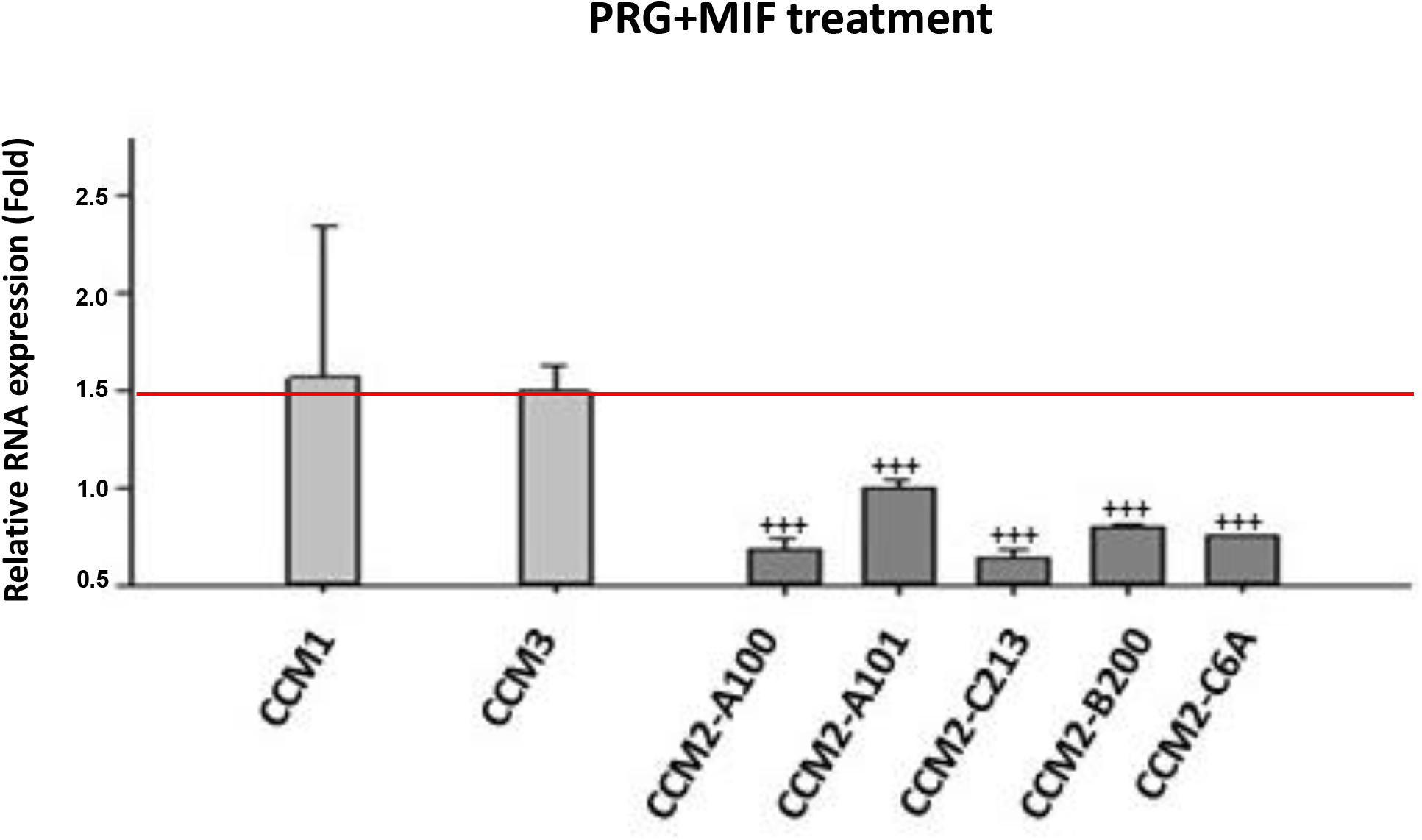
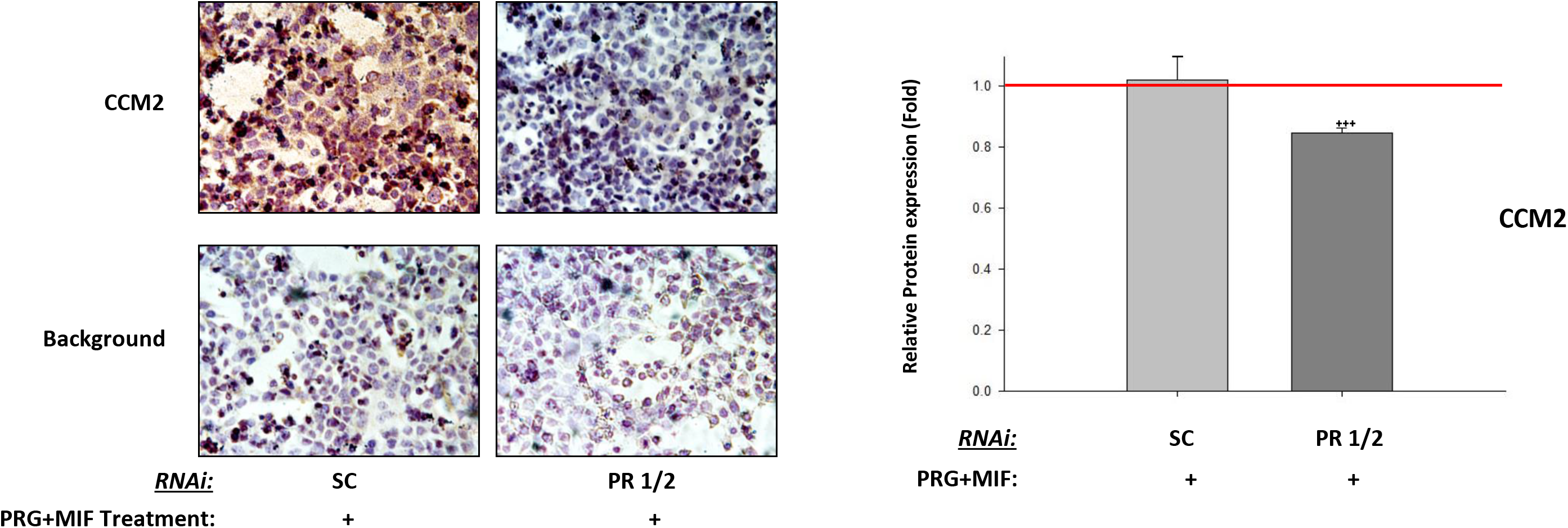
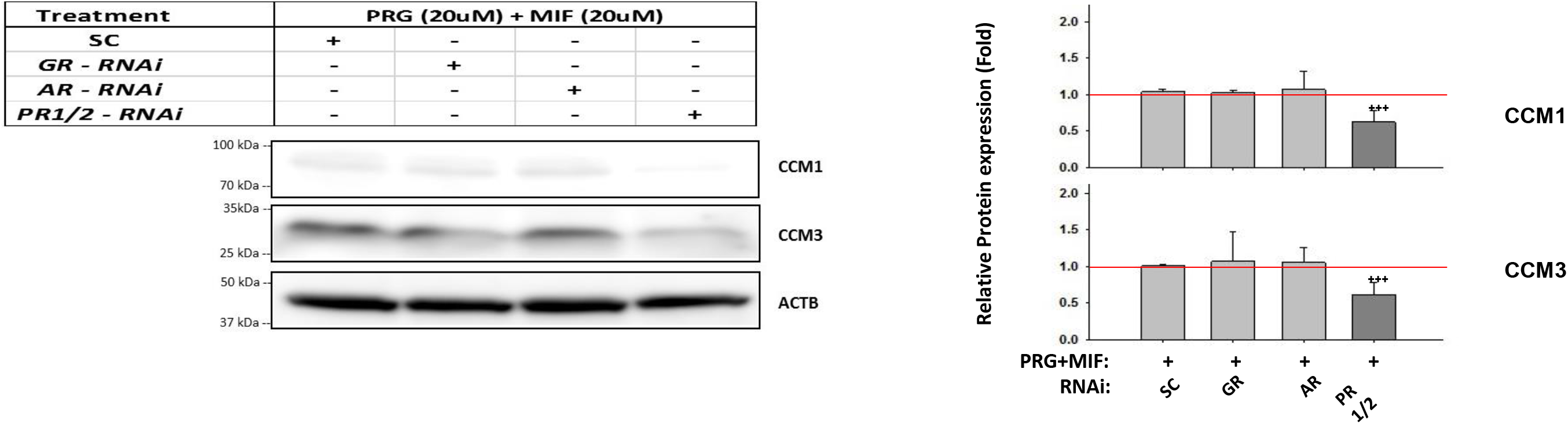
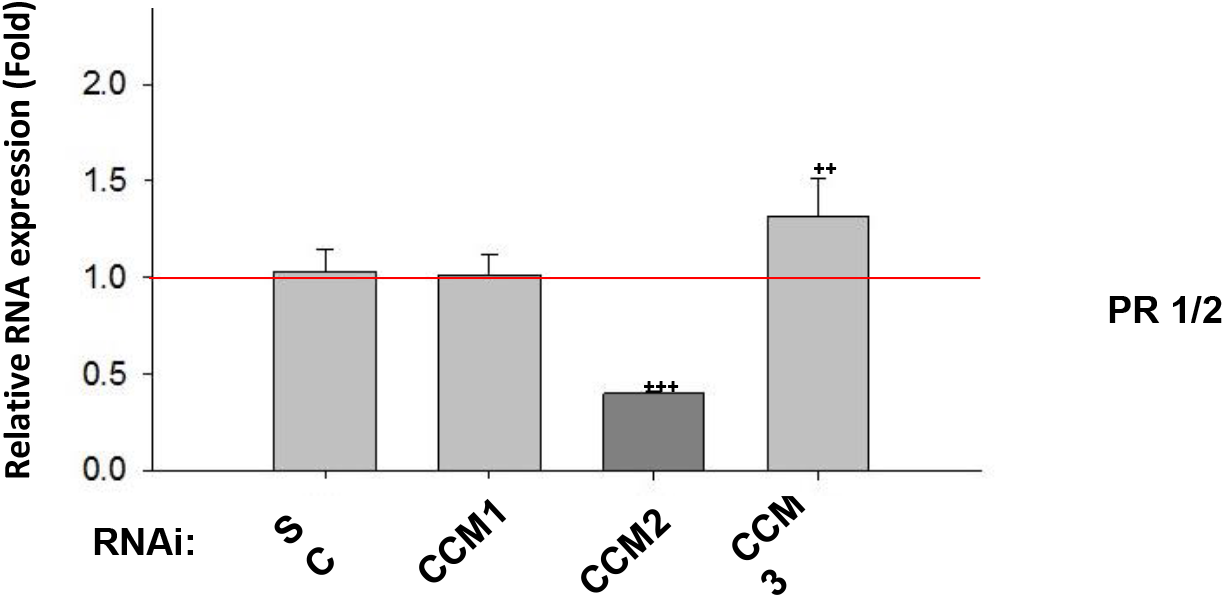
Expression levels of CCM proteins in PR-positive [PR(+)] breast cancer cells T47D are modulated by both progesterone and mifepristone. **A.** Expression levels of CCM1, CCM2, and CCM3 in T47D cells can be suppressed independently with either progesterone (PRG) or mifepristone (MIF). Cells were treated with vehicle control (ethanol/DMSO, VEH), mifepristone (MIF, 20 μM), progesterone (PRG, 20 μM), estrogen (EST, 10 nM), or media only (Untreated) respectively. While estrogen (EST) displayed little effect on the expression levels of CCM proteins, both PRG and its antagonist MIF can independently suppress protein expression of all three CCMs (columns indicated by red arrow, left panel). Relative protein expression levels of three selected CCM2 isoforms (T1, T2, T3), CCM1, and CCM3 was performed through quantification of band intensities of CCM proteins normalized against β-actin (ACTB) followed by control vehicle treatment (red line, right panel, n=5). **B.** Progesterone (PRG) and mifepristone (MIF) can work synergistically to enhance their inhibitory roles in protein expression of CCMs. Progesterone (PRG, 20 μM) and mifepristone (MIF, 20 μM) independently inhibiting expression of CCM1 and CCM3, were further confirmed (red arrows), and the combination of both progesterone and mifepristone (PRG+MIF, 20 μM each) synergistically enhance their inhibitory effects on expression levels of CCM1 and CCM3 (green arrow) while no similar effect was seen in estrogen combined with PRG (EST, 10 nM) or vehicle controls (ethanol/DMSO, VEH). **C.** The decreased expression levels of CCM2 proteins in T47D cells with combined progesterone/mifepristone treatment. CCM2 protein expression in T47D cells with combined steroid treatment (PRG+MIF, 20 μM each) or vehicle control (VEH, ethanol/DMSO) were further measured with CCM2 antibodies utilizing immunohistochemistry (IHC) applications with HRP/DAB detection system (left panel). CCM2 quantification demonstrated significant decrease of CCM2 proteins (right panel, ~10,000 ROI/section) normalized to background staining (red line, n=4). **D.** Both progesterone and mifepristone can independently induce the decreased protein expression of CCM1 and CCM3 in both time and dose-dependent manners. T47D cells were treated with progesterone (40 μM, PRG) and mifepristone (40 μM, MIF) for the times indicated in the left panel or treated for 72 hrs with a series of concentrations (0-160 μM) of progesterone (PRG) or mifepristone (MIF) (right panel). Decreased protein expression of CCM1 and CCM3 can be observed as early as 4 hr post-treatment, indicating the non-classic action of both PRG or MIF on the protein expression of CCM1 and CCM3 (red arrows). The newly identified non-classic action of both PRG or MIF on protein expression of CCM1 and CCM3 can be observed at very low doses (5-10 μM for PRG and 10-20 μM for MIF, green arrows), indicating the physiological sensitivity and relevance of PRG and its antagonist, MIF in CSC-mediated signaling. **E.** Only mRNA expression of CCM2 isoforms is suppressed by the PRG+MIF treatment. T47D cells with PRG+MIF treatment (20 μM each) show decreased mRNA expression of all CCM2 isoforms but not that of either CCM1 or CCM3. Relative RNA expression levels were measured through RT-qPCR (triplicates per experiment, n=3). **F.** Silencing of total progesterone receptors (both isoform A and B-termed PR1/2) further enhances the suppression of CCM2 protein expression in T47D cells under steroid action. Significantly decreased expression levels of CCM2 proteins in T47D cells were observed utilizing HRP/DAB staining (left panel) after treatment with siRNA-PR 1/2 or scrambled control (SC) for 24 hrs, then treated with progesterone/mifepristone (PRG+MIF, 20 μM each) for another 48 hrs (left panel). Quantification of CCM2 in T47D cells is displayed (right panel, ~10,000 ROI/section) normalized to background staining (red line, n=4). **G**. Both CCM1 and CCM3 proteins are also sensitized to MIF+PRG treatment. T47D cells were treated with RNAi-KD for progesterone receptors (PR1/2), androgen receptor (AR) or glucocorticoid receptor (GR) for 24 hrs, followed by PRG+MIF treatment (20 μM each) for 48 hrs. Significantly decreased protein levels of CCM1 and CCM3 were observed in PR1/2-KD T47D cells (Left and right panels, n=4). **H**. RNA expression level of total progesterone receptors (PR1/2) is influenced by the CSC in T47D cells. After silencing CCMs (1, 2, or 3) for 48 hrs, significant decreased RNA expression levels of progesterone receptors (PR1/2) in CCM2-KD was observed, while significant increased RNA expression levels of PR1/2 in CCM3-KD was observed. (More detailed in Figure S2).

#### PRG+MIF synergistic actions stabilize the CSC through classic progesterone receptors (PR1/2)

To investigate the relationship between PR1/2 and CCM2 under steroid action, PR1/2 genes were silenced by RNAi, followed by PRG+MIF treatment in T47D cells, resulting in significantly decreased expression of CCM2 proteins in PR-silenced cells (Fig. 2F), compared to scrambled control (SC), suggesting the protective role of PR1/2 on the stability of CCM2 under steroid action. MIF has been reported to bind to all three major steroid receptors: PR(1/2), GR (glucocorticoid receptor) and AR (androgen receptors), and commonly acts as an antagonist for all three receptors (*20, 21*). The degree of PR, GR, and AR inhibition by MIF is variable, depending on specific cell types (*22*). Both ARs and PRs are highly expressed, but the expression of GRs is very low and barely detectable in T47D cells. Using RNAi, we silenced these steroid hormone receptors (AR, GR, PR1/2) followed with PRG+MIF treatment. PR1/2-silenced T47D cells showed significantly enhanced inhibitory effects of PRG+MIF treatment on the expression of CCM1/3 proteins, compared to SC control (Fig. 2G), supporting the protective role of PR1/2 on the stability of the CSC under steroid actions. Both AR and GR play no role in the negative effects of the steroid action on the stability of the CSC (Fig. 2G). Knocking down CCM1, CCM2, and CCM3 independently revealed significantly decreased RNA expression levels of PR1/2 by silencing CCM2 (Fig. 2H), implying that PR1/2 and CCM2 reciprocally influence their expression levels at both transcriptional and post-translational levels. The novel synergistic actions of PRG and MIF on the CSC are quite surprising since MIF binds to PR1/2 with greater affinity than PRG, forming PR1/2-MIF complexes, inhibiting PR1/2- PRG-associated actions. MIF has been widely used as an antiprogestin for contraception and termination of early pregnancies (*23*). However, as a type-II antiprogestin, MIF can also act as an agonist in supporting PRG-associated actions in a cell-specific manner with unknown etiology (*24*).

#### PRG and MIF work synergistically to influence expression of both PR1/2 and membrane progesterone receptors (mPRs) through the CSC in T47D cells

Decreased endogenous CCM1/3 proteins were initially observed at 4 hrs after PRG+MIF treatment (Fig. 2D), suggesting the involvement of rapid non-classical PRG actions, probably through newly identified mPRs. Non-genomic effects of PRG mediated by mPRs have been reported to induce rapid intracellular changes (*4*). Further, a potential cross-talk mechanism between PR and mPRs has been suggested in mediating progesterone actions (*25*), but without any supporting data, yet. In one recent report, mPRα and β transactivates PR-2 in PR-positive [PR(+)] myometrial cells (*26*), suggesting potential cross-talk between classic and non-classic PRG receptors (*26*). Our data also correlated expression between CCM proteins (Figs. 1A, 1B) and PAQR7 (Fig. S1A) in breast tumors (*12*). RT-qPCR analysis revealed significantly decreased RNA expression levels of PR1/2 upon steroid treatment in T47D cells, in line with previous report (*27*). RNA expression levels among the majority of mPRs, PAQR6, PAQR7, and PAQR8 were significantly decreased, while increased expression of PAQR5 and PAQR9 were observed; no change was found for PGRMC1 (Fig 3A). Overall, mRNA levels of PR1/2 and most mPRs were remarkably diminished in combined steroid-treated T47D cells, suggesting inhibition of PR1/2 and mPRs by steroid actions at the transcriptional levels (Fig. 3A). Surprisingly, western blot data demonstrated significantly decreased expression levels of all PAQRs proteins, while no significant changes of protein expression for PGRMC1 under MIF+PRG treatment (Fig 3B), indicating that the overall expression of all PAQRs can be modulated by steroid action at both the transcriptional and translational levels.

**Fig. 3.**
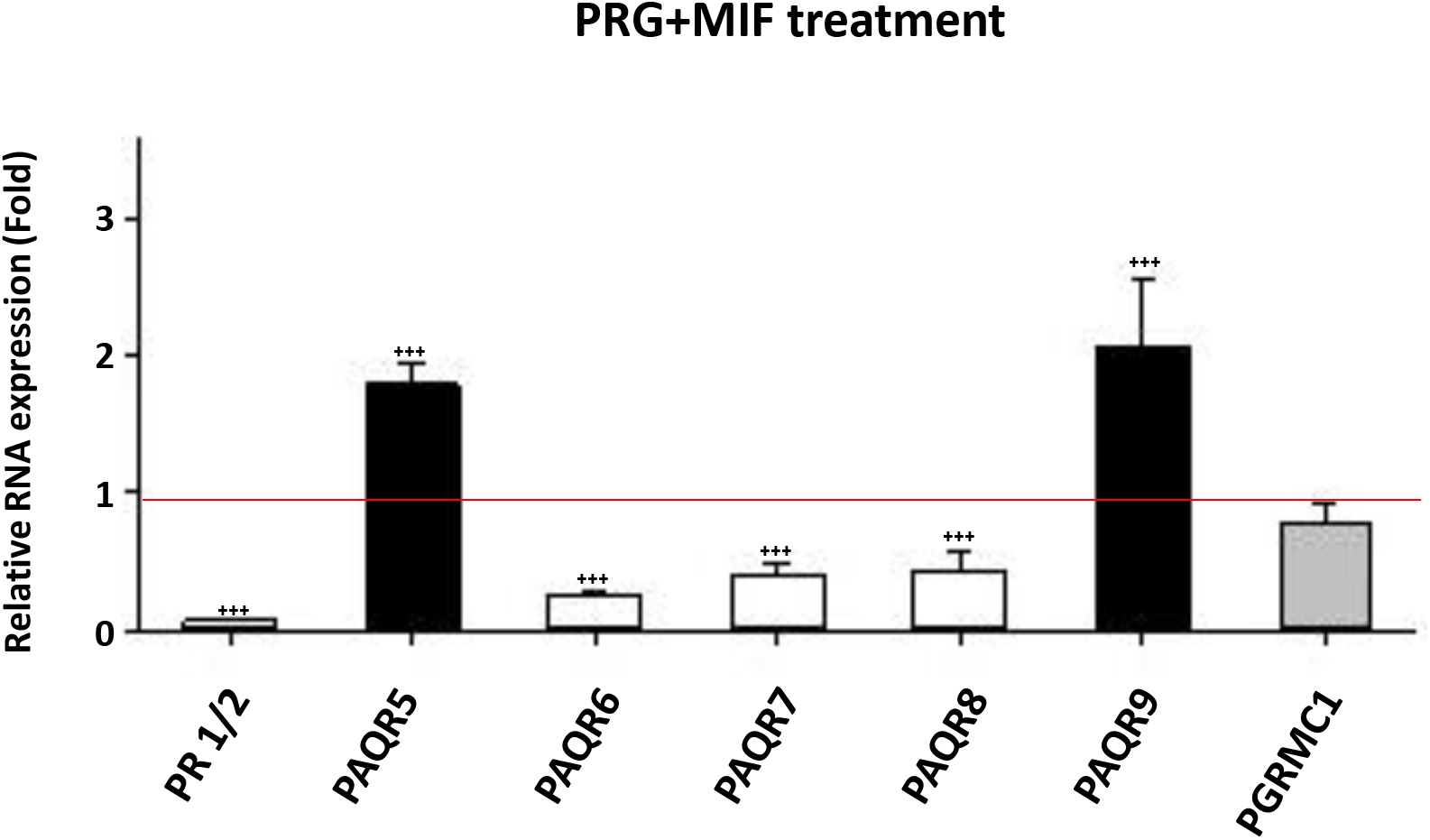
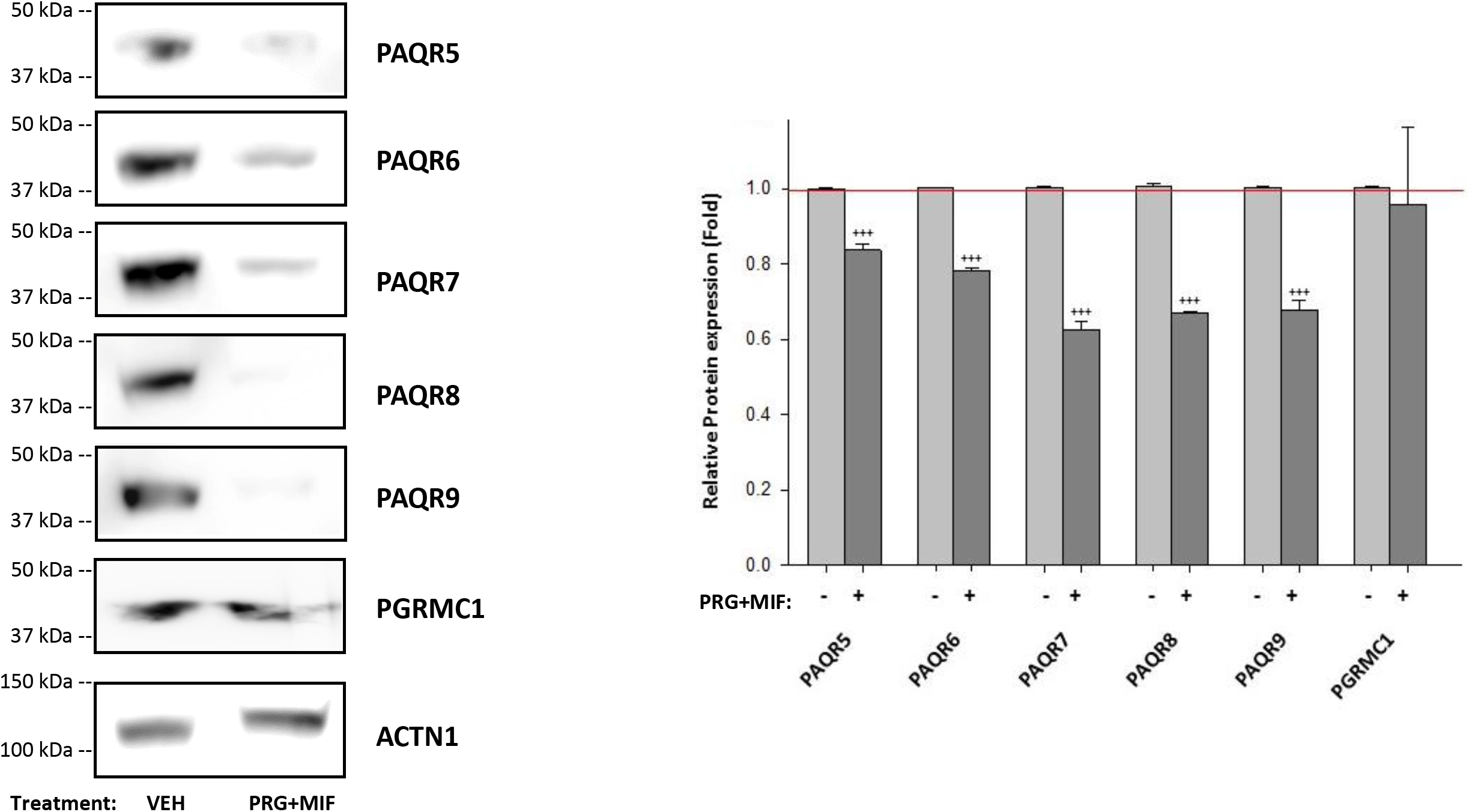
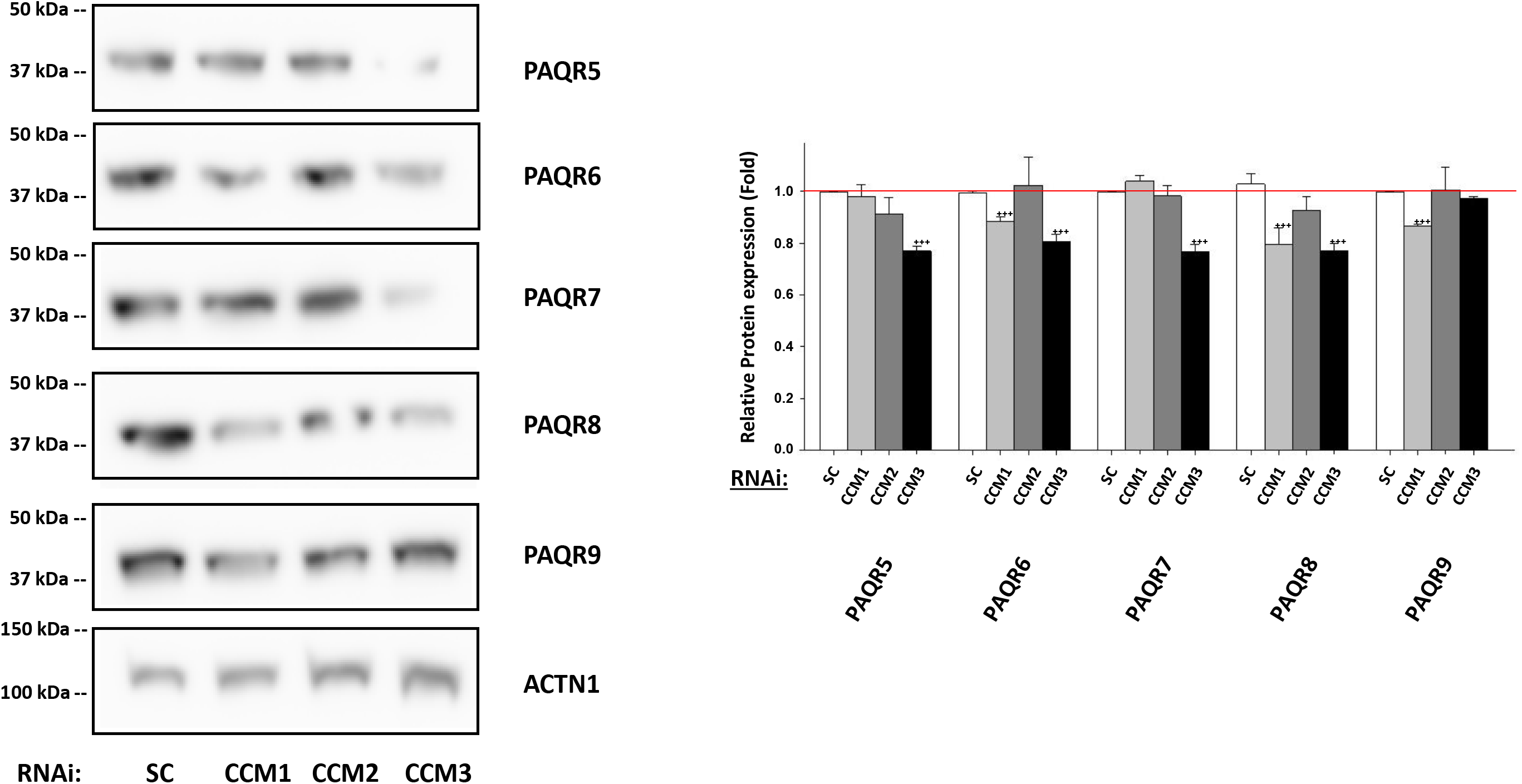
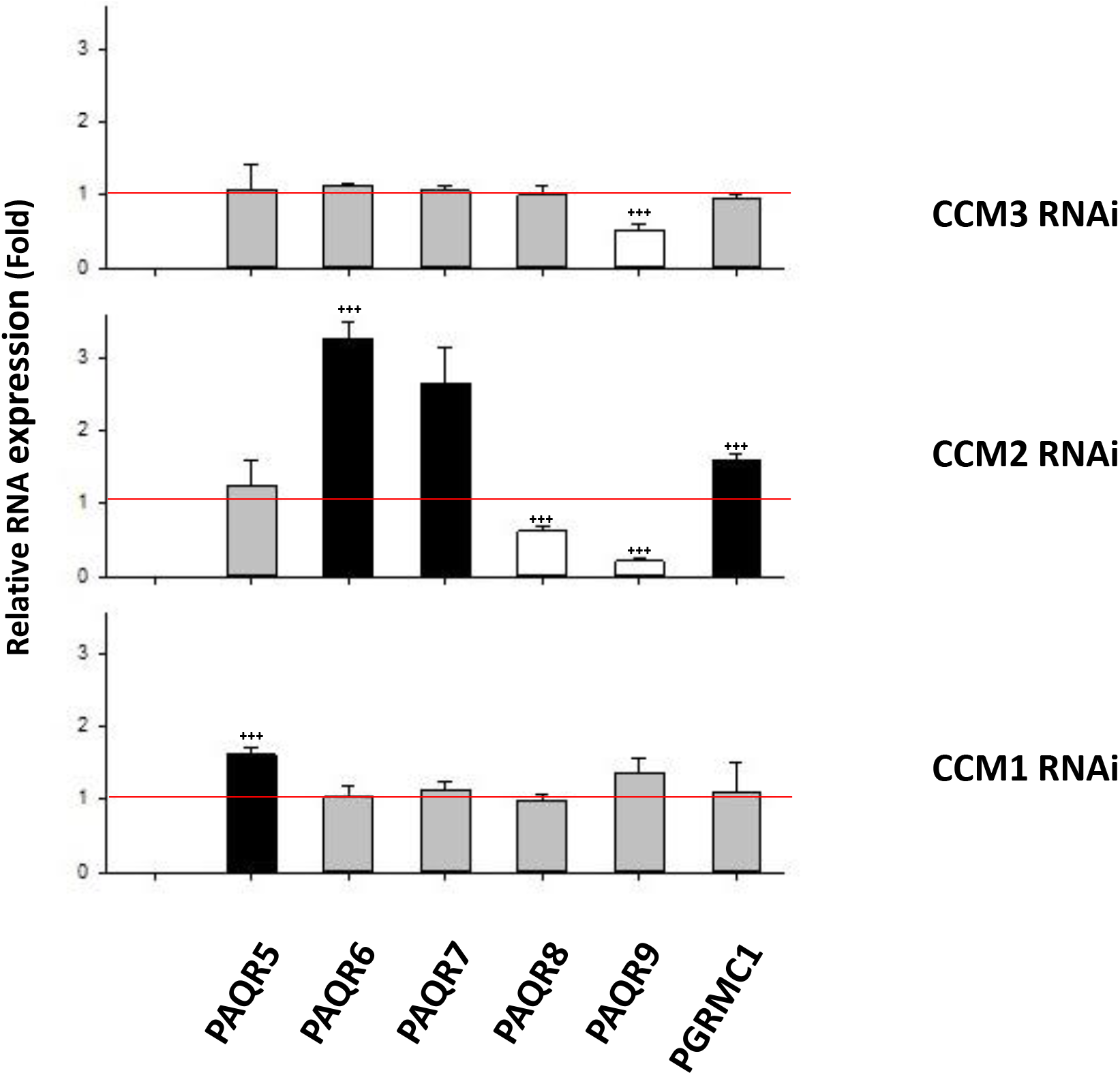
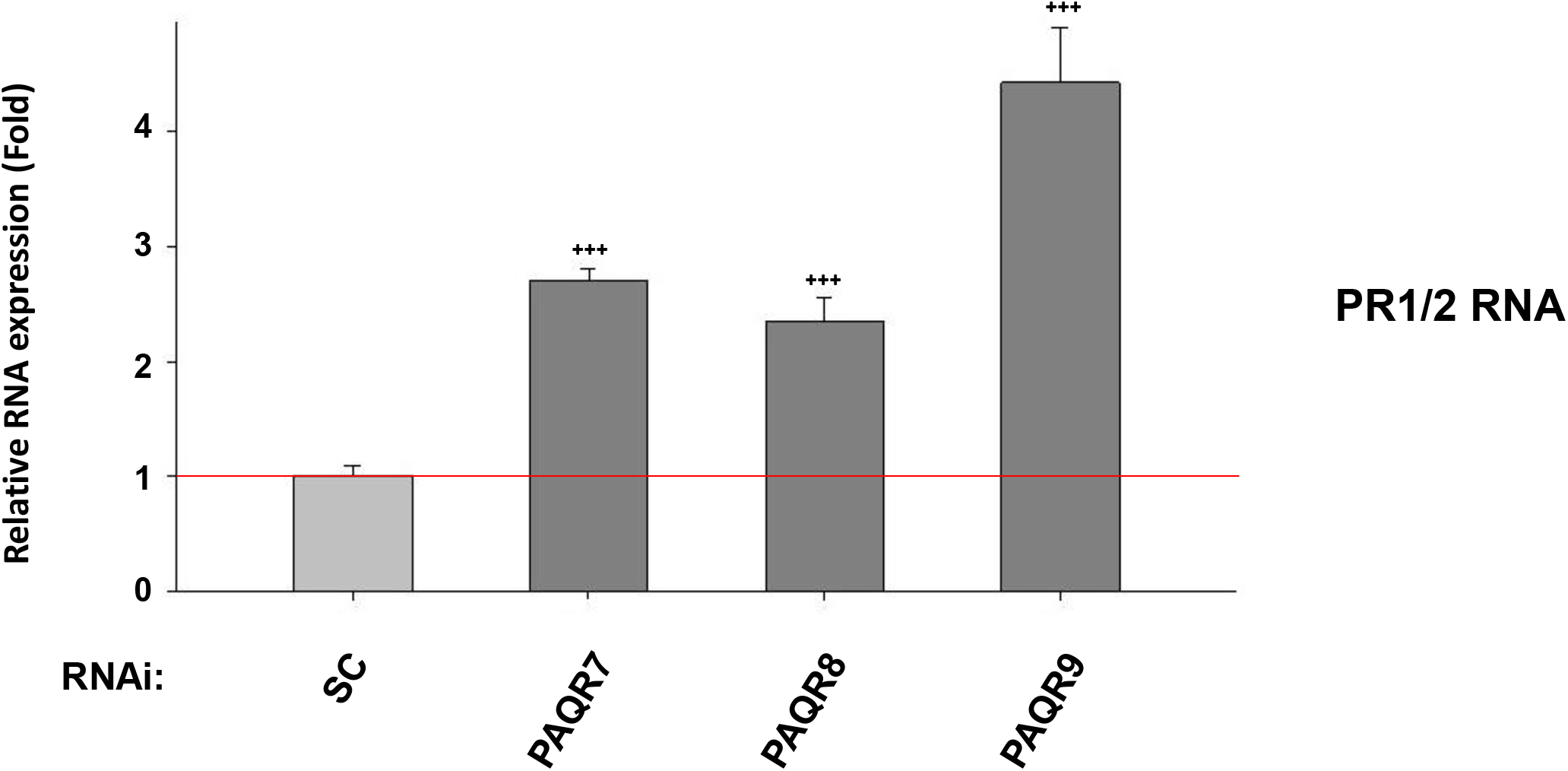
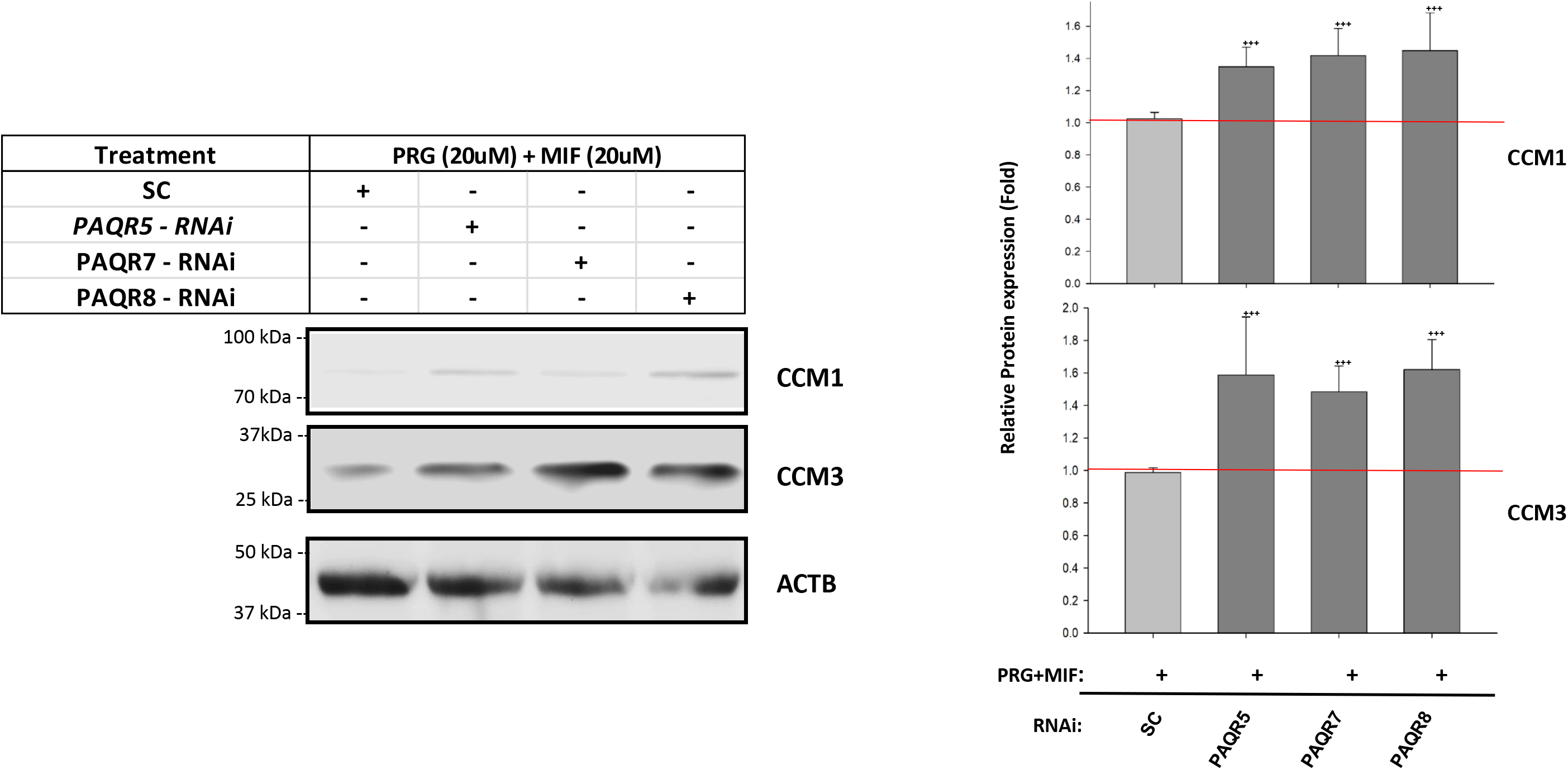
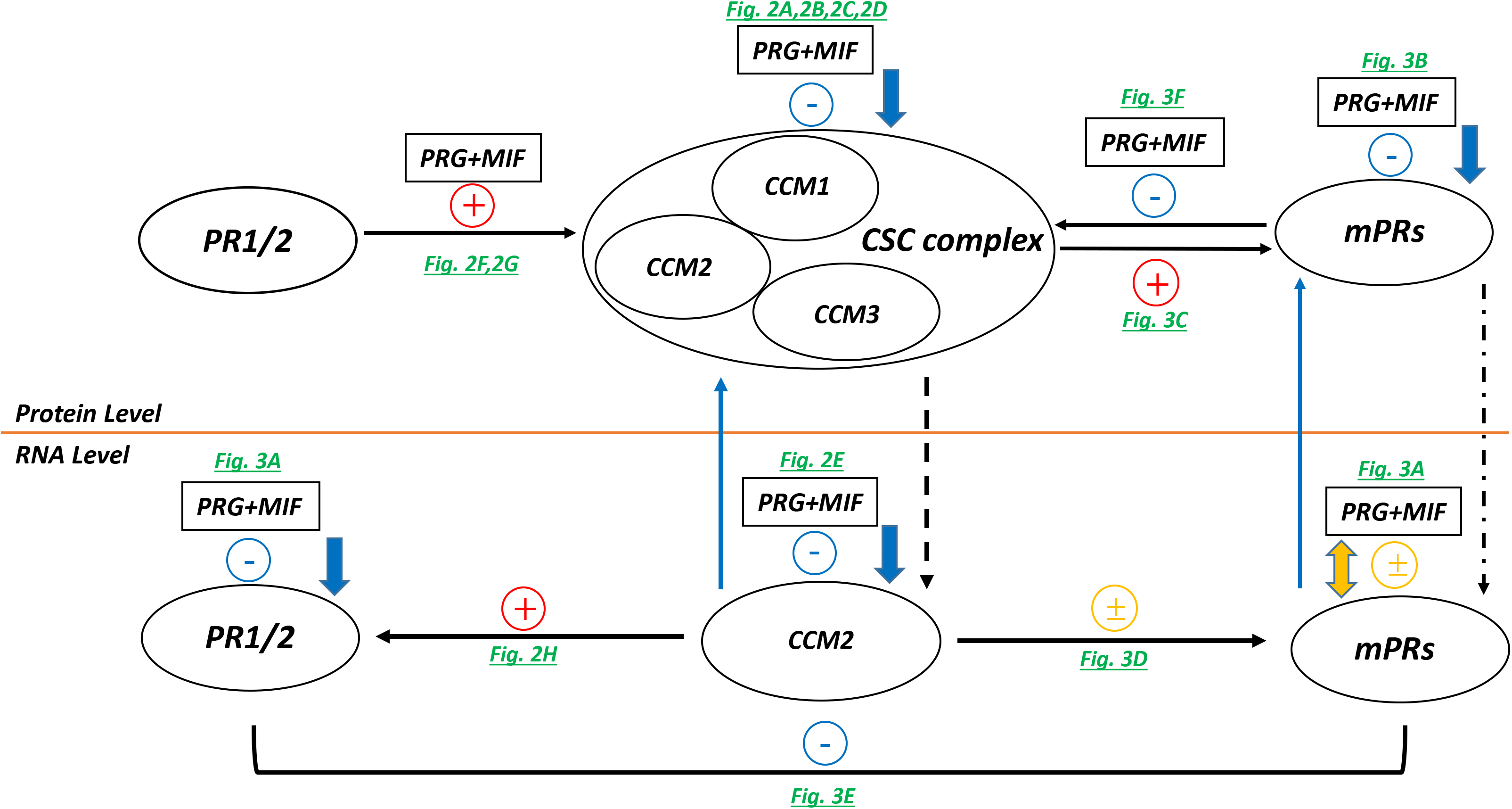
Expression levels of progesterone receptors (PR1/2) and membrane progesterone receptors (mPRs/PAQRs) are modulated by progesterone and mifepristone and the CSC in PR positive T47D cells. **A**. RNA expression levels of progesterone receptors (PR1/2) and membrane progesterone receptors [mPRα (PAQR7), mPRβ (PAQR8), mPRγ (PAQR5), mPRδ (PAQR6), mPR∊ (PAQR9) and PGRMC1] are influenced by steroid action. After PRG+MIF treatment (20 μM each) for 48 hrs, RNA expression levels of total progesterone receptors (PR1/2) and select mPRs (PAQR 6, 7, 8) were significantly decreased, while the RNA expression level of two membrane progesterone receptors (PAQR 5, 9) were increased and no changes were observed in PGRMC1. Relative RNA expression changes were measured by qPCR (Fold changes) and normalized to scramble control (red line, triplicates per experiment, n=3). **B**. Protein expression levels of all mPRs (PAQRs, PGRMC1) are suppressed by steroid action. After PRG+MIF treatment (20 μM each) for 48 hrs, Western blots demonstrated dramatic decreased expression levels of all mPR proteins [PAQR 5 (mPRγ), PAQR6 (mPRδ), PAQR7 (mPRα), PAQR8 (mPRβ), PAQR9 (mPR∊)], while no apparent change in PGRMC1 was observed, compared to vehicle control (VEH) (Left and right panels). Relative expression levels of mPR proteins were measured through quantification of band intensities and normalized against α-actinin (ACTN1) and vehicle control (VEH) (right panel, n=3). **C.** Protein expression levels of mPRs (PAQRs) are mainly suppressed by silencing *CCM1* and *CCM3* genes. After silencing all three CCMs (CCM1, CCM2 or CCM3) for 48 hrs, significantly decreased expression levels of most mPRs (PAQR 5, 6, 7, 8, 9) were demonstrated by silencing either CCM1 or CCM3, while no statistically significant changes was observed by silencing CCM2 (Left and right panels). Relative protein expression levels of mPRs (PAQR 5, 6, 7, 8, 9) were measured through quantification of band intensities and normalized against α-actinin (ACTN1) followed by scramble control (SC) treatment (right panel, red line, n=3). **D**. RNA expression levels of all membrane progesterone receptors (PAQRs, PGRMC1) are mainly modulated by CCM2. After silencing all three CCMs (1, 2 or 3) for 48 hrs, RNA expression levels of all mPRs (PAQRs, PGRMC1) were measured by qPCR (Fold changes). The majority of altered RNA expression of mPRs were only observed in silencing CCM2 (triplicates per experiment, n=3). **E**. RNA expression levels of total progesterone receptors (PR1/2) is significantly increased by silencing mPRs (PAQRs) genes. After silencing mPRs (PAQR 7, 8, 9) genes for 48 hrs, significant increased RNA expression levels of progesterone receptors (PR1/2) were observed. The relative RNA expression changes of progesterone receptors (PR1/2) in T47D cells were measured by qPCR (Fold changes) (triplicate per experiment, n=3). **F.** Both CCM1 and CCM3 proteins are stabilized to steroid action after silencing mPRs in T47D cells. Increased protein levels of CCM1 and CCM3 comparing to SC controls was observed when cells were treated with RNAi-knockdown (KD) for major mPRs (PAQR5, 7, 8) for 24 hrs, followed by PRG+MIF treatment (20 μM each) for an additional 48 hrs (Left panel). Relative expression levels of CCM1 and 3 proteins were measured through quantification of band intensities normalized against β-actin (ACTB) followed by SC controls, and represented with bar plots (red line right panel)(right panel, n=4). **G**. The summarized feedback regulatory networks among the CSC, PR (1/2), and mPRs signaling complex under steroid action for PR+ breast cancer cell T47D homeostasis. Yellow line separates transcriptional and translational levels. The + symbols represent enhancement, - symbols represent inhibition while ± symbol represent various regulation for the expression of targeted genes/proteins. Red colored symbols/lines represent positive effects of PRG+MIF treatment, blue colored symbols/lines represent negative effects of treatment, while orange colored symbols/lines represent variable effects. Dark green colored letters indicate the direct supporting data generated from this experiment. Arrow indicates effect direction, solid line is the direct impact, dotted line for indirect effects (More detailed in Figure S3C).

#### The expression of mPRs is modulated by the CSC while the expression of PR1/2 is modulated by mPRs in PR(+) T47D cells

CCM1, CCM2, and CCM3 genes were silenced, respectively, and protein expression levels of PAQRs were examined. Significantly decreased expression of PAQR proteins was observed mostly in either CCM1-Knockdown (KD) or CCM3-KD (Fig. 3C). Interestingly, increased RNA expression levels of PAQR5 in CCM1-KD and decreased RNA expression levels of PAQR9 in CCM3-KD were observed, respectively. RNA expression patterns of PAQRs were mostly altered in CCM2-KD, with increased RNA expression levels of PAQR6, PAQR7, and PGRMC1 while decreased RNA expression levels of PAQR8 and PAQR9 were observed (Fig. 3D). Thus, our data suggest that CCM2 might be the key player influencing the expression levels of all mPRs at both the transcriptional and post-translational levels, while CCM1/3 proteins only influence the expression levels of mPRs at the translational level (*21, 28*).

To further delineate the cellular relationship among PR1/2, mPRs/PAQRs, AR, and GR, we next knocked down AR, GR, and PR1/2 and discovered no change of mPRs/PAQRs expression either at the protein (Fig. S3A) or RNA level (Fig. S3B), indicating PR1/2, AR, and GR do not regulate the expression of mPRs/PAQRs either at the protein or RNA level. However, when silencing mPRs/PAQRs, significantly increased RNA expression of PR1/2 was observed (Figs. 3E), suggesting that PAQRs might negatively influence PR1/2 expression at the transcription level.

#### PRG+MIF actions destabilize the CSC through mPRs

To further delineate the relationship between CCMs and mPRs/PAQRs, we silenced three major mPRs (PAQR5, 7, 8) under PRG+MIF treatment; mPR-silenced T47D cells significantly reduced the inhibitory effect of steroid actions on the expression of CCM1/3 proteins, compared to SC (Fig. 3F), supporting the notion that mPRs destabilize the CSC under steroid actions, opposite to PR1/2 (Fig. 2G).

#### mPRs and PR1/2 signaling cascades are coupled through the CSC in PR(+) T47D cells

The novel finding of balanced modulation of the CSC stability through either positive effects of PR1/2 or negative effects of mPRs/PAQRs, are quite exciting, since it not only emphasizes the importance of the existence and balance of both, classic and non-classic PRG receptors on CSC functions, but also identified the unique role of the CSC in the cross-talk between two types of PRG receptors in PR(+) cells. Our notion is further supported by a previous observation that PRG can act simultaneously on both nuclear and membrane PRG receptors, and activation of mPR signaling can potentiate hormone activated nuclear PR-2 isoform (*26*). This intricate feedback regulatory network among the CSC-PR1/2-mPRs pathways, under steroid actions in PR(+)T47D cells, can be described with an integrated model of the CSC modulating PRG signaling on both classic and non-classic PRG receptors. In this model, steroid hormone-dependent actions are realized through balanced efforts between PR1/2 and mPRs, and is further fine-tuned by the CSC. This model is summarized in a schematic representation, where key feedback regulatory relationships at both transcriptional and translational levels are illustrated with respective supporting data (Figs. 3G).

### CCM2 is the cornerstone for the stability of the CSC complex

In the CSC complex, CCM1 can stabilize ICAP1α and CCM2 through its interactions with both (*29–31*). Then it was found that CCM1/CCM2 can enhance proteins stability reciprocally, and this reciprocal relations was further extended to all CCMs proteins (*8*), leading us to re-examine the relationship among three CCMs proteins.

Surprisingly, silencing of CCM2 resulted in significantly decreased expression of CCM1/CCM3 proteins, while silencing CCM1/CCM3 did not alter expression of CCM2 protein (Fig. 4A). RT-qPCR however, demonstrated significantly increased RNA expression level of CCM2 isoforms with silencing either CCM1 or CCM3 while no change of expression of CCM1/CCM3 RNAs was observed with silencing CCM2 (Fig. 4B), suggesting that there is a feedback regulatory loop to influence the expression of CCM2, based on cellular levels of CCM1 and CCM3 at the translational level. Identical results were obtained from 293T cell (Fig. S4A1-2), human brain microvascular endothelial cell (HBMVECs) (Fig. S4B1-2) and zebrafish Ccm1/Ccm2 mutant strains (Fig. S4C1-2), indicating that CCM2 is the cornerstone for the stability of the CSC complex.

**Fig 4.**
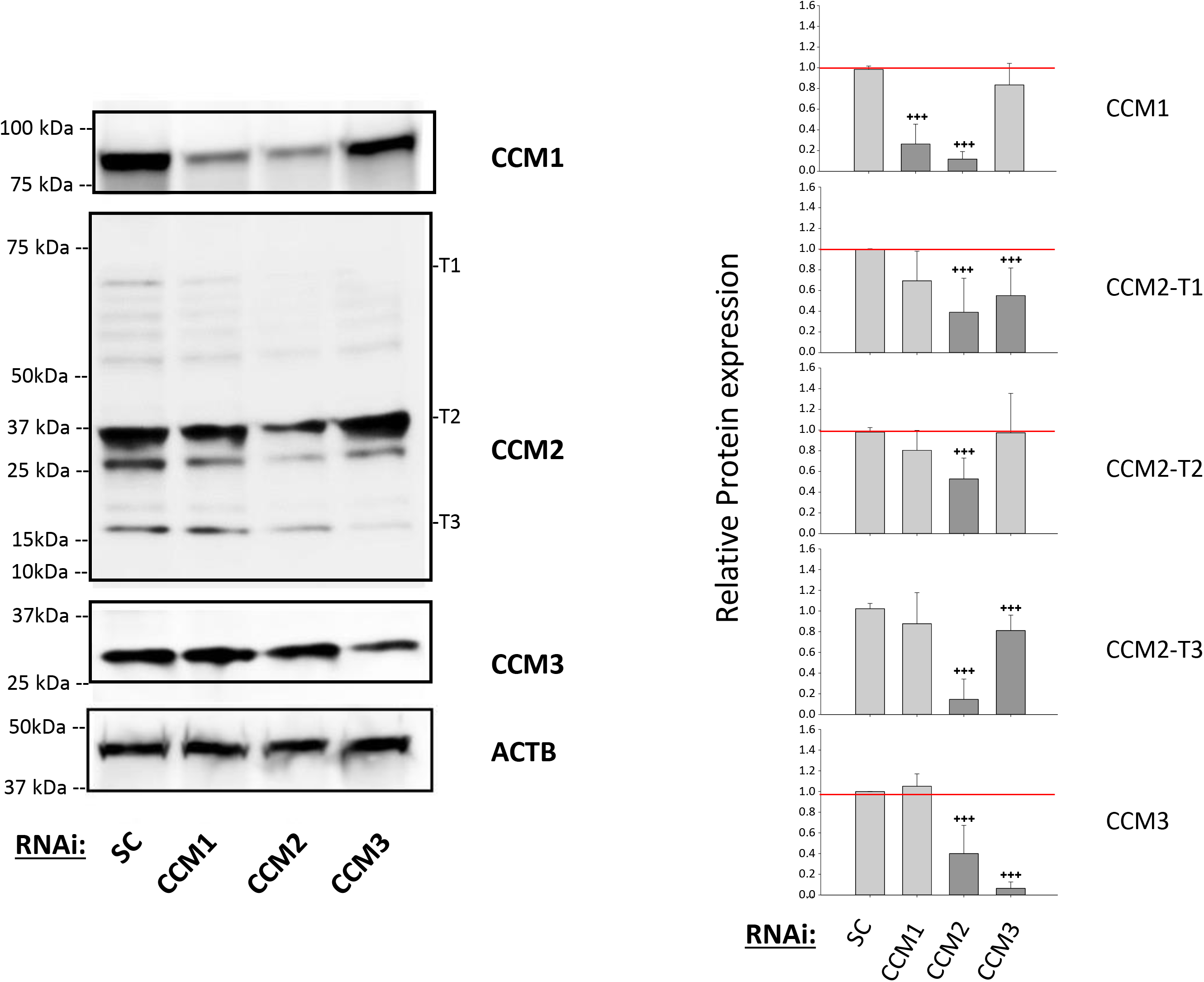
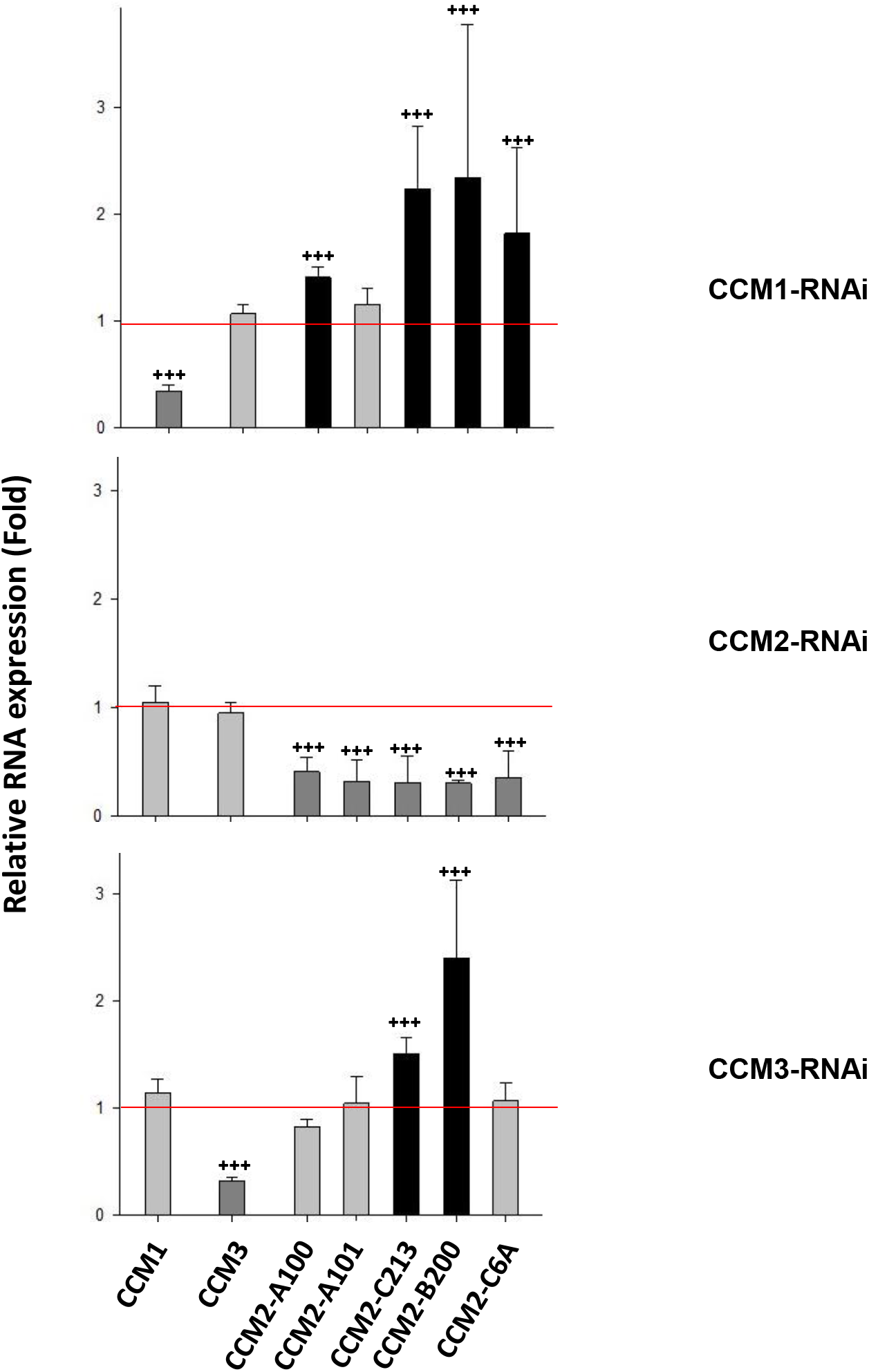
CCM2 is a cornerstone for the essential stability of CSC complex. Silencing of CCM2 decreases the expression level of both CCM1/3 proteins in T47D cells. **A)**. After silencing all three CCMs (1, 2 or 3) for 48 hrs, the expression levels of all three CCM proteins were efficiently targeted and silenced, however, a significantly decreased expression of both CCM1/3 proteins were observed in CCM2-KD T47D cells (Left upper and lower panels). The relative expression levels of CCM1/3 and three major isoforms of CCM2 proteins were measured through quantification of band intensities and normalized against β-actin (ACTB) followed by SC controls, and represented with bar plots where light grey bars represent no change and dark grey bars illustrate decreased expression (right panel) (n=3). **B)**. A significantly increased RNA level of CCM2 isoforms in silenced CCM1 (CCM1-KD) and CCM3 (CCM3-KD) T47D cells were observed. After silencing all three CCMs (1, 2 or 3) for 48 hrs, the relative RNA expression levels of CCM1, CCM3, and 5 isoforms of CCM2 in CCMs-KD in T47D cells were measured by RT-qPCR (Fold) and represented with bar plots where light grey bars represent no change, dark grey bars for decreased relative RNA levels, and black bars for increased relative RNA levels (n=3) (More detailed in Figure S4D).

### Relationship among classic, non-classic PRG receptors, and the CSC in hormone treated PR(+) T47D cells using omic approaches

Bioinformatics analyses identified saturable, high-affinity binding sites for PRG (*4, 6*), but no binding site for MIF in mPRs was discovered (*4*). Unlike PR1/2 which bears the same binding site for both mPRs and MIF, with MIF having higher binding affinity. The binding preference of MIF to PR1/2 but not mPRs might differentiate classic PRG actions from non-classic mPR-mediated ones (*4, 6*).

To define MIF binding partners in the synergistic action with PRG, we examined the expression patterns of T47D cells under either combined (MIF+PRG) or MIF-only treatments, at both transcriptional and translational levels using high throughput omics, including RNAseq and LC MS/MS. Among identified DEGs, we were able to visualize hierarchical clustering, and found similar patterns in both intersection and union of DEGs between two treatments (Figs. 5A-B). Similar phenomena were observed in DEGs found between the two treatments (Figs. 5C-D), suggesting shared signaling cascade variations between them. Finally, several key signaling cascades in tumorigenesis were most frequently observed in KEGG pathways and shared by both treatments, including cellular senescence, cell cycle, p53, microRNA’s in cancer, AMPK, MAPK, WNT, RAS and hormone signaling pathways (Figs. 5E-F)(Figs S5A1-5E2). Similar results were observed in PR(-) breast cancer cells (in preparation), leading us to propose that MIF binds to mPRs as well as synergistically with PRG. Interestingly, our RNAseq data confirmed the decreased expression seen with CCM2 isoforms (Fig. 2E) under steroid actions (Tables S5 and S7E). Similarly, we also observed decreased expression of PAQR8 in both MIF and MIF+PRG treated samples as well as increased expression of PAQR5 (Tables S5 and S7E) in accordance with our RT-PCR data (Fig. 3A), further validating our results and supporting our proposed model (Fig. 3G).

**Figure 5:**
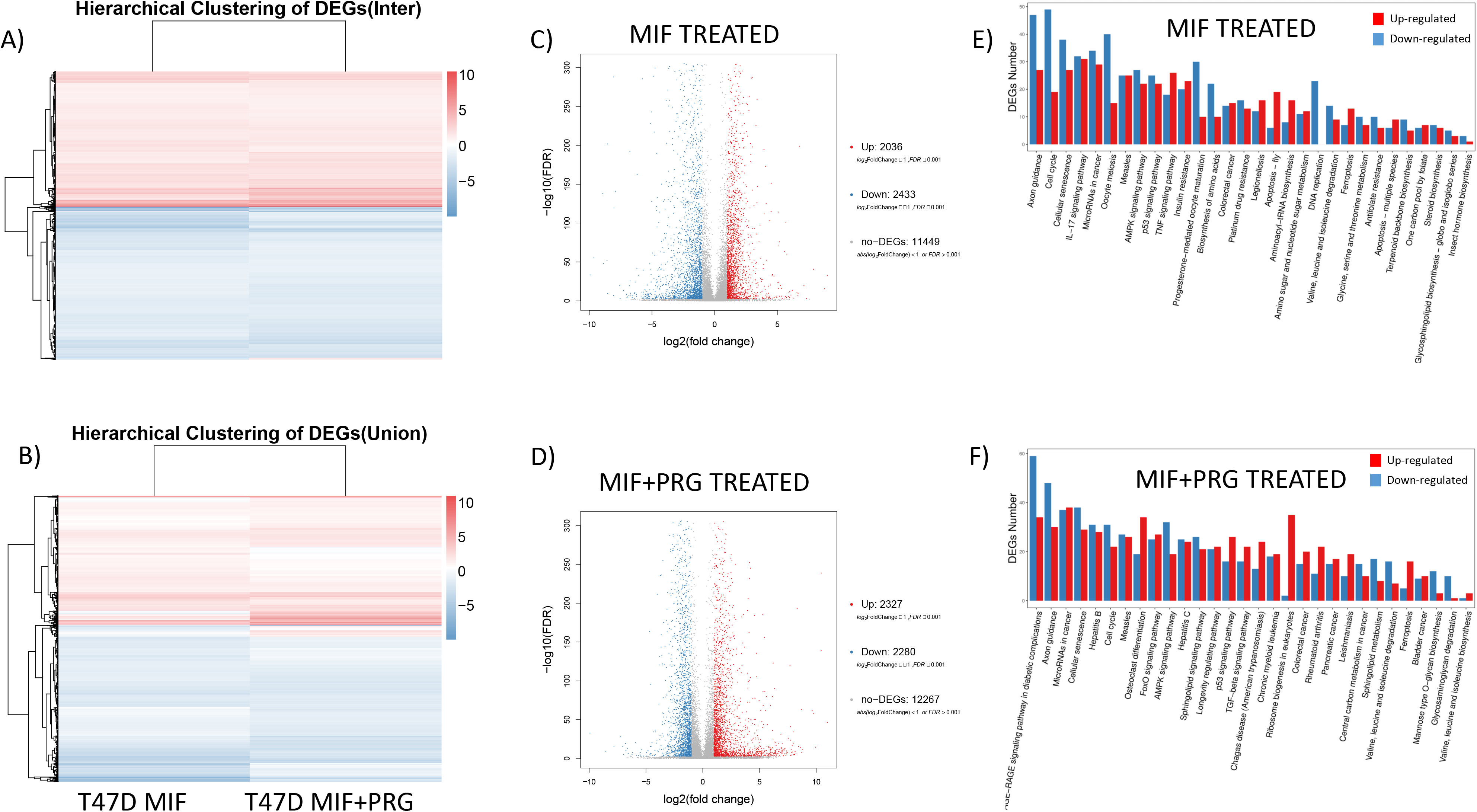
Differentially expressed gene (DEG) detection through high-throughput sequencing: *A-B)*. Cluster software and Euclidean distance matrixes were used for the hierarchical clustering analysis of the expressed genes (RNA) and sample program at the same time to generate the displayed Heatmap of hierarchical clustering for the intersection of DEGs *(A)* or union of DEGs *(B)* of expression clustering scheme; x axis represents each comparing sample and Y axis represents DEGs. Coloring indicates the log2 transformed fold change (high: red, low: blue). *C-D)* Volcano plot of DEGs for mifepristone treated cells *(C)* or Mifepristone + progesterone *(D)*; X axis represents log2 transformed fold change. Y axis represents -log10 transformed significance. Redpoints represent up-regulated DEGs. Blue points represent down-regulated DEGs. Gray points represent non-DEGs. *E-F).* Pathway functional enrichment result for up/down regulation of genes with Mifepristone treatment *(E)* or Mifepristone + Progesterone *(F).* X axis represents the terms of Pathway. Y axis represents the number of up/down regulated genes.

Using a similar approach, we analyzed our proteomic data with additional samples from silencing three CCMs genes (Tables S6A-D). After comparing each silenced treated sample to its respective control, we pooled the significant, differentially expressed proteins to represent a disturbed CSC since all 3 proteins are required to form the CSC (*8–10*). Similar to our RNA results, hierarchical clustering showed similar pattern with less degree between both hormone treatments (Figs 6A-B), supporting our RNA data conclusion and further emphasizing the different regulatory mechanisms of sex steroid actions at transcriptional and translational levels. However, there is significant differences in the CSC disruption group compared to both hormone-treated groups, indicating the independent role of the CSC in this signaling network (Figs. 6A-C). Interestingly, when we identified all the differentially expressed proteins (DEPs), we found that all three treatment groups displayed a larger amount of significantly down-regulated proteins compared to up-regulated proteins (Figs. 6D-F), which was not the case for our RNAseq analysis. Similar to our KEGG analysis with RNAseq data, our proteomics data for both hormone treatments (Figs. 6G-H), also showed differentially expressed proteins in cell cycle, AMPK and apoptosis signaling pathways (compare Figs. 5E-F, 6G-H). Shared pathways between hormone treatments at the translational level included proteoglycans in cancer, further supporting the intricate balance between the CSC and steroid actions in tumorigenesis. When evaluating a disrupted CSC, it was quite a surprise to see that the most amount of DEPs are involved in pathways in cancer signaling cascades, further validating the CSCs novel role in tumorigenesis (Fig. 6I). Additionally, Ras signaling, apoptosis, tight junction and regulation of actin/cytoskeleton were among some of the other signaling pathways affected with a disrupted CSC in PR(+) T47D cells (Fig. 6I). Our proteomics data further supports our RNAseq conclusions that both MIF and PRG bind to mPRs to modulate their synergistic effects.

**Figure 6:**
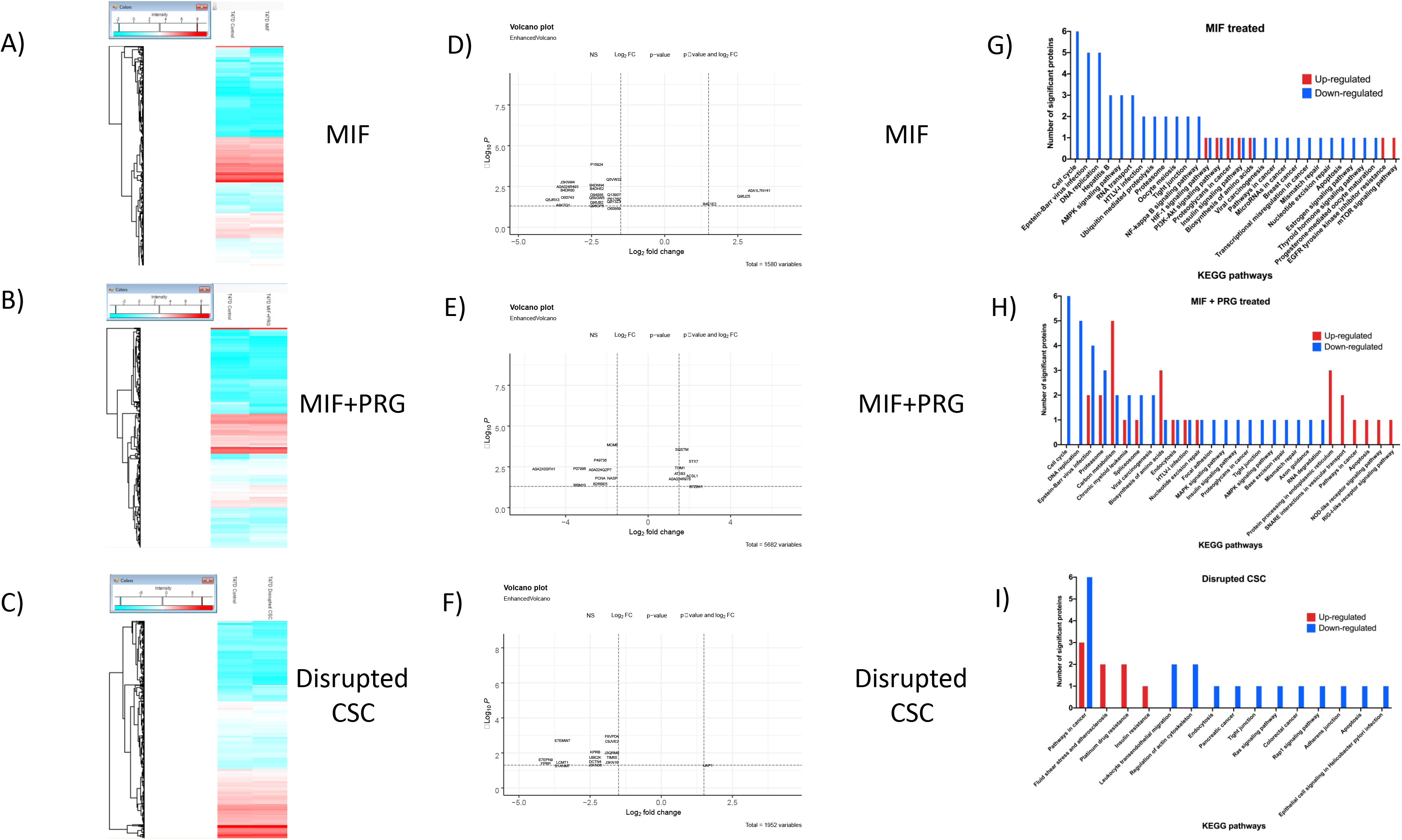
Differentially expressed gene (DEG) detection through proteomics: *A-C)*. Cluster software and Euclidean distance matrixes were used for the hierarchical clustering analysis of the expressed proteins and sample program at the same time to generate the displayed Heatmap of hierarchical clustering for the differentially expressed proteins (DEPs) under MIF-only treatment *(A)* MIF+PRG *(B)* and our disrupted CSC model (C) of expression clustering scheme; x axis represents each comparing sample and Y axis represents DEPs. Coloring indicates the log2 transformed fold change (high: red, low: blue). *D-F)* Volcano plot of DEGs for MIF-only treatment *(D)* MIF+PRG *(E)* and our disrupted CSC model (F); X axis represents log2 transformed fold change. Y axis represents -log10 transformed significance. Red points represent DEPs validated with both p-value and fold change significance, blue points showing significant p-value but lacking fold change criteria, green points having fold change but lacking significance and grey/black points representing non-DEPs. *G-I)* Pathway functional enrichment result for up/down regulation of genes with MIF-only treatment *(G)* MIF+PRG *(H)* and our disrupted CSC model (I). X axis represents the terms of Pathway. Y axis represents the number of up/down regulated proteins.

### Shared signaling cascades and relationship of classic, non-classic PRG receptors, and the CSC in hormone treated PR(+) T47D cells at both the transcriptional and translational levels

We recently identified differential expression patterns of the CSC complex are correlated with certain types and grades of major human cancers, especially in breast cancer (*10, 12, 13*), further validating a role for the CSC in tumorigenesis. To further delineate the role of not only the CSC, but also the role of classic and non-classic PRG receptors under hormone treatments, we compared our two omic approaches to identify altered signaling pathways at both the transcriptional and translational levels. Before doing this, we wanted to identify the signaling cascades and genes altered with each respective hormone combination. PRG-only comparisons (proteome and transcriptome) was assembled using published data to create a working database that could be compared to our MIF and MIF+PRG samples (*32, 33*). First, we identified overlaps, as well as unique genes/proteins at both the transcriptional (Fig. 7 top right panel) (Tables S7A-7D) and translational (Fig 7. top left panel) (Table S7E) levels for each hormone treatment. Once identified, we matched any shared DEGs among hormone treatments as well as our disrupted CSC model. We identified 5 genes overlapped with a disrupted CSC and MIF-only treatment, 8 genes overlapped with a disrupted CSC and PRG-only treatment while 6 overlaps were identified with a disrupted CSC and MIF+PRG treatment (Fig 7. Bottom panel). Using functional enrichment data for initially generated comparisons, we then identified shared pathways among the three hormone treatment groups with a disrupted CSC, which included cell cycle, apoptosis/cell death, kinase, DNA mechanisms and development signaling cascades. Shared Signaling between a disrupted CSC, MIF and MIF+PRG included cancer, RHO/GTPases, RNA mechanisms and autophagy/mitophagy pathways (Fig 7. Bottom panel) (Table S7F).

**Figure 7:**
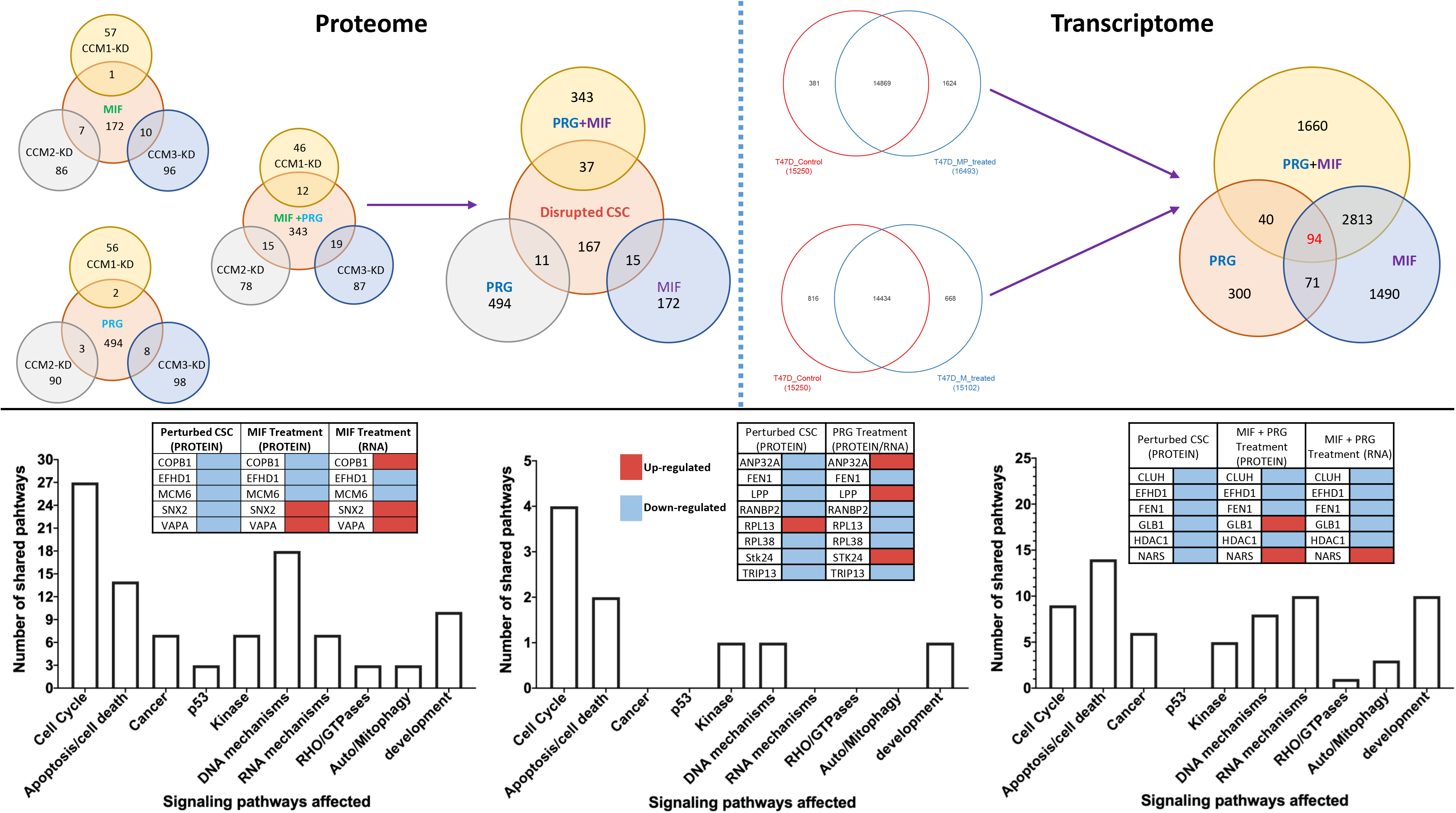
Overlapped genes using omics approaches: Gene overlaps were calculated at both the transcriptional (top right panel), proteome level (top left panel) and overlaps between both transcriptome and proteome (bottom panel) to elucidate any shared signaling pathways between treatment groups. Proteomic overlaps (top right panel) are demonstrated for both CSC disturbance as well as hormone treatments. Individual comparisons between hormone treatments and CCM knockdowns are illustrated and the combined knockdowns (disrupted CSC) compared to hormone treatments. Transcriptomic overlaps (top right panel) are demonstrated for hormone treatments. Individual comparisons between hormone treatments are illustrated as well as the combined hormone treatments comparison. Pathway classification and functional enrichment was performed on overlapped genes at both the proteome and transcriptome levels to evaluate any shared signaling mechanisms altered through a disturbed CSC and hormone treatments. For all analysis, PRG-only comparisons (proteome and transcriptome) were conducted using published proteomic/transcriptomic manuscripts to assemble a working database that could be compared to our MIF and MIF+PRG samples. If available, fold changes and p-values were obtained from the manuscripts to increase the depth of analysis of the multi-omics comparison.

## Discussion

In this study, we identified that the CSC has a major impact on PRG signaling through modulating the crosstalk between PR1/2 and mPRs/PAQRs in PR (+) T47D cells, providing novel insights into cellular coupling between CSC-mediated signaling and PRG modulated pathways via its classic and non-classical receptors involved in tumorigenesis. This could revolutionize the current understandings of molecular mechanisms of breast cancer tumorigenesis, leading to new therapeutic strategies.

There have been supporting evidences that PRG promotes cell proliferation (*34*) and inhibit cell death in human breast cancer T47D cells, suggesting it might not be the benign hormones for breast cancer, rather being pro-oncogenic (*35*). It has been speculated that PRG and its cellular metabolites promote cell proliferation through induced activation of MAPK signaling in both PR(+) and PR(-) breast cancer cell lines, which is independent of PR and ER (*36*), suggesting that mPRs may play a role in this signaling pathway. The involvement of MAPK signaling was confirmed in our omics data (Fig. 6H and Fig. S5C).

As an antagonist of PRG, MIF certainly earned its candidacy at the beginning for the treatment of breast, prostate, ovarian, endometrial cancers, and endometriosis and is currently on many active clinical trials (*24*). It was demonstrated that elevated levels of MIF can enhance the growth inhibition and induction of apoptosis triggered by high doses of PRG in PR(+) and PR(-) cancer cells (*37, 38*). It has also been reported that a clinically relevant dose of MIF significantly improves the treatment efficacy of cisplatin-paclitaxel chemotherapy regimens for human ovarian carcinoma cells (*39*).

However, many contradictory results have been reported regarding whether MIF has growth inhibition or stimulation for hormone-dependent breast cancer cells, as an anti-progestin (*40*). By screening 10 cancer cell lines with various genetic backgrounds, regardless of tissue of origin and hormone responsiveness, it was found that the anti-proliferative activity of MIF in cancer cells is independent of PR1/2 (*28*). Similarly, the cellular effects of MIF on proliferation is also widely observed (*41, 42*). It seems that pro- or anti-proliferative activity of MIF is determined by specific cell-types (*24*), concentration of MIF, as well as ratio of PR1/2 isoforms (*20*). It is essential to uncover the key mediators of MIF’s anti-tumor activity and its relationship to mPRs (*28*). Our findings that upon treatment of PRG/MIF, the altered expression pattern of three CCM proteins in T47D cells strongly suggests the involvement of the CSC in breast tumorigenesis. Furthermore, utilizing high throughput omic approaches, we have dissected alterations in signaling mechanisms under hormone therapy or disrupted CSC conditions affecting numerous essential signaling cascades. These signaling pathways include P53 signaling, P13K-AKT signaling, cell cycle, apoptosis, WNT, MAPK, various pathways in cancer as well as steroid hormone biosynthesis signaling pathways. These results solidify a novel signaling network among the CSC, classic and non-classic PRG receptors, which is dynamically modulated and fine-tuned with a series of feedback regulations.

## Conclusion

Despite its significance, the relationship between classic and non-classic PRG receptors has been drastically unexplored. It has been reported that activation of mPRs lead to activation of nuclear PR1/2, leading to an integrated model where steroid hormone-dependent mPRs contribute to later nuclear PR1/2 actions (*43*). In this study, we provide strong evidence that the CSC plays an essential role to bridge the crosstalk between PR1/2, mPRs/PAQRs and their ligand(s) to modulate this cascade among PR(+) cells. The possible convergence of genomic, non-genomic PRG actions and CSC signaling on their common cellular targets in PR(+) cells is an attractive model by which PRG or MIF can fine tune this intricate balance among these signaling pathways. This also raises the possibility that PRG or MIF may intervene different signaling pathways depending on the cellular context. Perturbation of this intricate balance, such as hormone therapy in the postmenopause or hormonal contraception regimens could result in increased risks in breast cancer or compromising tumor therapy.

## Supporting information

entire suppl materials

## Acknowledgments

We wish to thank Kamran Falahati, Mark Smith, Khalid Shoukat, Deepak Muthyala, Mike Yao, Yanchun Qu, Shen Sheng, Ahmed Badr, Junli Zhang, Amna Siddiqui, P. Dubey, Saafan Malik, and Edna Lopez at Texas Tech University Health Science Center El Paso (TTUHSCEP) for their technical help during the experiments.

## Author contributions

JZ: Conceptualization, Methodology, Investigation, Writing-Original draft preparation, Writing-Reviewing and Editing; JAF:Software, Data curation, Validation, Writing-Reviewing and Editing, XTJ: Investigation, Visualization; BG: Software, Data curation, Validation; AP:Investigation, Writing-Original draft preparation; CCE: Data curation, Validation.

## Data and materials availability

All data is available in the main text or the supplementary materials.

