## Supplementary material for "CCM signaling complex (CSC) coupling both classic and non-classic progesterone receptor signaling": entire suppl materials

*Supplemental Figures/tables are numbered to follow along with the numbering of figures in the main text, so there are some gaps in the numbering of the supplemental figures/tables. We have done this to aid in readability.

### Materials and Methods

#### Immunohistochemistry (IHC) and immunofluorescence (IF)

Deparaffinization of paraffin-embedded breast cancer tissue sections, antigen retrieval, blocking, antibody incubation, and imaging parameters were conducted as previously described (*1*). IHC/IF staining methods, imaging and quantification were performed as previous described (*1*), Antibody information is detailed in Suppl. Table 1.

#### Cell culture and treatments, real-time quantitative PCR (qPCR) and western blot analysis

T47D cells were cultured in RPMI1640 medium following manufacturer's instructions (ATCC), when cells reached 80% confluency, cells were treated with either vehicle control (ethanol/DMSO, VEH), mifepristone (MIF, 20 μM), progesterone (PRG, 20 μM), estrogen (EST, 10 nM), gradient concentrations of progesterone (PRG, 1-80 μM) and mifepristone (MIF, 5-160 μM), combined hormones (PRG+MIF; 20μM each), or media only (Untreated) respectively for steroid treatments. For RNA knockdown experiments, 80% confluent T47D Cells were transfected with a set of siRNAs, targeting specific genes (Suppl. Table 2), by RNAiMAX (Life Technologies) as described before (*1, 2*). After cells were treated in various conditions, they were harvested, and RNA expression levels of *CCMs, PR* and *mPRs* genes were determined through quantitative PCR (qPCR) (Suppl. Table 3). qPCR was performed to quantify the RNA levels using Power SYBR Green Master Mix with ViiA7 Real-Time PCR System (Applied Biosystems), and data was analyzed with DataAssist (ABI) and Rest 2009 software (Qiagen). All experiments were performed with triplicates as described before (*1, 2*). The relative expression levels of candidate proteins were measured with Western blots (WB) (Suppl. Table 1) as described before (*1, 2*).

#### Proteomic and transcriptomic (RNAseq) analysis

RNAseq, proteomic sample preparation, data acquisition to assemble interactomes, DEG analysis, pathways analysis and correlations between transcriptomics and proteomics was performed as previously described (*3, 4*).

**Abbreviations**: Cerebral cavernous malformation (CCM), CCM signaling complex (CSC), progesterone (PRG), mifepristone (MIF), membrane progesterone receptors (mPRS)

**Keywords:** Cerebral cavernous malformation, CCM signaling complex (CSC), progesterone (PRG), mifepristone (MIF), classic nuclear progesterone receptors (PR1/2), non-classic membrane progesterone receptors (mPRs/PAQRs), genomic action, non-genomic actions (rapid actions), proteomics, transcriptomics, RNAseq.

### Supplementary Text

Supplemental Figures/tables are numbered to follow along with the numbering of figures in the main text to aid in readability.

**Fig. S1.**

**Expression levels of CCM proteins in various breast cancer cells. A.** Significant increased protein expressions of PAQR7 with HRP/DAB staining in breast tumors (Breast Carcinoma) can be visualized from a selected set of breast tissue-pairs (left panel) and representative quantification displayed (ROI, tumor= 11363; normal= 7008, middle panel). Statistically significant increased levels of PAQR7 protein were found in tumors from the entire collection of paired samples (ROI=7008-14148/sample, right panel, P =< 0.001, n=10), **B.** Subtypes of breast cancer cell lines (MDA-MB-231, MCF7, MDA-MB-468, BT474, T47D, MDA-MB-453) and prostate cancer cell lines (PC3, and C4-2) were utilized to screen the expression patterns of CCM2 protein isoforms (A-I). T47D, which displays expression of all CCM2 isoforms, had the highest expression of the three most abundant isoforms, T1, T2 and T3, along with PC3 prostate cancer cell line. Expression and composition of CCM2 isoforms do not appear to follow a trend with either the immune-profile or molecular subtypes the cells originated from, indicating that protein expression and composition of CCM2 isoforms do not correlate with either immune-profiles or molecular subtypes of the breast cancer cells (bottom panel). **C.** In Figure 1 and Suppl. Fig. 1A, HRP/DAB staining (1A), the Red/Brown color from HRP/DAB reactivity with any specific antibody is quantified and averaged between the red and green channel quantification. In fluorescent staining experiments (panel 1B), CCM1 and CCM3 were quantified through ROI intensities using wavelength channels 488 and 647nm, respectively. Data for both microscopy approaches were normalized against its respective internal controls using the blue channel for cell nuclei (HRP/DAB) or 408nm wavelength for DAPI (fluorescent) and background staining. For each section pair, Region of Interest (*ROI*) i*ntensities* were automatically quantified (over 1000 times/per section). The relative expression levels of CCM1 and CCM3 proteins were measured through quantification of band intensities and normalized against β-actin (ACTB). All data from entire collections ($n>10$) were normalized by normal tissue among each tissue pair. All imaging data were acquired using Nikon EclipseTi confocal microscope and quantified with Elements Analysis software (Nikon). In all bar plots, red line is the control baseline for fold change measurements (-/+) and *** above any bar graphs indicate $P \leq 0.001$ for un-paired *t*-test.

**Fig. S2.**

**Expression levels of CCM proteins in PR-positive [PR(+)] breast cancer cells T47D.** In Figure 2, the relative transcription expression changes of PR isoforms in CCMs KD in T47D cells were measured by qPCR (Fold) (n=3). Red line indicates control baseline for fold change measurements (-/+). **, *** above bar indicates $P \le 0.01$ or 0.001 for paired t-test, respectively. In HRP/DAB staining, the Red/Brown color from HRP/DAB reactivity with a specific antibody is quantified and averaged between the red and green channel quantification and cell nuclei are quantified with the blue channel. Relative expression levels of CCM1 and CCM3 were measured through quantification of band intensities and normalized against β-actin (ACTB) and scrambled controls (SC). Data were normalized against its respective control using the blue channel for cell nuclei and background staining as described earlier and quantified with Nikon Elements Analysis software.

**Fig. S3.**

**Expression levels of membrane progesterone receptors (mPRs/PAQRs) are modulated by progesterone (PRG) and mifepristone (MIF) and the CSC in PR positive PR(+) T47D cells. A.** Protein expression levels of all mPRs (PAQRs, PGRMC1) are not influenced by silencing either androgen receptor (AR), glucocorticoid receptor (GR) or progesterone receptor (PR1/2) genes. After silencing AR, GR, and PR1/2 genes for 48 hrs, no significant changes in protein expression levels of mPRs (PAQRs, PGRMC1) in T47D cells were observed, indicating protein expression levels of mPRs are not influenced by these steroid receptors. **B**. RNA expression levels of mPRs (PAQR 7, 8, 9) are not influenced by silencing progesterone receptor (PR1/2) genes. After silencing PR1/2 genes for 48 hrs, no significant change in RNA expression levels of three representative mPRs (PAQR 7, 8, 9) were observed. Relative RNA expression changes of mPRs (PAQR 7, 8, 9) in T47D cells were measured by qPCR in (Fold changes) (triplicates per experiment, n=3). **C**. In Figure 3, Relative RNA expression changes were measured by qPCR (Fold changes) and normalized to scramble control (red line, triplicates per experiment, n=3). Relative expression levels of proteins were measured through quantification of band intensities and normalized against either α-actinin (ACTN1) or β-actin (ACTB) followed by SC controls (red line), and represented with bar plots. In all bar plots, red line is the control baseline for fold change measurements (-/+). **, *** above bar indicates $P \leq 0.01$ or 0.001 for paired t- test, respectively.

**Fig. S4.**

**CCM2 is a cornerstone for the essential stability of the CSC complex. A.** Silencing of CCM2 decreases the expression level of both CCM 1/3 proteins in 293T cells. **A.1)**. After silencing CCMs (1, 2 or 3) for 48 hrs, the expression levels of all three CCM proteins were efficiently silenced, however, a significantly decreased expression of both CCM1 and CCM3 proteins were observed in CCM2-KD 293T cells (Left upper and lower panels). The relative expression levels of CCM (1, 2 and 3) proteins were measured through quantification of band intensities and normalized against β-actin (ACTB) followed by SC controls, and represented with bar plots where light grey bars represent no change and dark grey bars illustrate decreased relative protein levels (right panel) (n=3). **A.2)**. A significantly increased RNA level of some CCM2 isoforms in silenced CCM1 (CCM1-KD) and CCM3 (CCM3-KD) 293T cells were observed. The relative transcription expression changes of CCM1, CCM3, and 5 isoforms of CCM2 in CCMs-KD 293T cells were measured by RT-qPCR (Fold), normalized against SC controls (red line) and represented with bar plots where light grey bars represent no change, dark grey bars illustrate decreased relative RNA levels, and black bars display increased relative RNA levels (n=3). **B.** Silencing of CCM2 decreases the expression level of both CCM 1/3 proteins in human brain microvascular endothelial cells (HBMVEC). **B.1)**. After silencing all three CCMs (1, 2 or 3) for 48 hrs, the expression levels of all three CCM proteins were efficiently targeted and silenced, however, a significantly decreased expression of both CCM1/3 proteins were observed in CCM2-KD HBMVEC cells (Left upper and lower panels). The relative expression levels of CCMs (1, 2, 3) proteins were measured through quantification of band intensities and normalized against β-actin (ACTB) followed by SC controls (red line), and illustrated with bar plots where light grey bars represent no change and dark grey bars display decreased relative protein levels (right panel) (n=3). **B.2)**. A significant increased RNA level of CCM2 isoforms in both CCM1-KD and CCM3-KD HBMVEC cells were observed. The relative transcription expression changes of CCM1, CCM3, and 5 isoforms of CCM2 in CCMs KD HBMVEC cells were measured by RT-qPCR (Fold) and illustrated in the bar plots where light grey bars represent the relative RNA levels of CCM1, dark grey bars display relative RNA levels of CCM2 isoforms, and black bars for the relative RNA levels of CCM3 (n=3). **C.** Knockout of CCM2 decreases the expression level of CCM1/3 proteins in zebrafish embryo. **C.1**). A significant decreased expression of both CCM1/3 proteins were observed in CCM2-knockout (KO) zebrafish embryo (*vtn*) (Left upper and lower panels). The relative expression levels of CCMs (1, 2, 3) proteins were measured through quantification of band intensities and normalized against α-actinin (ACTN1) followed by WT strains (red line), and illustrated with bar plots where light grey bars represent no change and dark grey bars illustrate significantly decreased relative protein levels (right panel) (n=3). **C.2)**. A significant increased RNA level of CCM2 isoforms in CCM1-KO zebrafish (*san*) were observed. The relative RNA expression changes of CCM1 and 4 isoforms of CCM2 in CCM1-KO and CCM2-KO zebrafish (*san, vtn, respectively*) were measured by RT-qPCR (Fold) and represented with bar plots where light grey bars represent the relative RNA level of CCM1 and dark grey bars illustrate the relative RNA level of CCM2 (n=3). Primer information for zebrafish RT-qPCR can be provided upon request. **D)** In all bar plots, red line is the control baseline for fold change measurements (-/+). **, *** above bar indicates P ≤ 0.01 or 0.001, respectively, for paired t- test.

**Fig. S5. Aathways analysis.**

Fig. S5A. **KEGG pathways analysis of RNAseq data for P53 and Breast cancer signaling pathways with hormone treated T47D cells**: Using Differentially Expressed Genes (DEGs), we performed KEGG pathway classification and functional enrichment. With the KEGG annotation result, we classify DEGs according to official classification, and we also perform pathway functional enrichment using phyper, a function of R. **Top left panel**). P53 signaling pathways under MIF treatment, **Bottom left panel**). P53 signaling pathways under MIF+PRG treatment, **Top right panel**). Breast cancer signaling pathways under MIF treatment, **Bottom right panel**). Breast cancer signaling pathways under MIF+PRG treatment. Up-regulated genes are marked with red borders and down-regulated genes with green borders. Non-change genes are marked with black borders.

Fig. S5B1. **KEGG pathways analysis of RNAseq data for cell cycle/apoptosis, JAK-STAT, and TGF-β signaling pathways with MIF treated T47D cells**: Using Differentially Expressed Genes (DEGs), we performed KEGG pathway classification and functional enrichment. With the KEGG annotation result, we classify DEGs according to official classification, and we also perform pathway functional enrichment using phyper, a function of R. *a)*. Apoptosis signaling pathways *(b)* JAK-STAT signaling pathways *c)* Cell Cycle signaling pathways *(d)* TGF-ß signaling pathway. Up-regulated genes are marked with red borders and down-regulated genes with green borders. Non-change genes are marked with black borders.

Fig. S5B2. **KEGG pathways analysis of RNAseq data for cell cycle/apoptosis, JAK-STAT, and TGF-β signaling pathways with MIF+PRG treated T47D cells**: Using Differentially Expressed Genes (DEGs), we performed KEGG pathway classification and functional enrichment. With the KEGG annotation result, we classify DEGs according to official classification, and we also perform pathway functional enrichment using phyper, a function of R. *a)*. Apoptosis signaling pathways *(b)* JAK-STAT signaling pathways *c)* Cell Cycle signaling pathways *(d)* TGF-ß signaling pathway. Up-regulated genes are marked with red borders and down-regulated genes with green borders. Non-change genes are marked with black borders.

Fig. S5C1. **KEGG pathways analysis of RNAseq data for WNT/MAPK/RAS/P13K-AKT signaling pathways with MIF treated T47D cells**: Using Differentially Expressed Genes (DEGs), we performed KEGG pathway classification and functional enrichment. With the KEGG annotation result, we classify DEGs according to official classification, and we also perform pathway functional enrichment using phyper, a function of R. *a)*. WNT signaling pathway *(b)* P13K-AKT signaling pathway *c)* MAPK signaling pathways *(d)* RAS signaling pathway. Up-regulated genes are marked with red borders and down-regulated genes with green borders. Non-change genes are marked with black borders.

Fig. S5C2. **KEGG pathways analysis of RNAseq data for WNT/MAPK/RAS/P13K-AKT signaling pathways with MIF+PRG treated T47D cells**: Using Differentially Expressed Genes (DEGs), we performed KEGG pathway classification and functional enrichment. With the KEGG annotation result, we classify DEGs according to official classification, and we also perform pathway functional enrichment using phyper, a function of R. *a)*. WNT signaling pathway *(b)* P13K-AKT signaling pathway *c)* MAPK signaling pathways *(d)* RAS signaling pathway. Up-regulated genes are marked with red borders and down-regulated genes with green borders. Non-change genes are marked with black borders.

Fig. S5D1. **KEGG pathways analysis of RNAseq data for Cancer signaling pathways**

**with MIF treated T47D cells**: Using Differentially Expressed Genes (DEGs), we performed KEGG pathway classification and functional enrichment. With the KEGG annotation result, we classify DEGs according to official classification, and we also perform pathway functional enrichment using phyper, a function of R. *a)*. Proteoglycans in cancer signaling pathways *(b)* Cancer signaling pathways. Up-regulated genes are marked with red borders and down-regulated genes with green borders. Non-change genes are marked with black borders.

Fig. S5D2. **KEGG pathways analysis of RNAseq data for Cancer signaling pathways with MIF+PRG treated T47D cells**: Using Differentially Expressed Genes (DEGs), we performed KEGG pathway classification and functional enrichment. With the KEGG annotation result, we classify DEGs according to official classification, and we also perform pathway functional enrichment using phyper, a function of R. *a)*. Proteoglycans in cancer signaling pathways *(b)* Cancer signaling pathways. Up-regulated genes are marked with red borders and down-regulated genes with green borders. Non-change genes are marked with black borders.

Fig. S5E1. **KEGG pathways analysis of RNAseq data for Steroid Hormone signaling pathways with MIF treated T47D cells**: Using Differentially Expressed Genes (DEGs), we performed KEGG pathway classification and functional enrichment. With the KEGG annotation result, we classify DEGs according to official classification, and we also perform pathway functional enrichment using phyper, a function of R. *a)*. Steroid hormone biosynthesis signaling pathway *(b)* Progesterone mediated oocyte maturation. Up-regulated genes are marked with red borders and down-regulated genes with green borders. Non-change genes are marked with black borders.

Fig. S5E2. **KEGG pathways analysis of RNAseq data for Steroid Hormone signaling pathways with MIF+PRG treated T47D cells**: Using Differentially Expressed Genes (DEGs), we performed KEGG pathway classification and functional enrichment. With the KEGG annotation result, we classify DEGs according to official classification, and we also perform pathway functional enrichment using phyper, a function of R. *a)*. Steroid hormone biosynthesis signaling pathway *(b)* Progesterone mediated oocyte maturation. Up-regulated genes are marked with red borders and down-regulated genes with green borders. Non-change genes are marked with black borders.

Table S1. **Antibodies used in this study**. Detailed information about application, antigen, clone code, secondary antibody, manufacturer, and category number are listed. * IHC, Immunohistochemistry; IF, immunofluorescence; DAB, HRP/DAB staining, WB, Western Blots.

Table S2. **Small double-stranded interfering RNAs (siRNA) used for gene silencing in RNA interference (RNAi) experiments.** Gene-specific knockdown was achieved by introducing small double-stranded siRNA oligos. Detailed information about application, siRNA oligo duplexes, manufacturers, and category numbers are listed.

Table S3. **Primers for quantitative PCR (qPCR) for the detection of differentially expressed targeted genes**. The sequence, qPCR product size and optimized annealing temperature of each primer pairs are listed.

Table S4. **N/A**

Table S5. **RAW data for RNAseq analysis:** Based on the gene expression level, we can identify the DEG (Differentially expressed genes) between samples or groups. We used Possion Distribution algorithms to detect the DEGs. The columns in the table represent the following information: GeneID: Gene ID, Length: Gene Length, Group1-Expression: Gene expression in group1, Group2-Expression: Gene expression in group2, log2FoldChange(group2/group1): The log2 value of ratio of Group1-Expression to Group2-Expression, FDR: Adjusted p-value, Up/Down-Regulation: Up/down-regulated, Pvalue: p-value, Symbol: Gene Symbol ID(Gene Name)

Table S6A. **RAW data for Proteomics analysis of T47D CCM1,2,3-KDs:** Based on the protein expression level, we can identify the DEPs (Differentially expressed Proteins) between samples or groups. The columns in the tables represent the following information: Protein name, Accession Number, Alternate ID, Molecular weight, T-Test: (p-value)($p < 0.05$), Fold Change by Category, and RAW spectral counts for each replicate.

Table S6B. **RAW data for Proteomics analysis of T47D MIF+PRG treated cells:** Based on the protein expression level, we can identify the DEP (Differentially expressed Proteins) between samples or groups. The columns in the tables represent the following information: Protein name, Accession Number, Alternate ID, Molecular weight, T-Test: (p-value)($p < 0.05$), Fold Change by Category, and RAW spectral counts for each replicate.

Table S6C. **RAW data for Proteomics analysis of T47D MIF+PRG treated cells-set 2:** Based on the protein expression level, we can identify the DEP (Differentially expressed Proteins) between samples or groups. The columns in the tables represent the following information: Protein name, Accession Number, Alternate ID, Molecular weight, T-Test: (p-value)($p < 0.05$), Fold Change by Category, and RAW spectral counts for each replicate.

Table S6D. **RAW data for Proteomics analysis of T47D MIF treated cells:** Based on the protein expression level, we can identify the DEP (Differentially expressed Proteins) between samples or groups. The columns in the tables represent the following information: Protein name, Accession Number, Alternate ID, Molecular weight, T-Test: (p-value)($p < 0.05$), Fold Change by Category, and RAW spectral counts for each replicate.

Table S7A. **Unique/shared Proteins identified between MIF treatment and Disrupted CSC (Individual).** Unique and overlapped genes displayed in Figure 7 (Top left panel) are provided with the alternate ID, T-Test: (p-value)($p < 0.05$), and Fold Change by Category.

Unique proteins found in CCM1-KD, CCM2-KD, CCM3-KD and MIF are provided as well as the shared proteins between MIF vs CCM1, MIF vs CCM2 and MIF vs CCM3.

Table S7B. **Unique/shared Proteins identified between PRG treatment (Curated database) and Disrupted CSC (Individual).** Unique and overlapped genes displayed in Figure 7 (Top left panel) are provided with the Gene Symbol, T-Test: (p-value)($p < 0.05$)(IF PROVIDED FOR PRG), and Fold Change by Category (IF PROVIDED FOR PRG). Unique proteins found in CCM1-KD, CCM2-KD, CCM3-KD and PRG are provided as well as the shared proteins between PRG vs CCM1, PRG vs CCM2 and PRG vs CCM3.

Table S7C. **Unique/shared Proteins identified between MIF+PRG treatment and Disrupted CSC (individual).** Unique and overlapped genes displayed in Figure 7 (Top left panel) are provided with the alternate ID, T-Test: (p-value)($p < 0.05$), and Fold Change by Category. Unique proteins found in CCM1-KD, CCM2-KD, CCM3-KD and MIF+PRG are provided as well as the shared proteins between MIF+PRG vs CCM1, MIF+PRG vs CCM2 and MIF+PRG vs CCM3.

Table S7D. **Unique/shared Proteins identified between Steroid treatment and Disrupted CSC (Pooled).** Unique and overlapped genes displayed in Figure 7 (Top left panel) are provided with the alternate ID, T-Test: (p-value)($p < 0.05$), and Fold Change by Category. Unique proteins found in CSC-KD (pooled CCM1-KD, CCM2-KD, CCM3-KD) and Steroid Treatments (PRG, MIF, MIF+PRG) are provided as well as the shared proteins between Steroid Treatments vs CSC-KD.

Table S7E. **Unique/shared Genes (RNAseq) identified between Steroid treatment groups.** Unique and overlapped genes displayed in Figure 7 (Top right panel) are provided with the gene symbol, p-value ($p < 0.05$), and Log2Fold Change. Unique proteins found in Steroid treatment groups are provided as well as the shared proteins between Steroid treatment groups.

Table S7F. **Shared functional enrichment pathways identified between Steroid treatments and Disrupted CSC (Pooled) at both the transcriptional and translational levels.** Overlapped Biological Processes, Molecular Functions, KEGG pathways and Reactome pathways displayed in Figure 7 (bottom panel) are provided with the GO#, and pathway description. The pathways analyzed included cell cycle, apoptosis/cell death, cancer, P53, kinase, DNA/RNA mechanisms, RHO/GTPases, Autophagy, Mitophagy, and development signaling cascades (color coding provided in tab 1). Genes identified in each comparison are also provided in a color-coded table that specifies up/down regulation to see trends across the transcriptional/translational level for each identified gene and treatment comparison.

1A

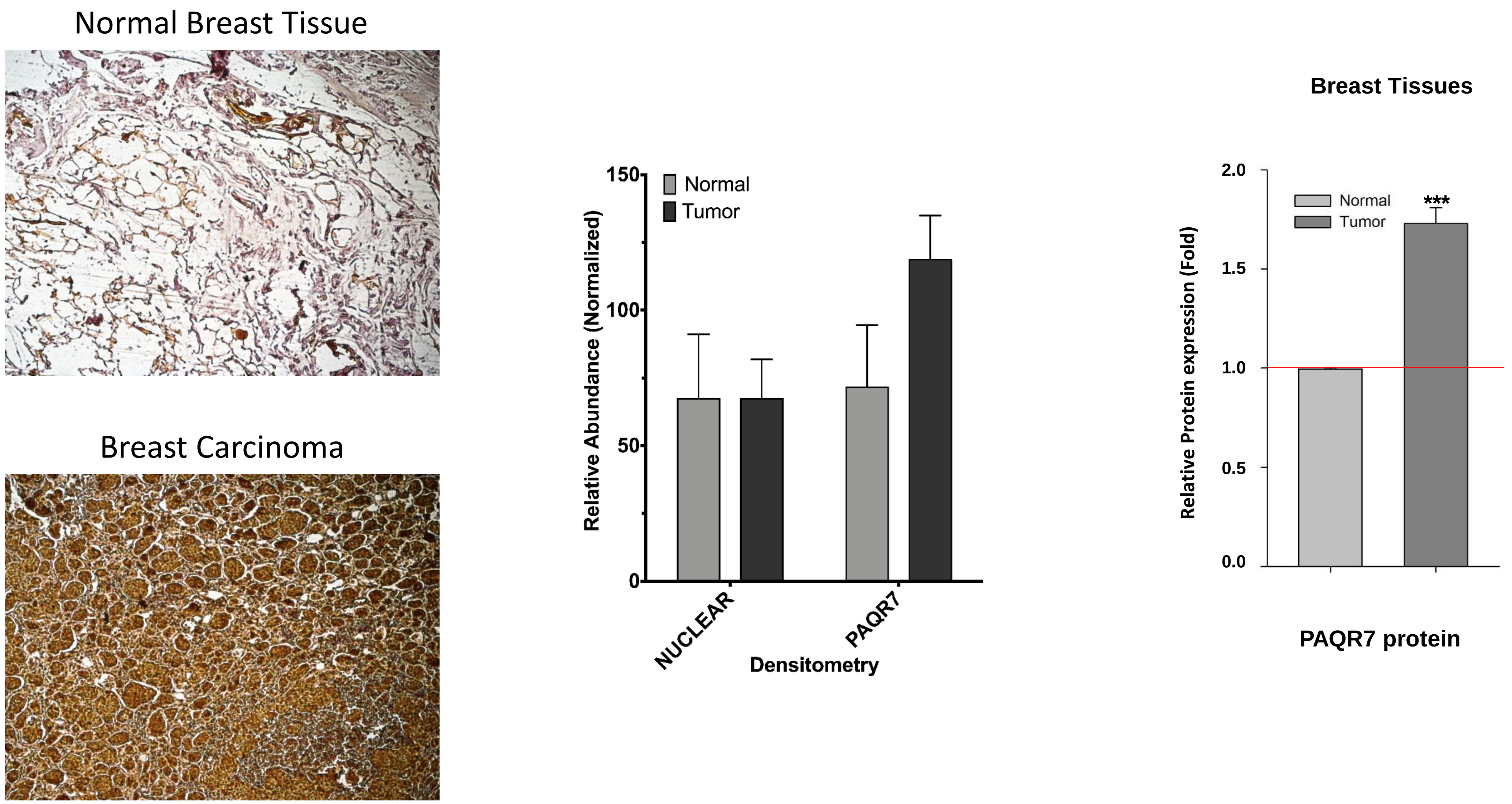
Normal Breast Tissue
Breast Carcinoma
150
100
50
0
Normal
Tumor
Relative Abundance (Normalized)
NUCLEAR
PAQR7
Densitometry
Breast Tissues
2.0
1.5
1.0
0.5
0.0
Normal
Tumor
***
Relative Protein expression (Fold)
PAQR7 protein

**Suppl. Fig. 1B**

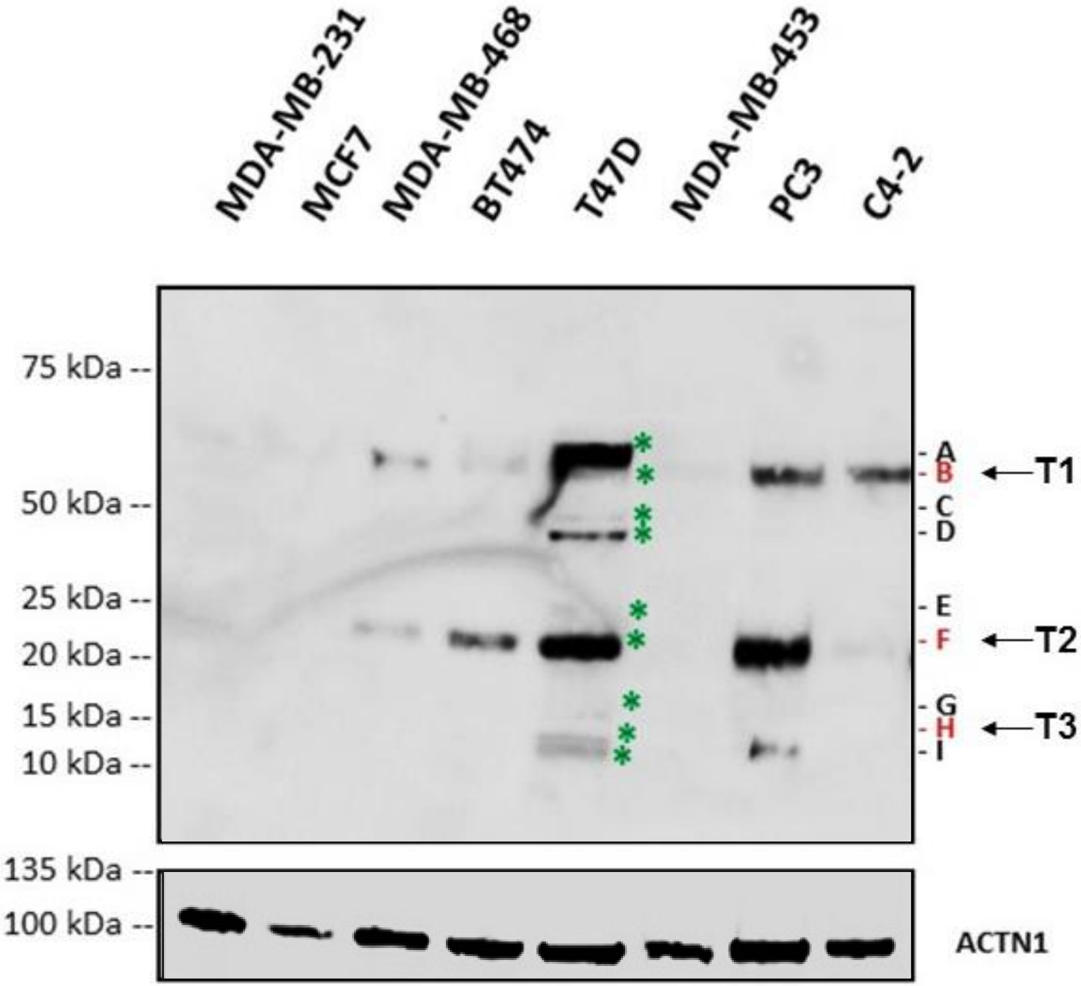


| | | | | | | | | |
|---|---|---|---|---|---|---|---|---|
| ER | – | † | – | † | † | – | † | † |
| PR | – | † | – | † | † | – | – | – |
| AR | † | † | † | † | † | † | – | † |
| GR | † | † | † | † | – | † | † | – |
| HER2 | † | † | – | † | – | † | – | – |

**Suppl. Fig. 2**

All information in legends.

**Suppl Fig. 3A**

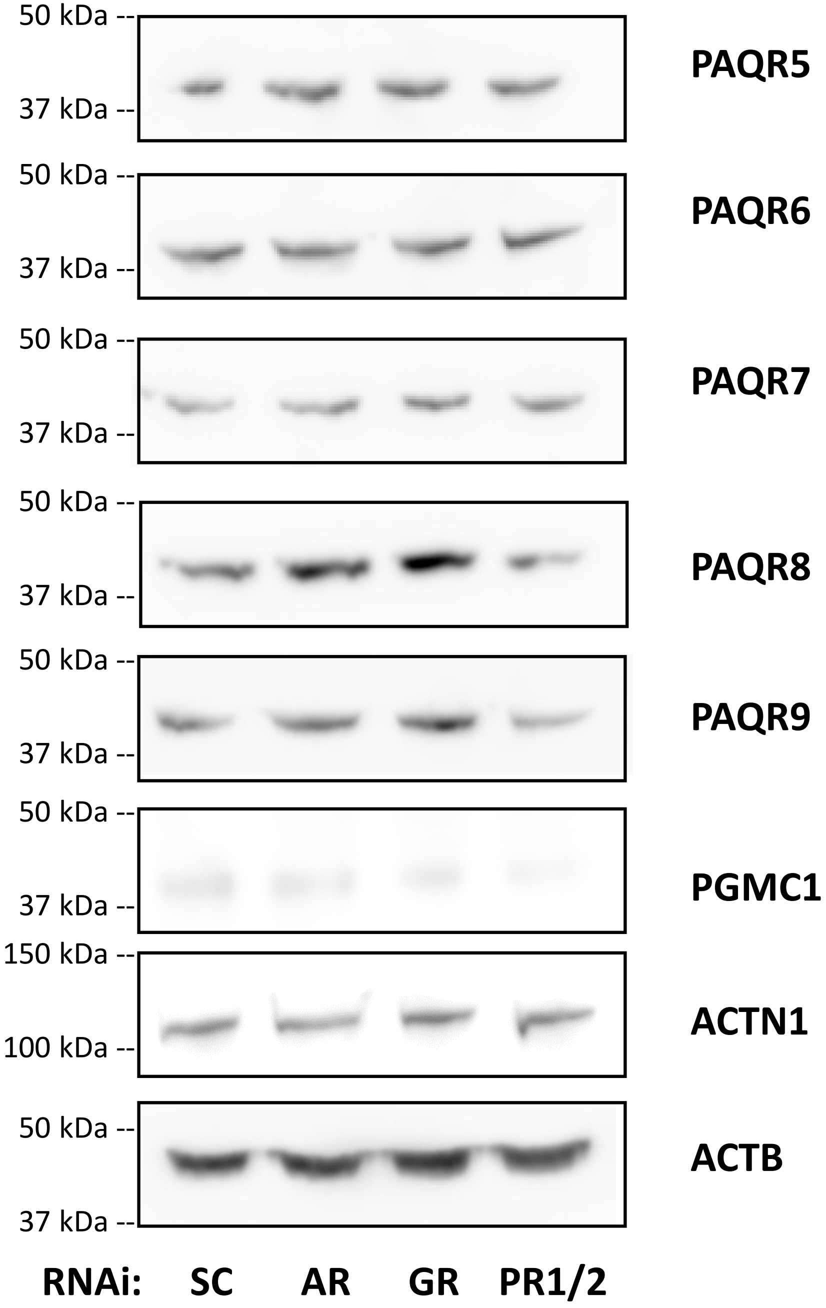
50 kDa
37 kDa
PAQR5
50 kDa
37 kDa
PAQR6
50 kDa
37 kDa
PAQR7
50 kDa
37 kDa
PAQR8
50 kDa
37 kDa
PAQR9
50 kDa
37 kDa
PGMC1
150 kDa
100 kDa
ACTN1
50 kDa
37 kDa
ACTB
RNAi: SC AR GR PR1/2

**Suppl Fig. 3B**

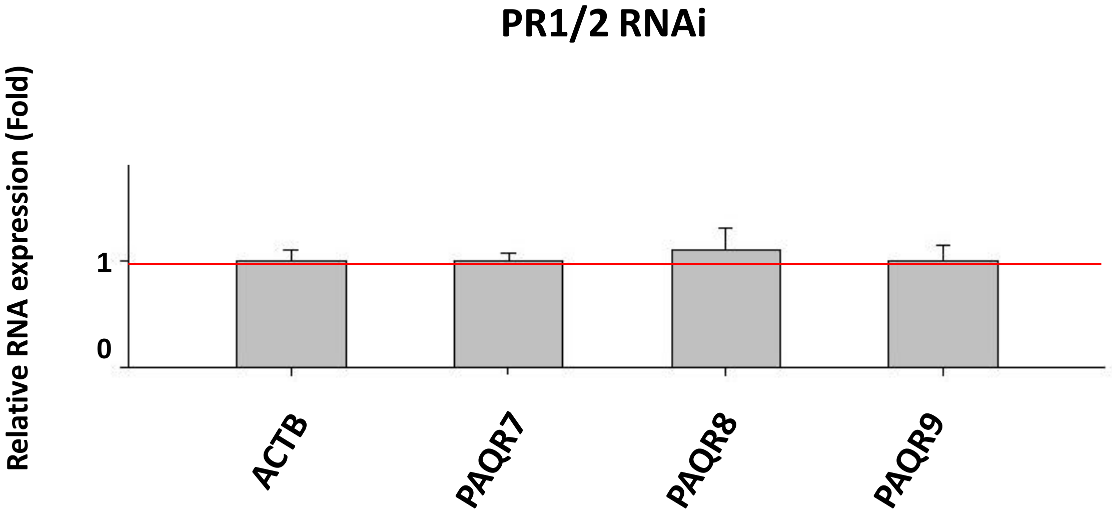
PR1/2 RNAi
Relative RNA expression (Fold)
1
0
ACTB
PAQR7
PAQR8
PAQR9

4A.1

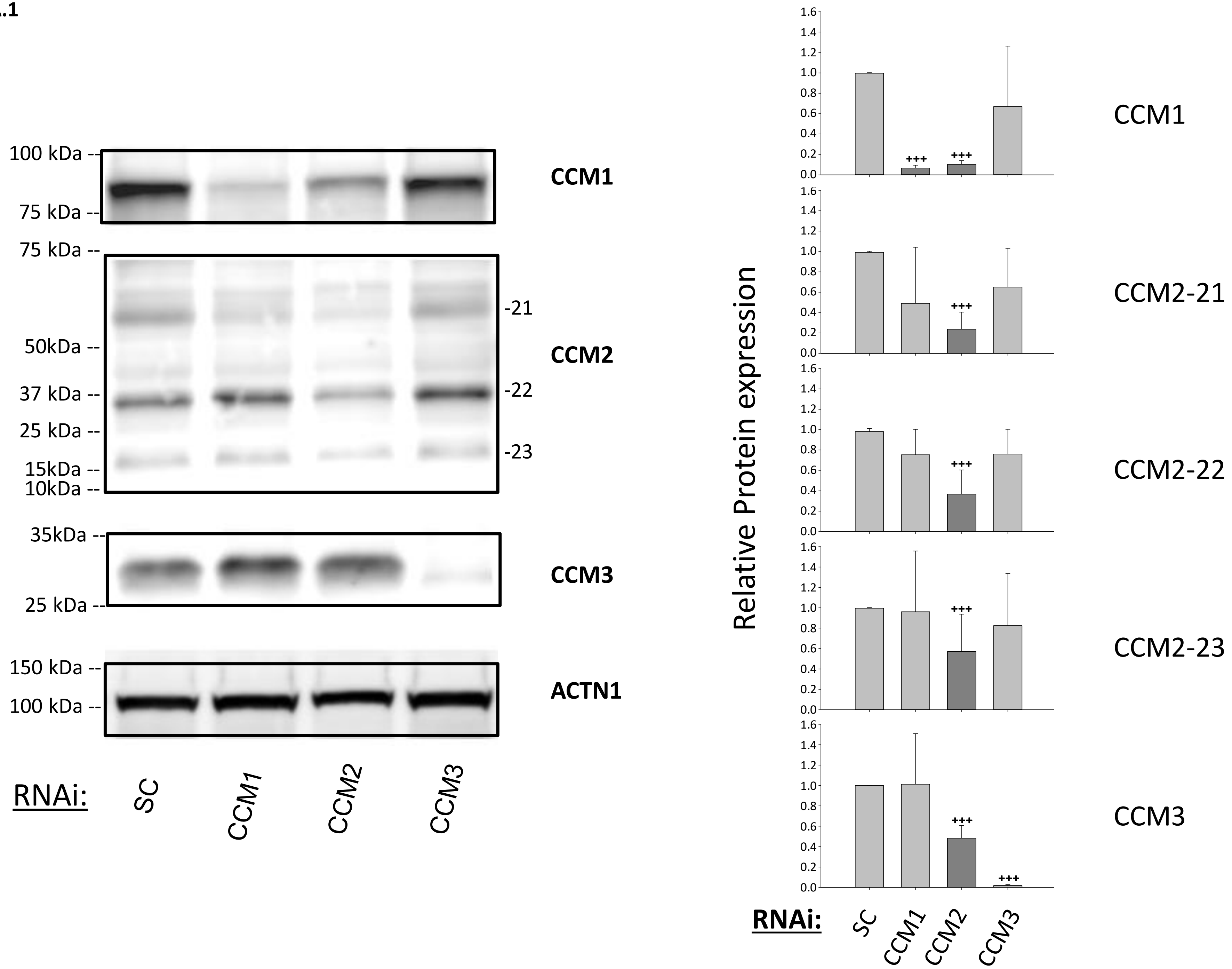

100 kDa
75 kDa
CCM1
75 kDa
50kDa
37 kDa
25 kDa
15kDa
10kDa
-21
-22
-23
CCM2
35kDa
25 kDa
CCM3
150 kDa
100 kDa
ACTN1
RNAi:
SC
CCM1
CCM2
CCM3
Relative Protein expression
CCM1
CCM2-21
CCM2-22
CCM2-23
CCM3
+++
RNAi:
SC
CCM1
CCM2
CCM3

4A.2

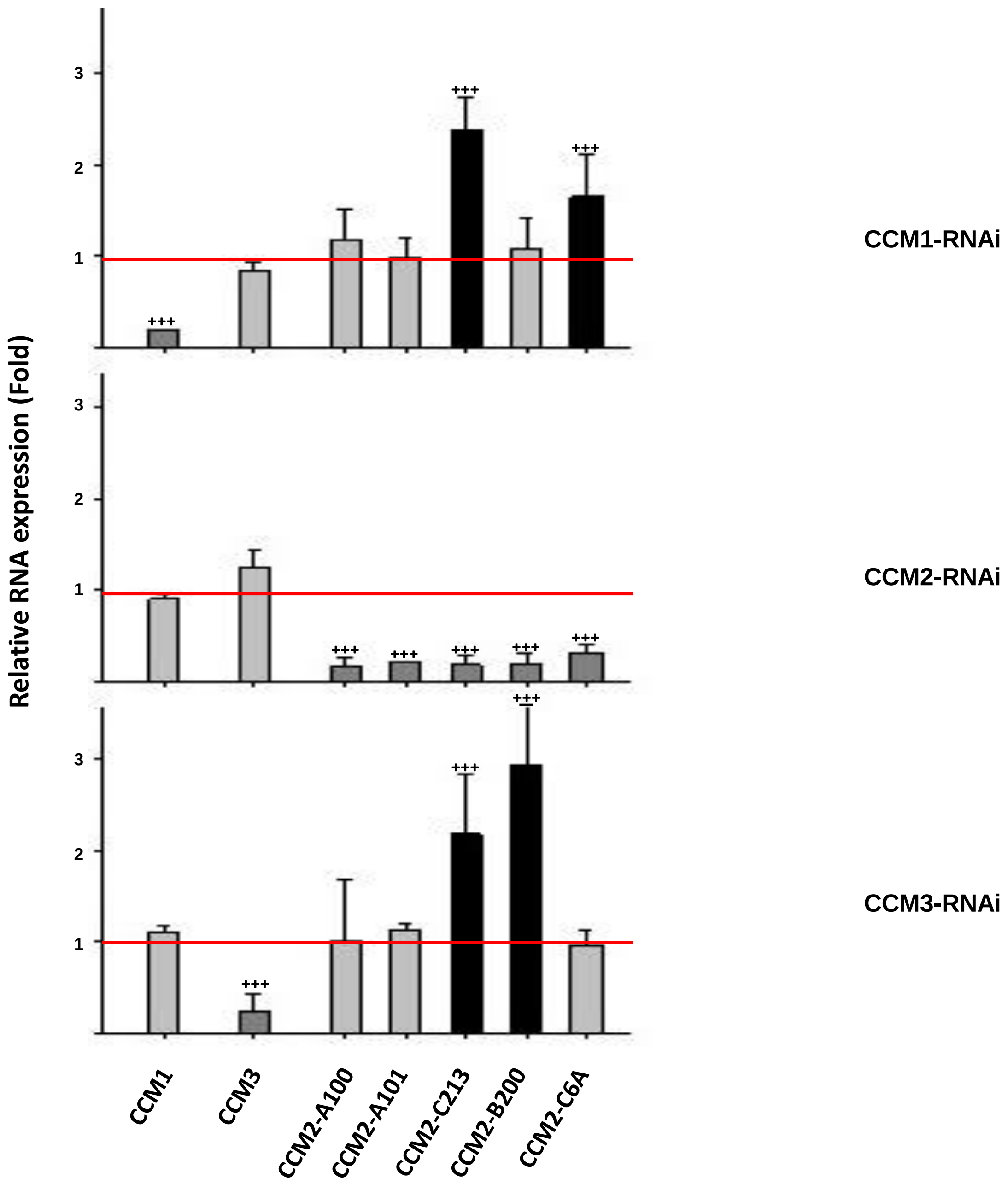
Relative RNA expression (Fold)
3
2
1
+++
CCM1-RNAi
CCM2-RNAi
CCM3-RNAi
CCM1
CCM3
CCM2-A100
CCM2-A101
CCM2-C213
CCM2-B200
CCM2-C6A

**4B.1**

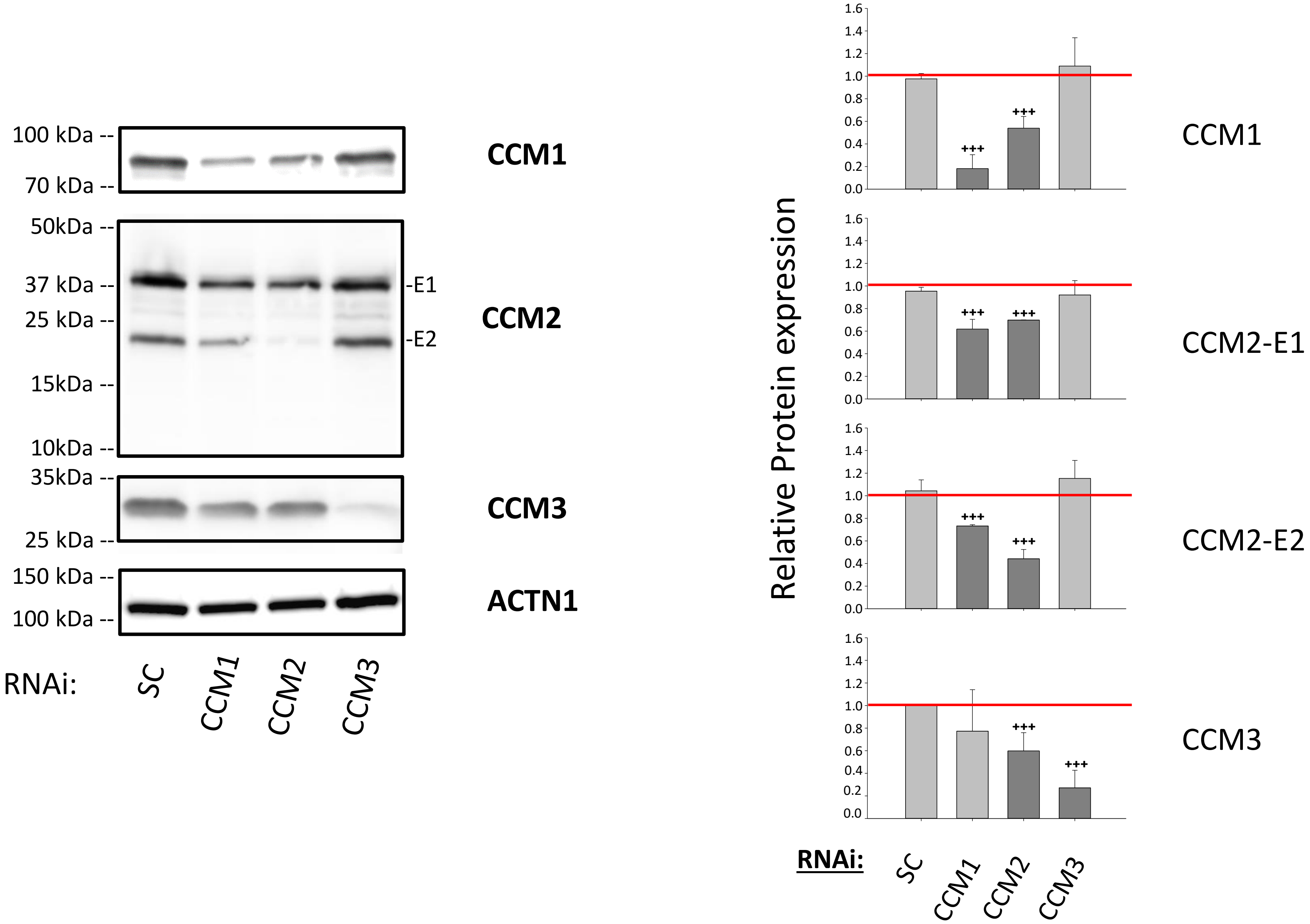

100 kDa
70 kDa
CCM1
50kDa
37 kDa
25 kDa
15kDa
10kDa
-E1
-E2
CCM2
35kDa
25 kDa
CCM3
150 kDa
100 kDa
ACTN1
RNAi:
SC
CCM1
CCM2
CCM3
Relative Protein expression
1.6
1.4
1.2
1.0
0.8
0.6
0.4
0.2
0.0
+++
CCM1
CCM2-E1
CCM2-E2
CCM3
RNAi:
SC
CCM1
CCM2
CCM3

**4B.2**

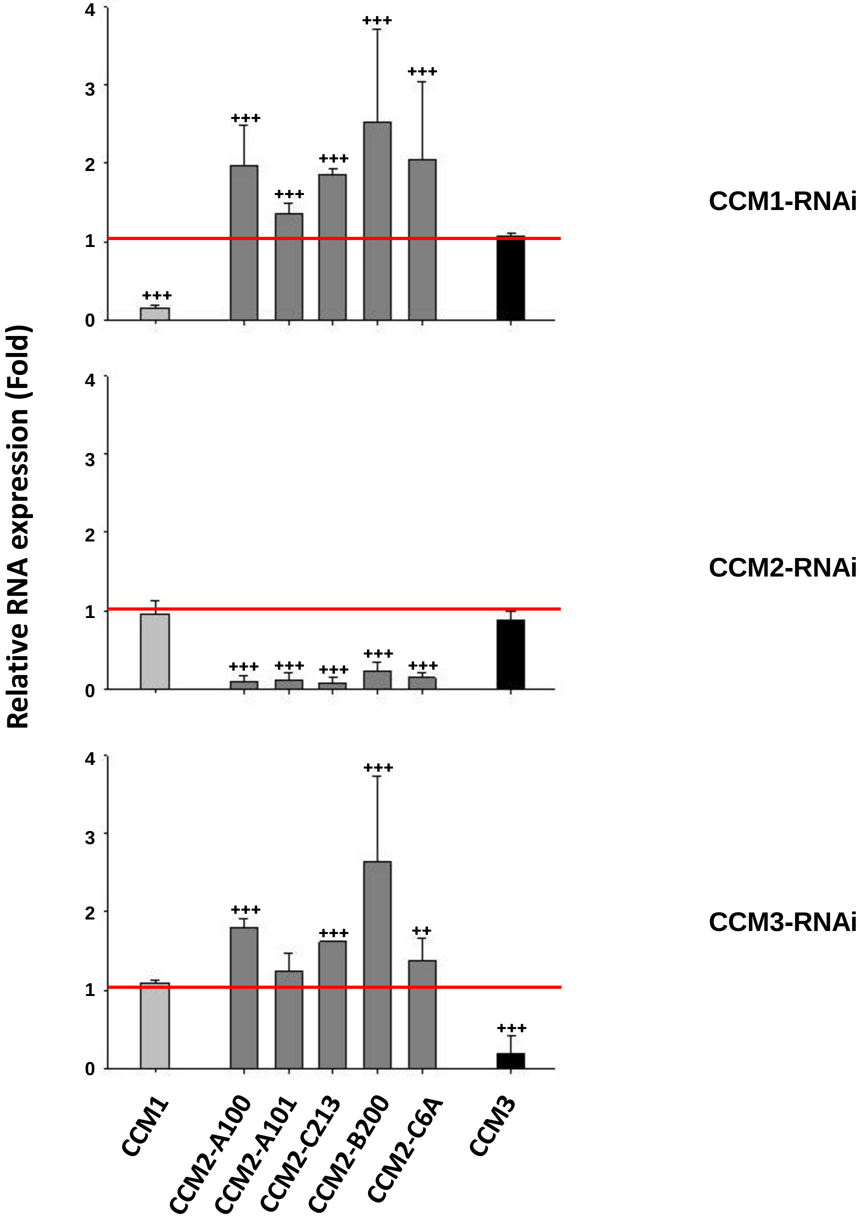

Relative RNA expression (Fold)
CCM1-RNAi
CCM2-RNAi
CCM3-RNAi
0
1
2
3
4
+++
++
CCM1
CCM2-A100
CCM2-A101
CCM2-C213
CCM2-B200
CCM2-C6A
CCM3

**4C.1**

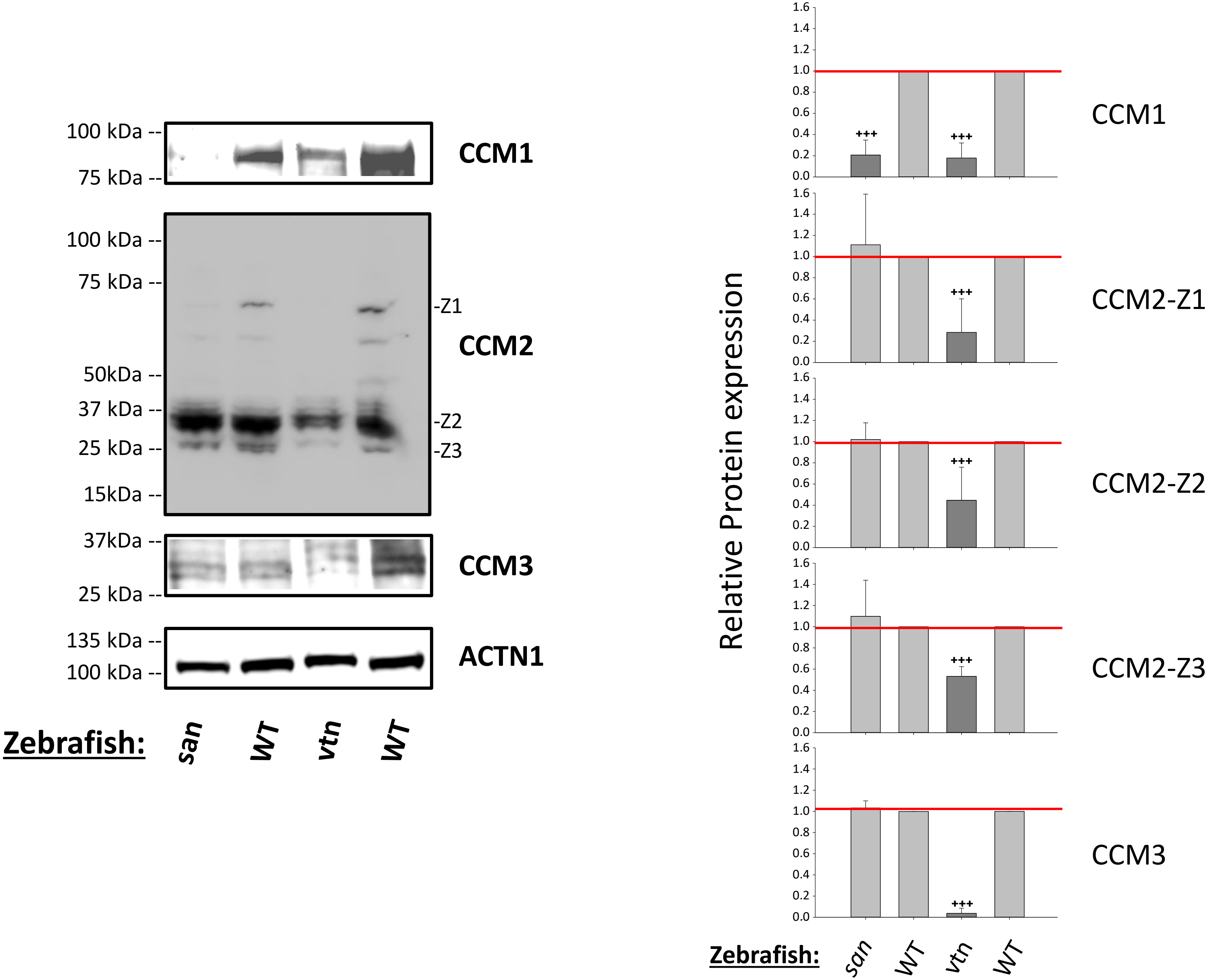

100 kDa
75 kDa
CCM1
100 kDa
75 kDa
50kDa
37 kDa
25 kDa
15kDa
-Z1
-Z2
-Z3
CCM2
37kDa
25 kDa
CCM3
135 kDa
100 kDa
ACTN1
Zebrafish:
san
WT
vtn
WT
Relative Protein expression
+++
CCM1
CCM2-Z1
CCM2-Z2
CCM2-Z3
CCM3
0.0
0.2
0.4
0.6
0.8
1.0
1.2
1.4
1.6
Zebrafish:
san
WT
vtn
WT

4C2

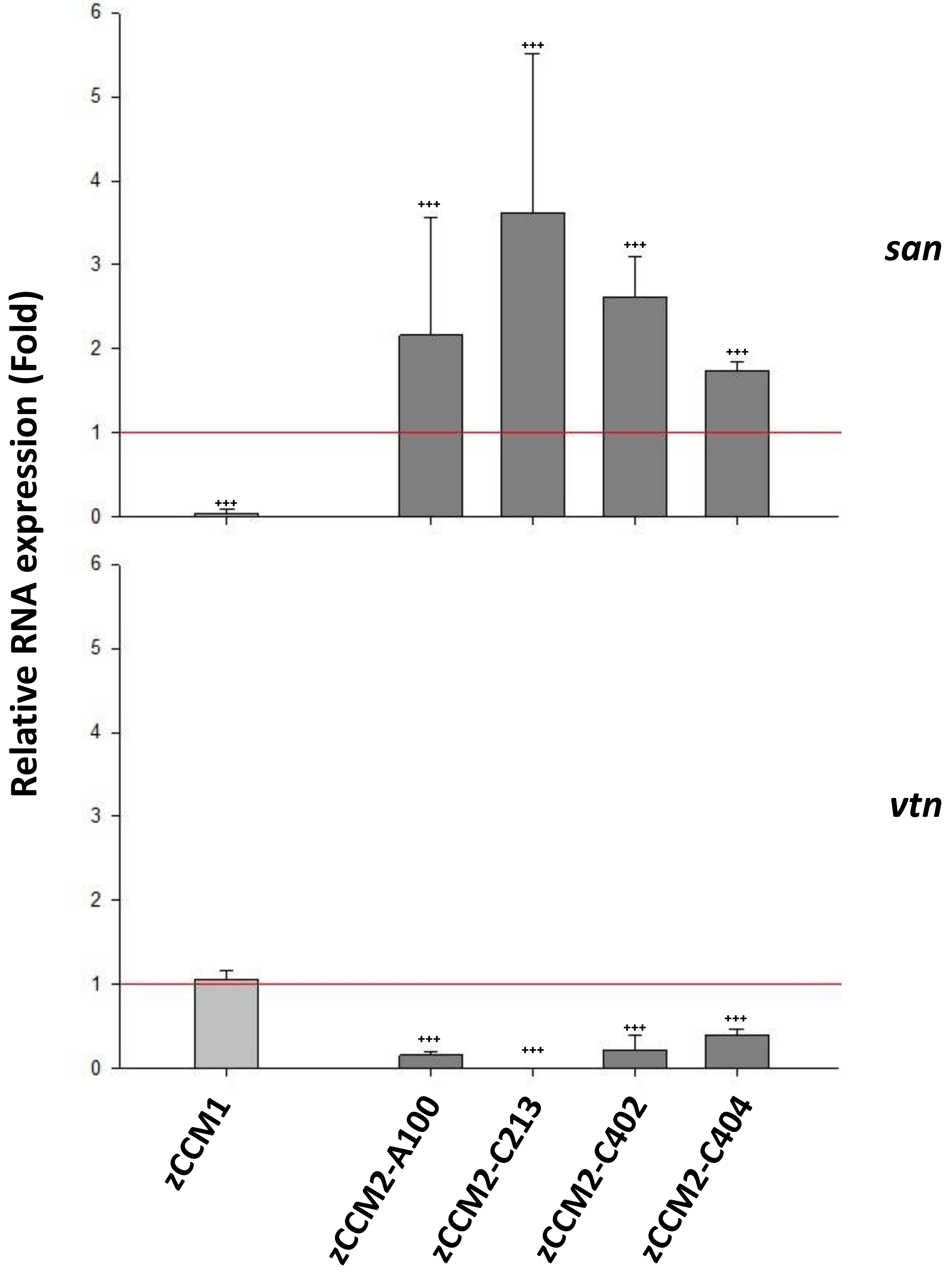
Relative RNA expression (Fold)
6
5
4
3
2
1
0
+++
+++
+++
+++
+++
san
6
5
4
3
2
1
0
+++
+++
+++
+++
vtn
zCCM1
zCCM2-A100
zCCM2-C213
zCCM2-C402
zCCM2-C404

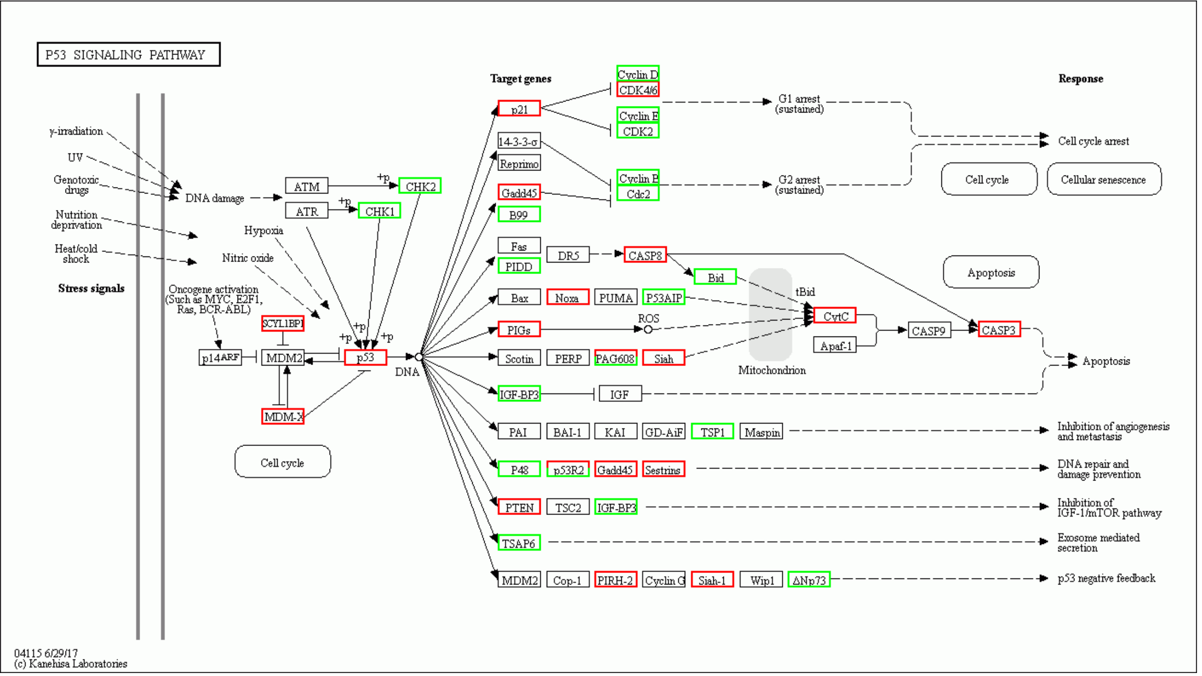
P53 SIGNALING PATHWAY
γ-irradiation
UV
Genotoxic drugs
Nutrition deprivation
Heat/cold shock
Stress signals
DNA damage
Hypoxia
Nitric oxide
Oncogene activation (Such as MYC, E2F1, Ras, BCR-ABL)
ATM
ATR
CHK2
CHK1
SCYL1BP1
p14ARF
MDM2
MDM-X
p53
DNA
Cell cycle
Target genes
p21
14-3-3-σ
Reprimo
Gadd45
B99
Fas
PIDD
DR5
CASP8
Bid
Bax
Noxa
PUMA
P53AIP
PIGs
ROS
Scotin
PERP
PAG608
Siah
IGF-BP3
IGF
PAI
BAI-1
KAI
GD-AiF
TSP1
Maspin
P48
p53R2
Sestrins
PTEN
TSC2
TSAP6
Cop-1
PIRH-2
Cyclin G
Siah-1
Wip1
ΔNp73
Cyclin D
CDK4/6
Cyclin E
CDK2
Cyclin B
Cdc2
G1 arrest (sustained)
G2 arrest (sustained)
tBid
CytC
Apaf-1
CASP9
CASP3
Mitochondrion
Response
Cell cycle arrest
Cell cycle
Cellular senescence
Apoptosis
Inhibition of angiogenesis and metastasis
DNA repair and damage prevention
Inhibition of IGF-1/mTOR pathway
Exosome mediated secretion
p53 negative feedback
04115 6/29/17
(c) Kanehisa Laboratories

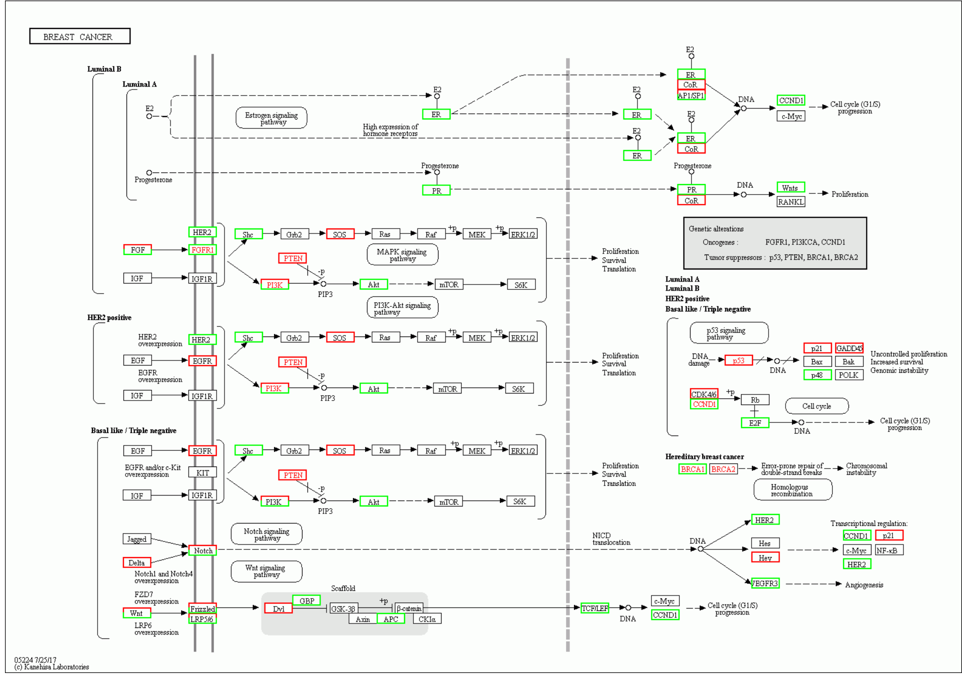
BREAST CANCER
Luminal B
Luminal A
HER2 positive
Basal like / Triple negative
Genetic alterations
Oncogenes : FGFR1, PI3KCA, CCND1
Tumor suppressors : p53, PTEN, BRCA1, BRCA2
Hereditary breast cancer
05224 7/25/17
(c) Kanehisa Laboratories

MIF

MIF + PRG

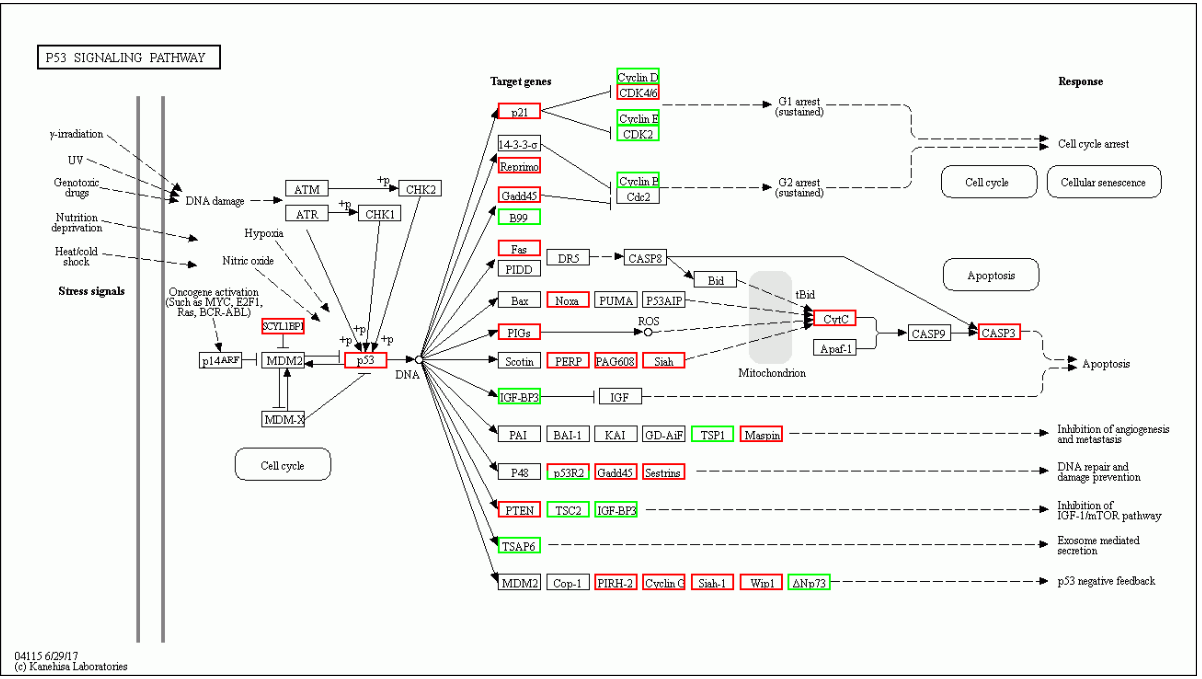
P53 SIGNALING PATHWAY
Stress signals
Target genes
Response
Cell cycle arrest
Cellular senescence
Apoptosis
Inhibition of angiogenesis and metastasis
DNA repair and damage prevention
Inhibition of IGF-1/mTOR pathway
Exosome mediated secretion
p53 negative feedback
04115 6/29/17
(c) Kanehisa Laboratories

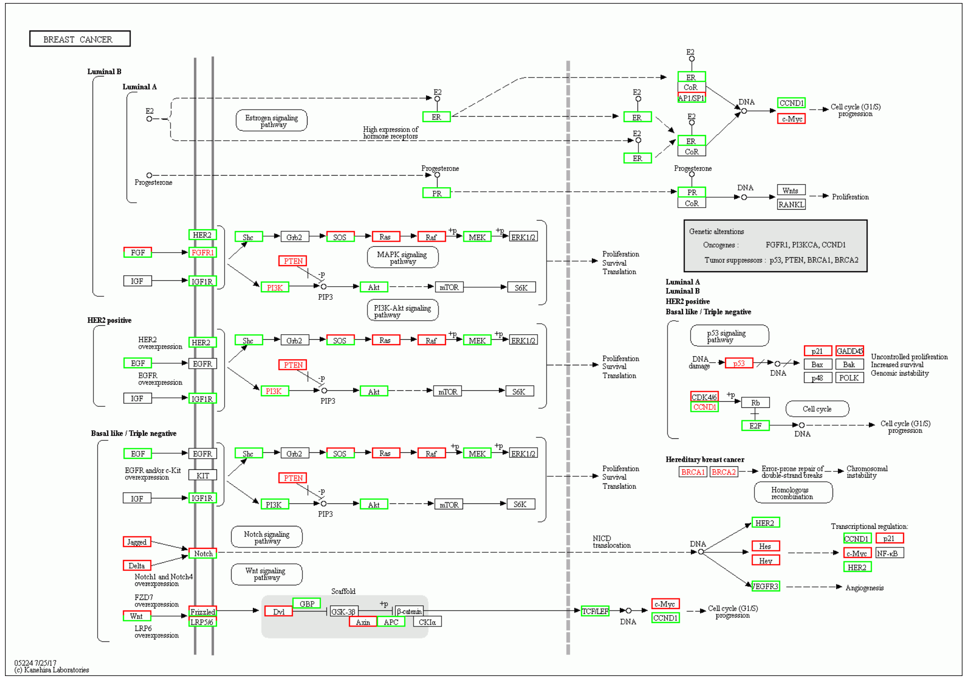
BREAST CANCER
Luminal B
Luminal A
HER2 positive
Basal like / Triple negative
Genetic alterations
Oncogenes : FGFR1, PI3KCA, CCND1
Tumor suppressors : p53, PTEN, BRCA1, BRCA2
Hereditary breast cancer
05224 7/25/17
(c) Kanehisa Laboratories

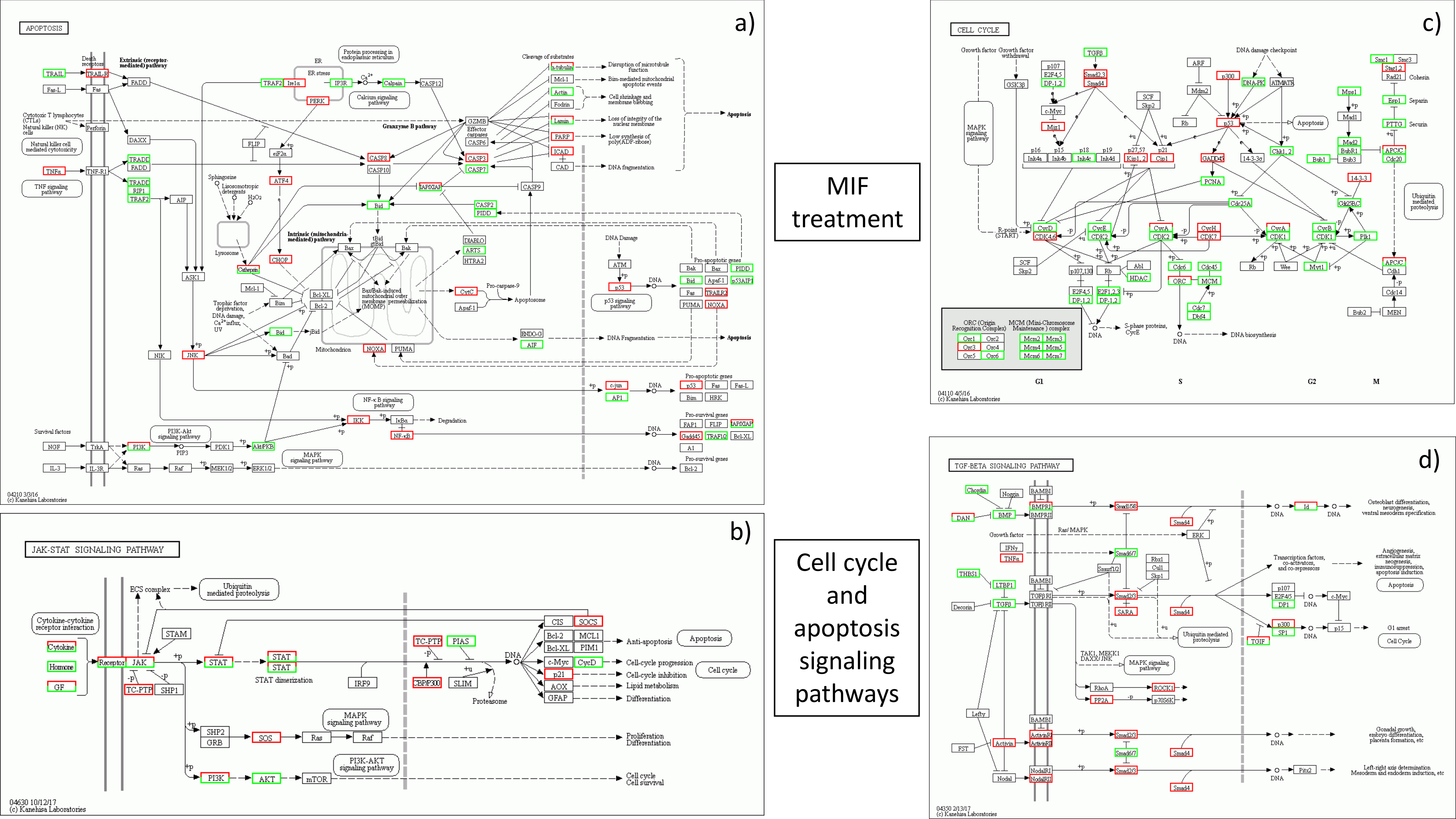

a)
APOPTOSIS
Extrinsic (receptor-mediated) pathway
Intrinsic (mitochondria-mediated) pathway
Granzyme B pathway
Apoptosis
Pro-apoptotic genes
Pro-survival genes
04210 3/3/16
(c) Kanehisa Laboratories
b)
JAK-STAT SIGNALING PATHWAY
Cytokine-cytokine receptor interaction
Cytokine
Hormone
GF
Receptor
JAK
STAT
STAT dimerization
ECS complex
Ubiquitin mediated proteolysis
STAM
TC-PTP
SHP1
IRF9
PIAS
CBP/P300
SLIM
Proteasome
DNA
CIS
SOCS
Bcl-2
MCL1
Bcl-XL
PIM1
c-Myc
CycD
p21
AOX
GFAP
Anti-apoptosis
Apoptosis
Cell-cycle progression
Cell-cycle inhibition
Lipid metabolism
Differentiation
Cell cycle
SHP2
GRB
SOS
Ras
Raf
MAPK signaling pathway
Proliferation
Differentiation
PI3K
AKT
mTOR
PI3K-AKT signaling pathway
Cell cycle
Cell survival
04630 10/12/17
(c) Kanehisa Laboratories
MIF treatment
Cell cycle and apoptosis signaling pathways
c)
CELL CYCLE
G1
S
G2
M
DNA biosynthesis
04110 4/5/16
(c) Kanehisa Laboratories
d)
TGF-BETA SIGNALING PATHWAY
04350 2/13/17
(c) Kanehisa Laboratories

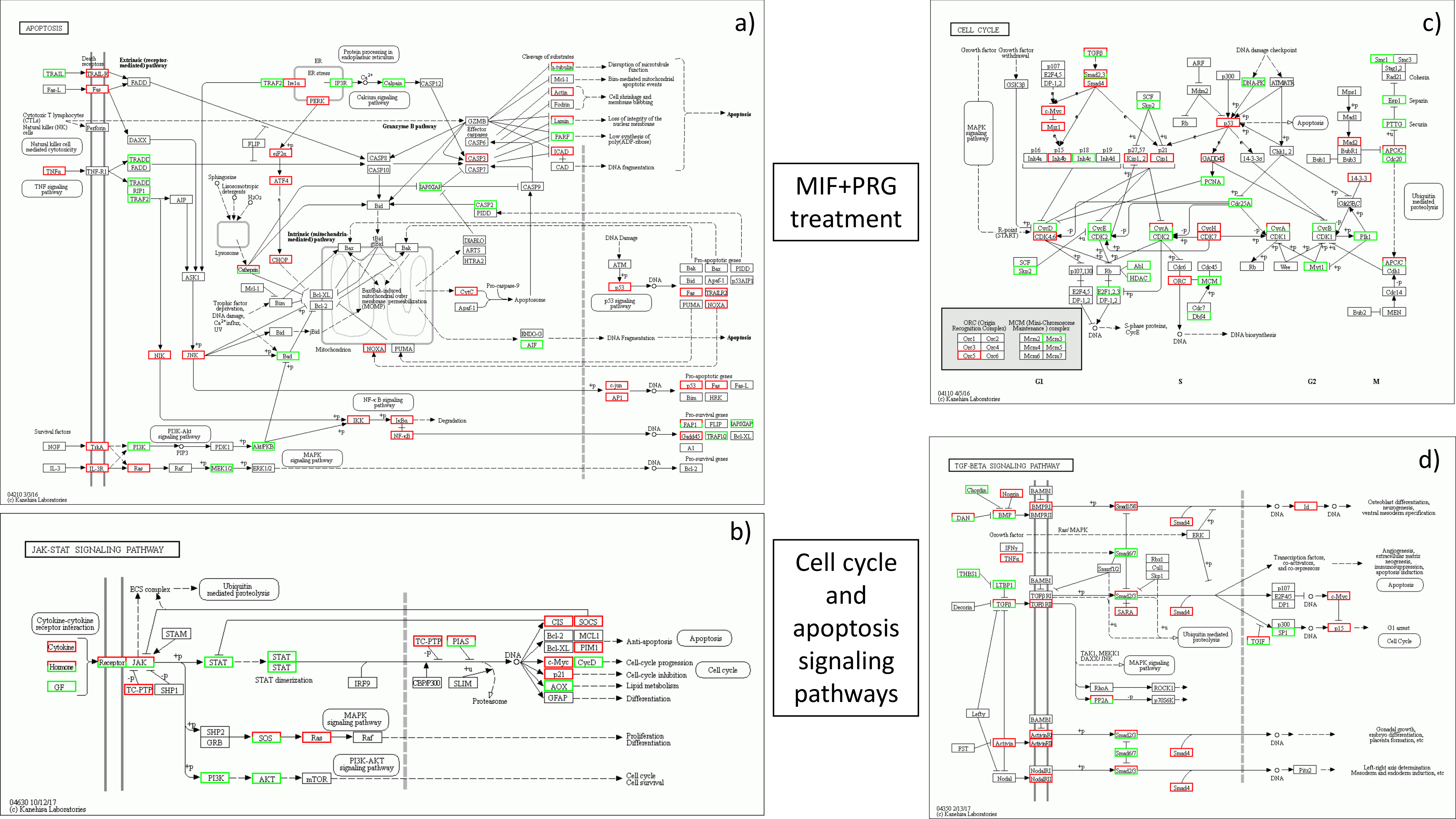
a)
APOPTOSIS
b)
JAK-STAT SIGNALING PATHWAY
MIF+PRG treatment
Cell cycle and apoptosis signaling pathways
c)
CELL CYCLE
d)
TGF-BETA SIGNALING PATHWAY

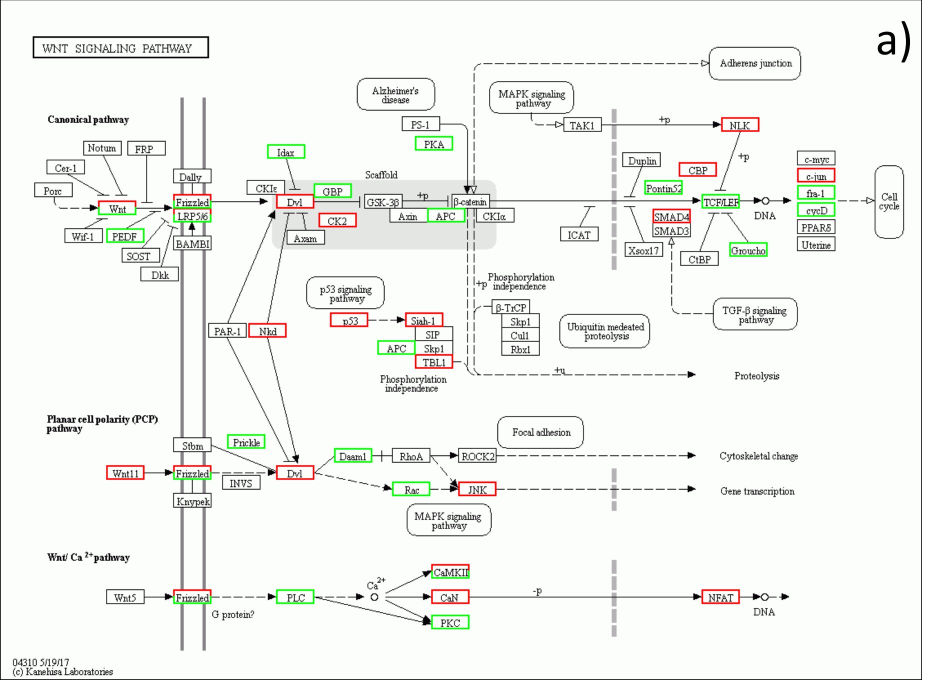


MIF treatment

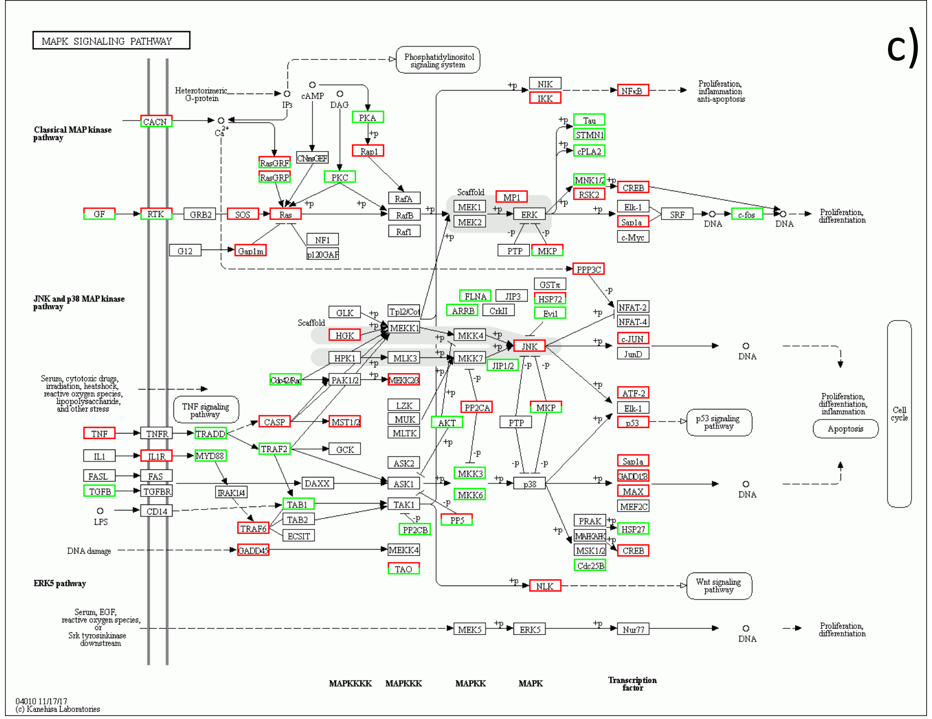


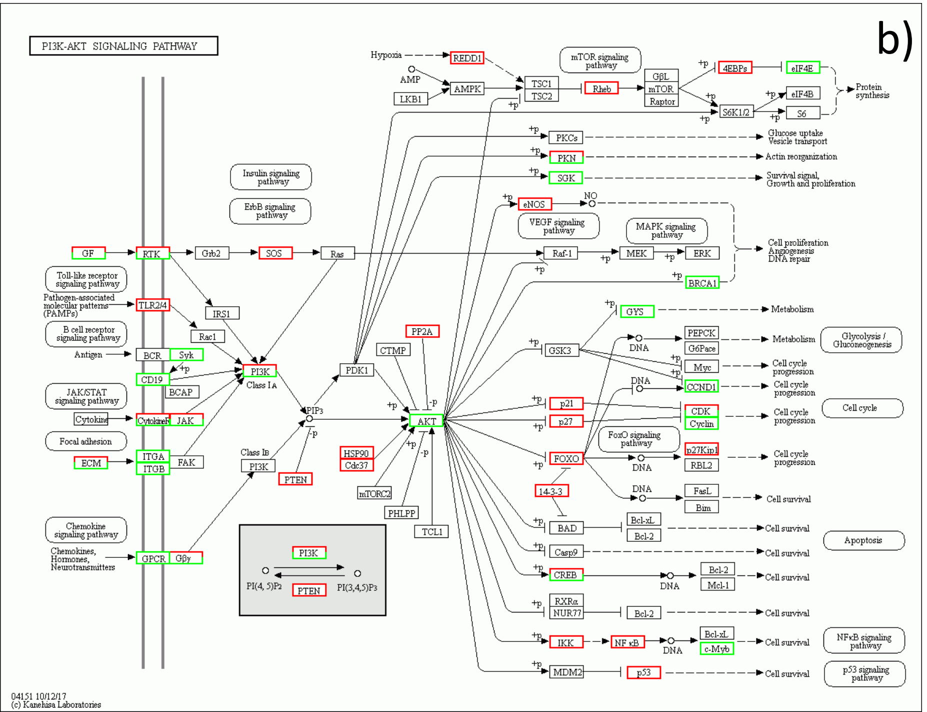


WNT/MAPK RAS and P13K-AKT signaling pathways

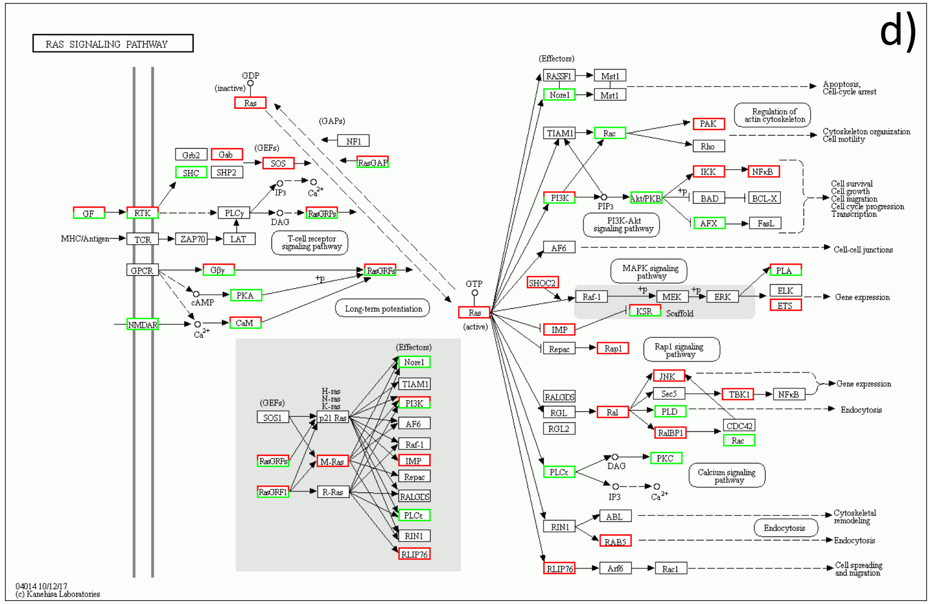

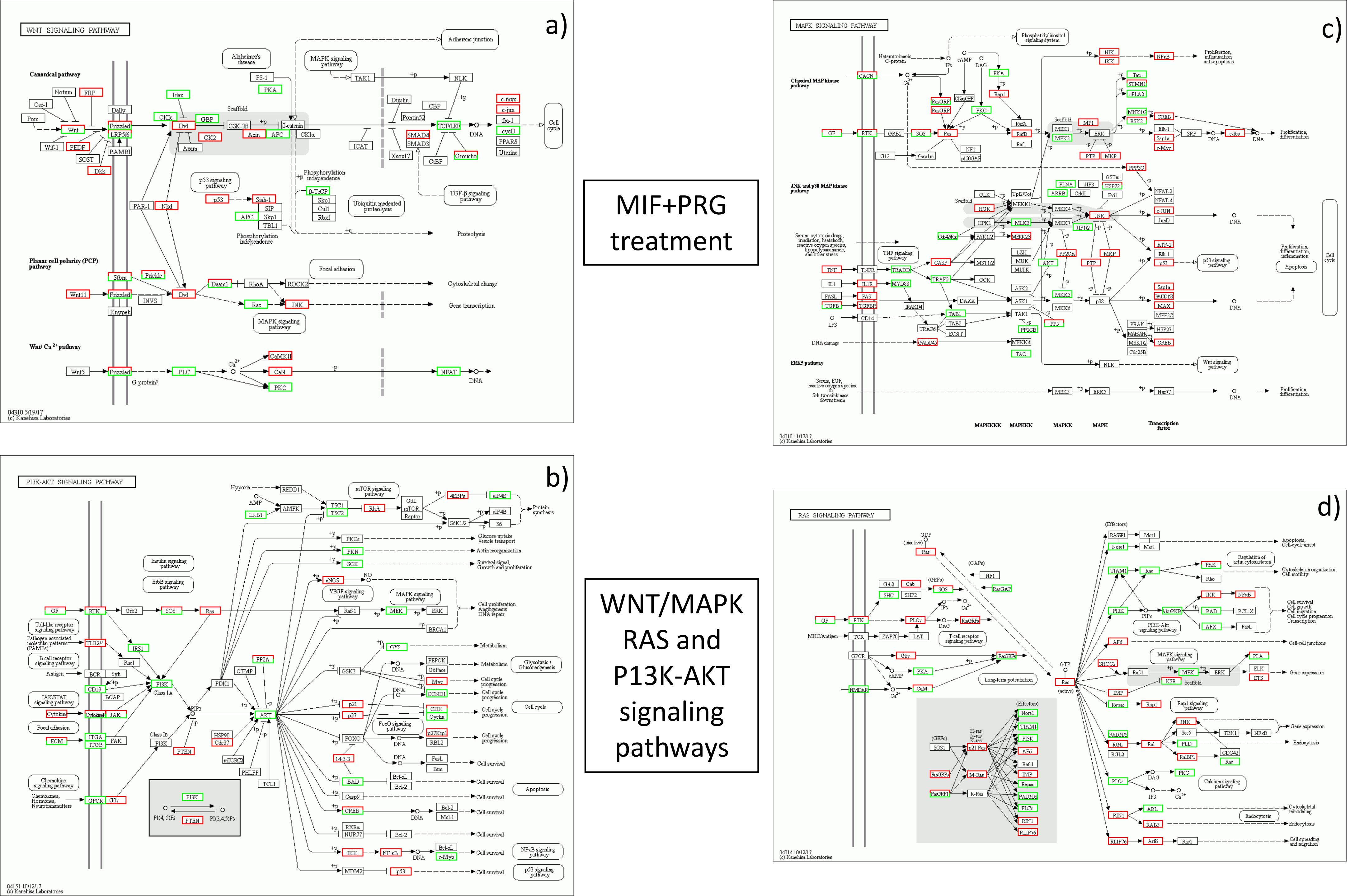

WNT SIGNALING PATHWAY
a)
Canonical pathway
Planar cell polarity (PCP) pathway
Wnt/ Ca 2+ pathway
04310 5/19/17
(c) Kanehisa Laboratories
PI3K-AKT SIGNALING PATHWAY
b)
04151 10/12/17
(c) Kanehisa Laboratories
MIF+PRG treatment
WNT/MAPK RAS and P13K-AKT signaling pathways
MAPK SIGNALING PATHWAY
c)
Classical MAP kinase pathway
JNK and p38 MAP kinase pathway
ERK5 pathway
MAPKKKK
MAPKKK
MAPKK
MAPK
Transcription factor
04010 11/17/17
(c) Kanehisa Laboratories
RAS SIGNALING PATHWAY
d)
04014 10/12/17
(c) Kanehisa Laboratories

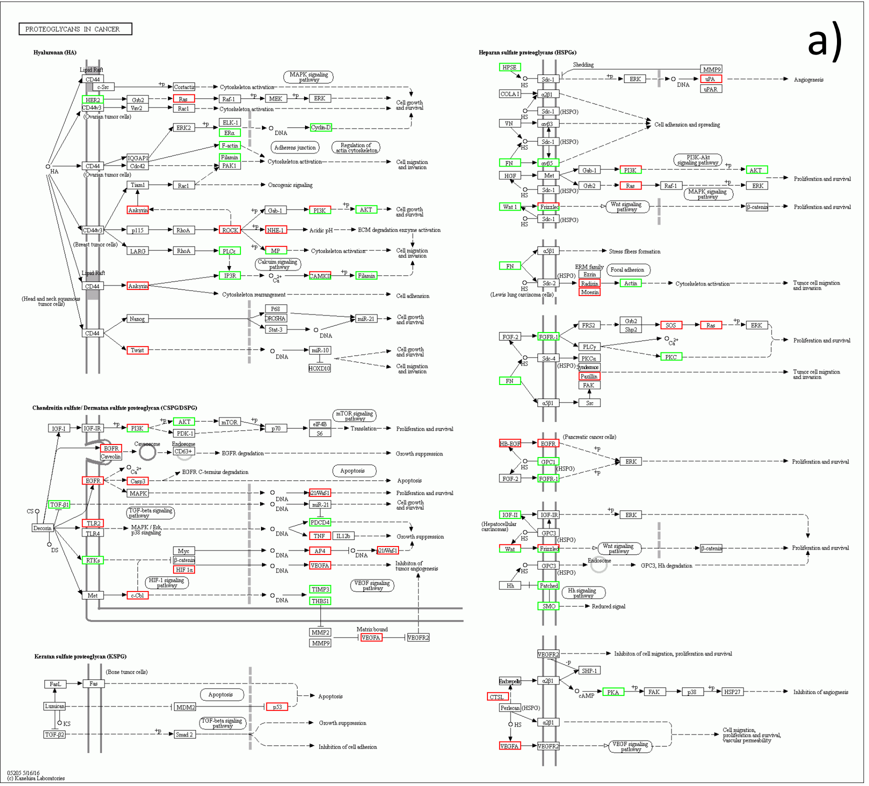


MIF treatment

Cancer Pathways

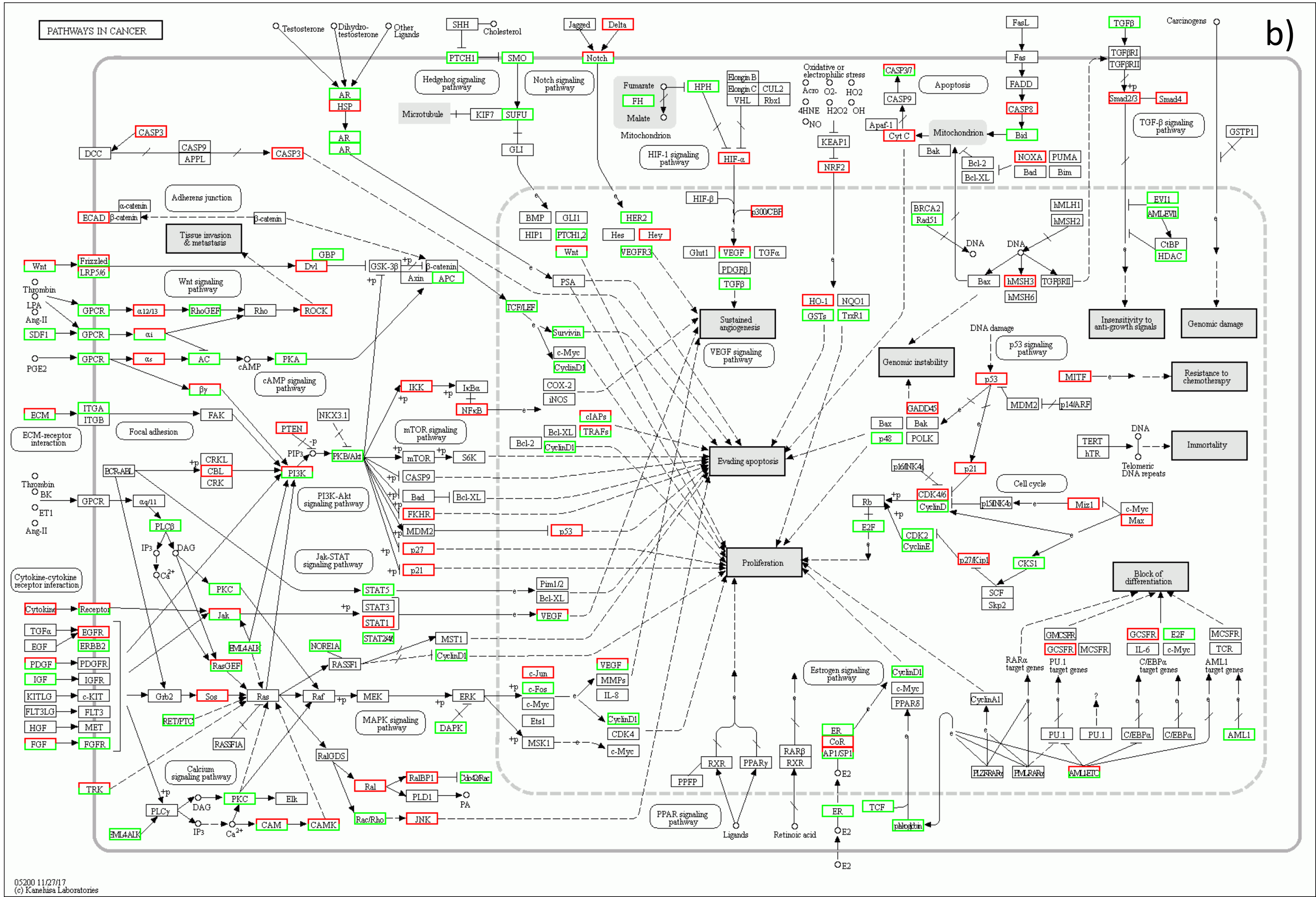

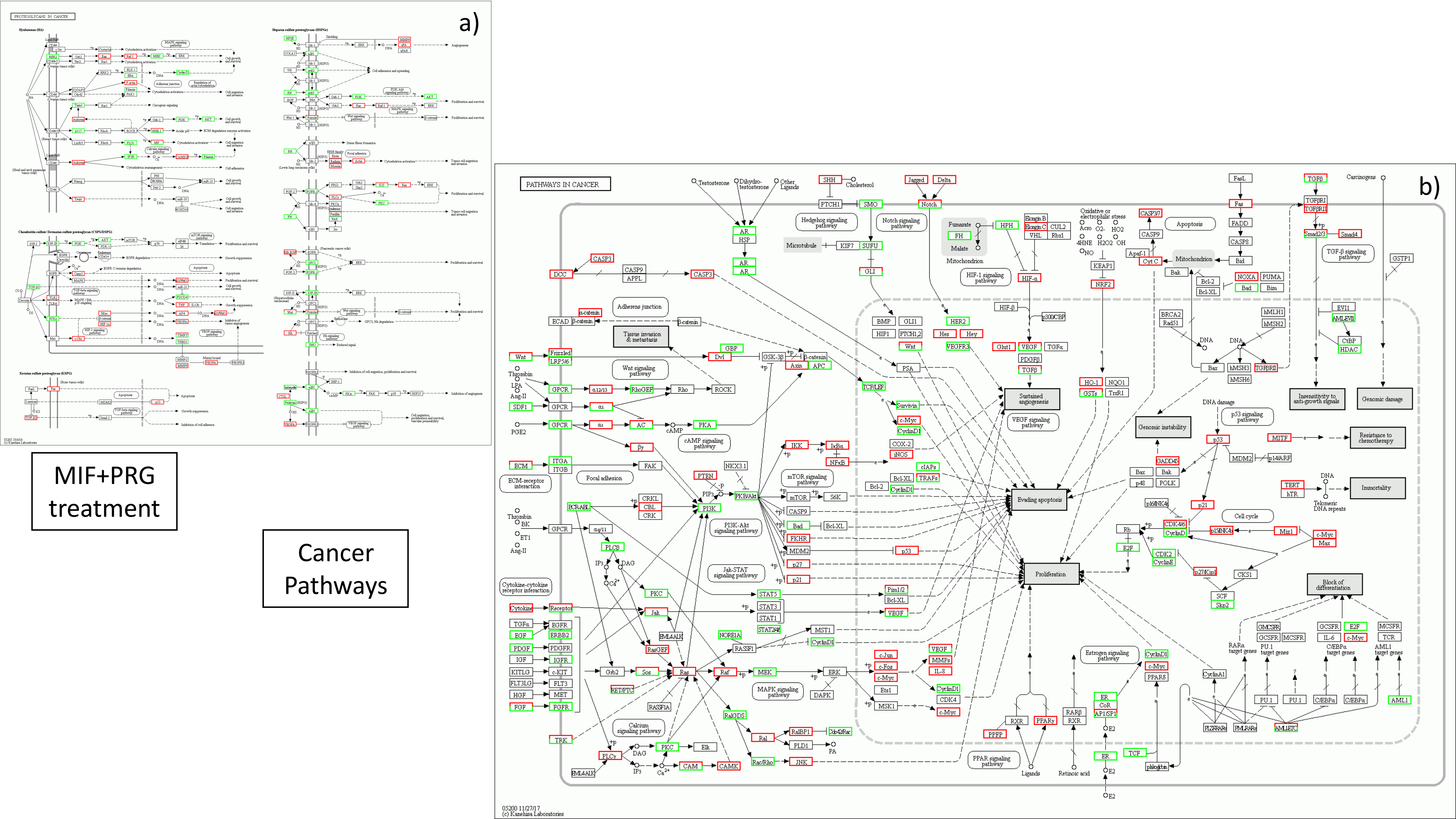

a)
PROTEOGLYCANS IN CANCER
Hyaluronan (HA)
Heparan sulfate proteoglycans (HSPG)
Chondroitin sulfate / Dermatan sulfate proteoglycan (CSPG/DSPG)
Keratan sulfate proteoglycan (KSPG)
MIF+PRG treatment
Cancer Pathways
b)
PATHWAYS IN CANCER
Tissue invasion & metastasis
Sustained angiogenesis
Evading apoptosis
Proliferation
Genomic instability
Insensitivity to anti-growth signals
Genomic damage
Resistance to chemotherapy
Immortality
Block of differentiation
05200 11/27/17
(c) Kanehisa Laboratories

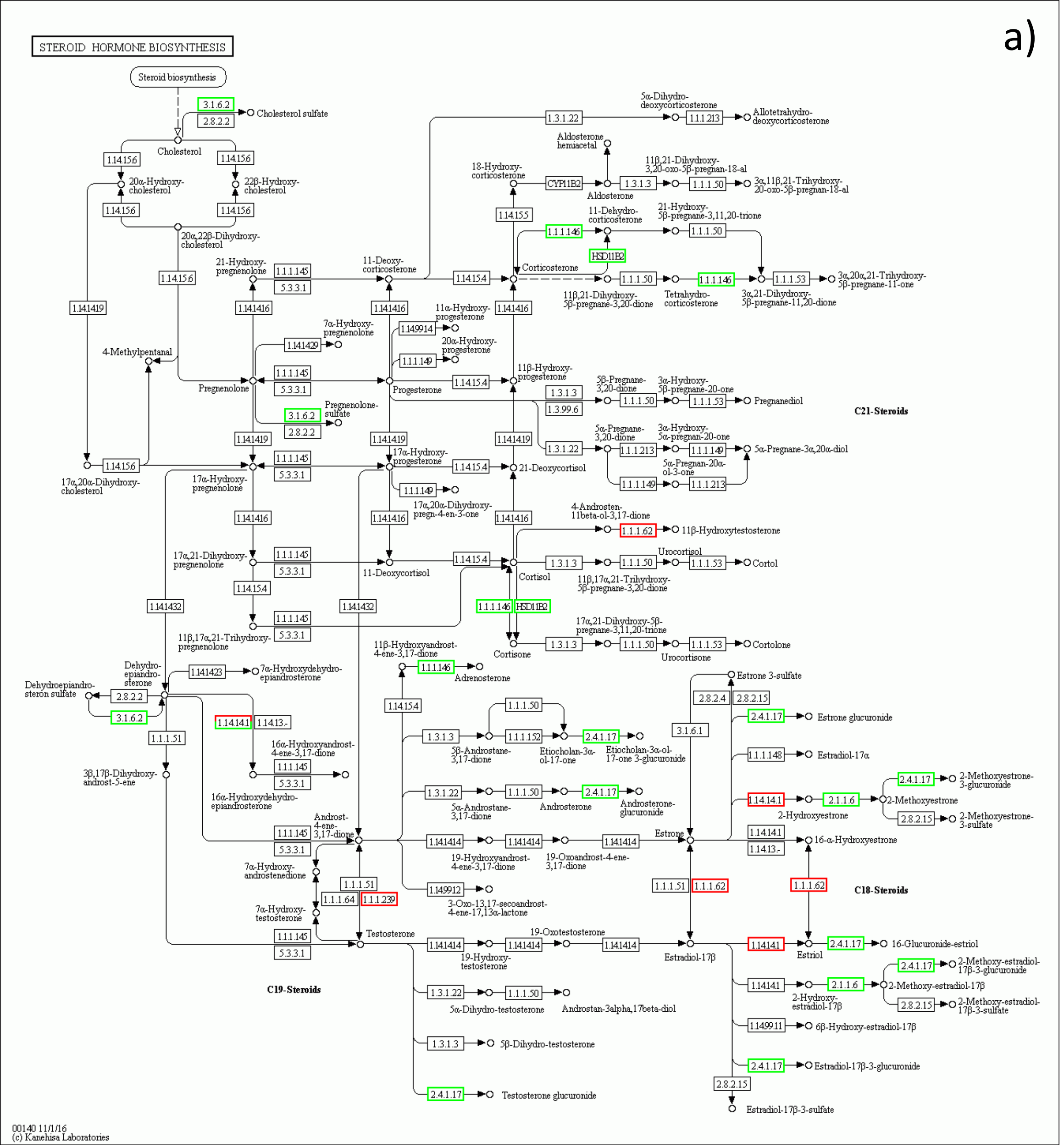


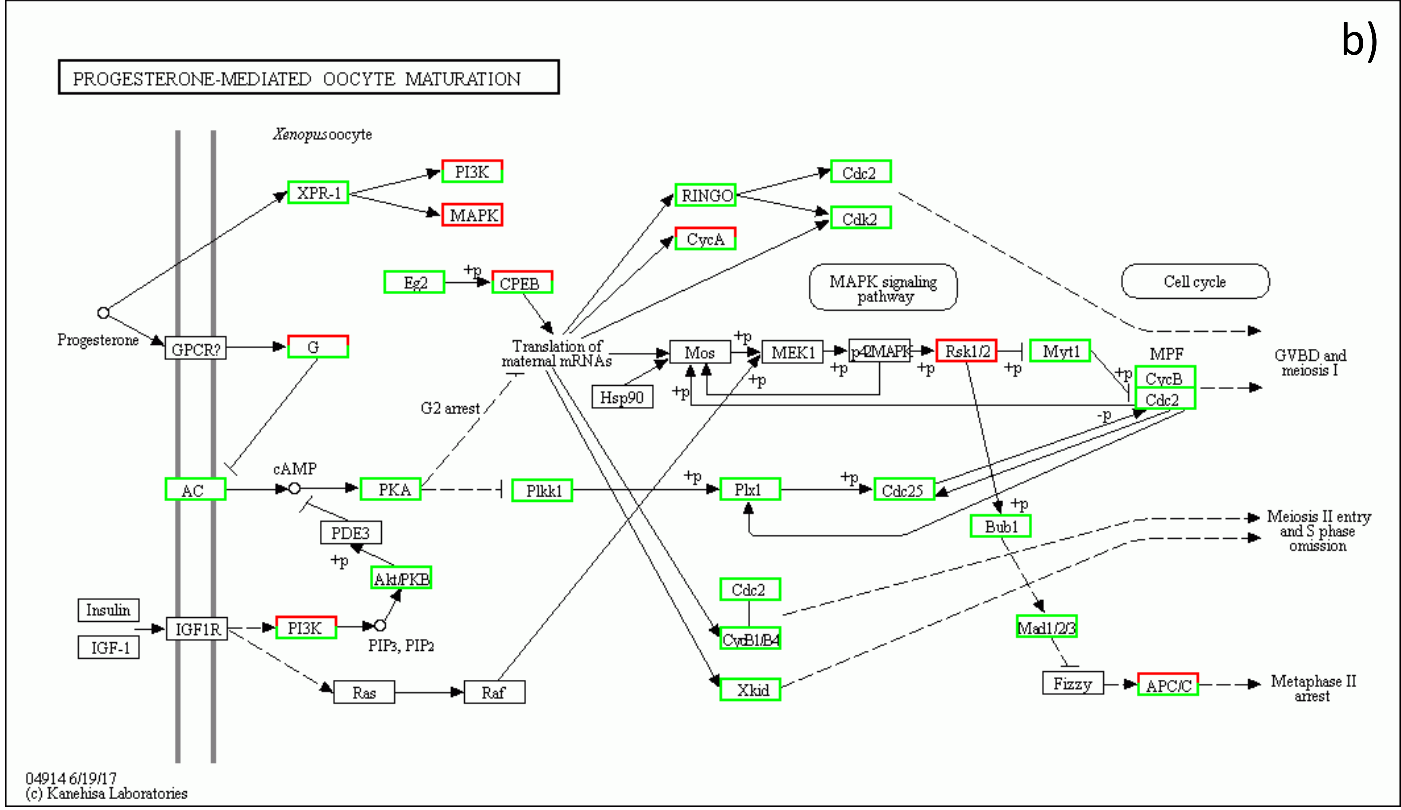


MIF treatment

Hormone signaling pathways

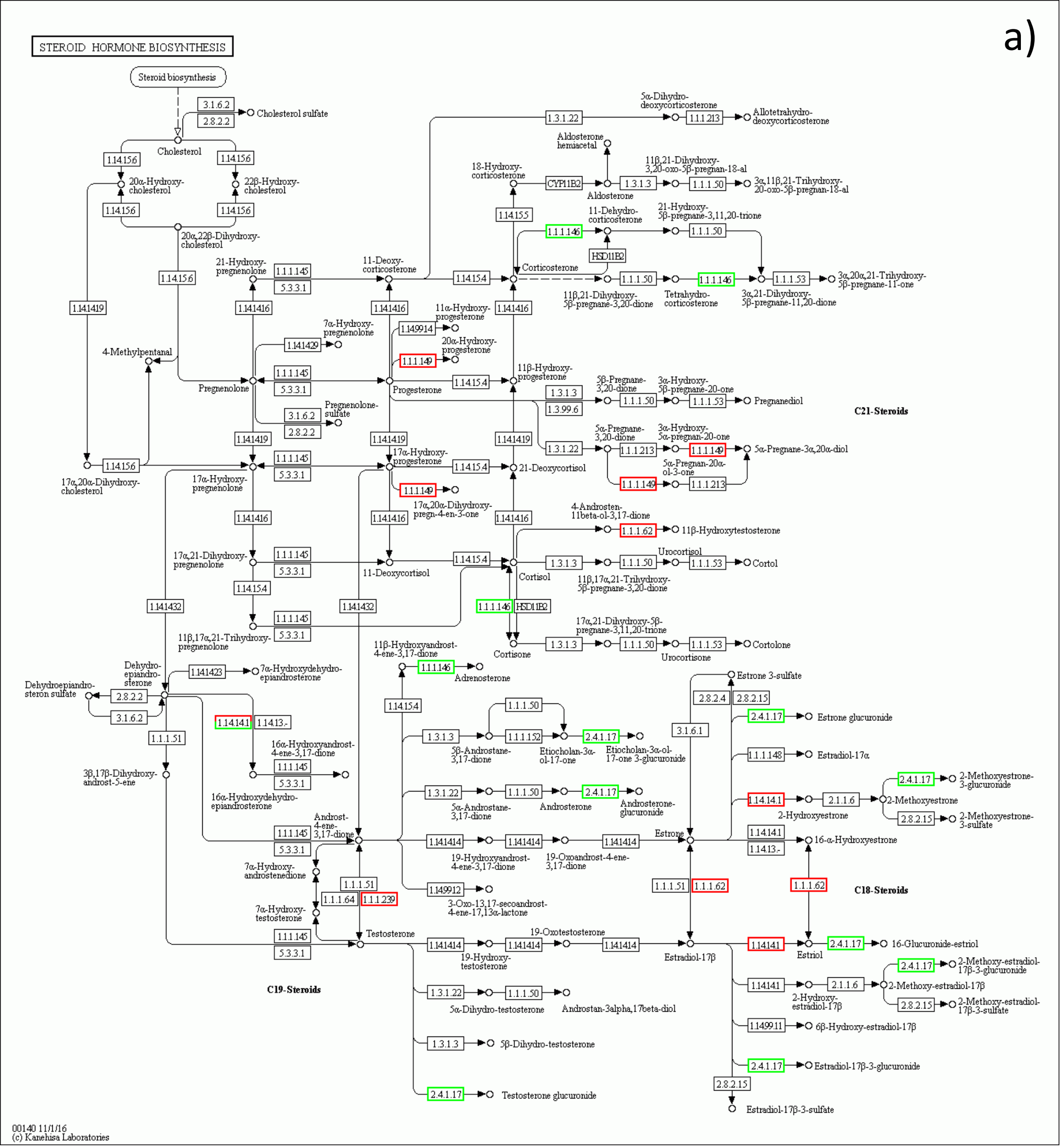


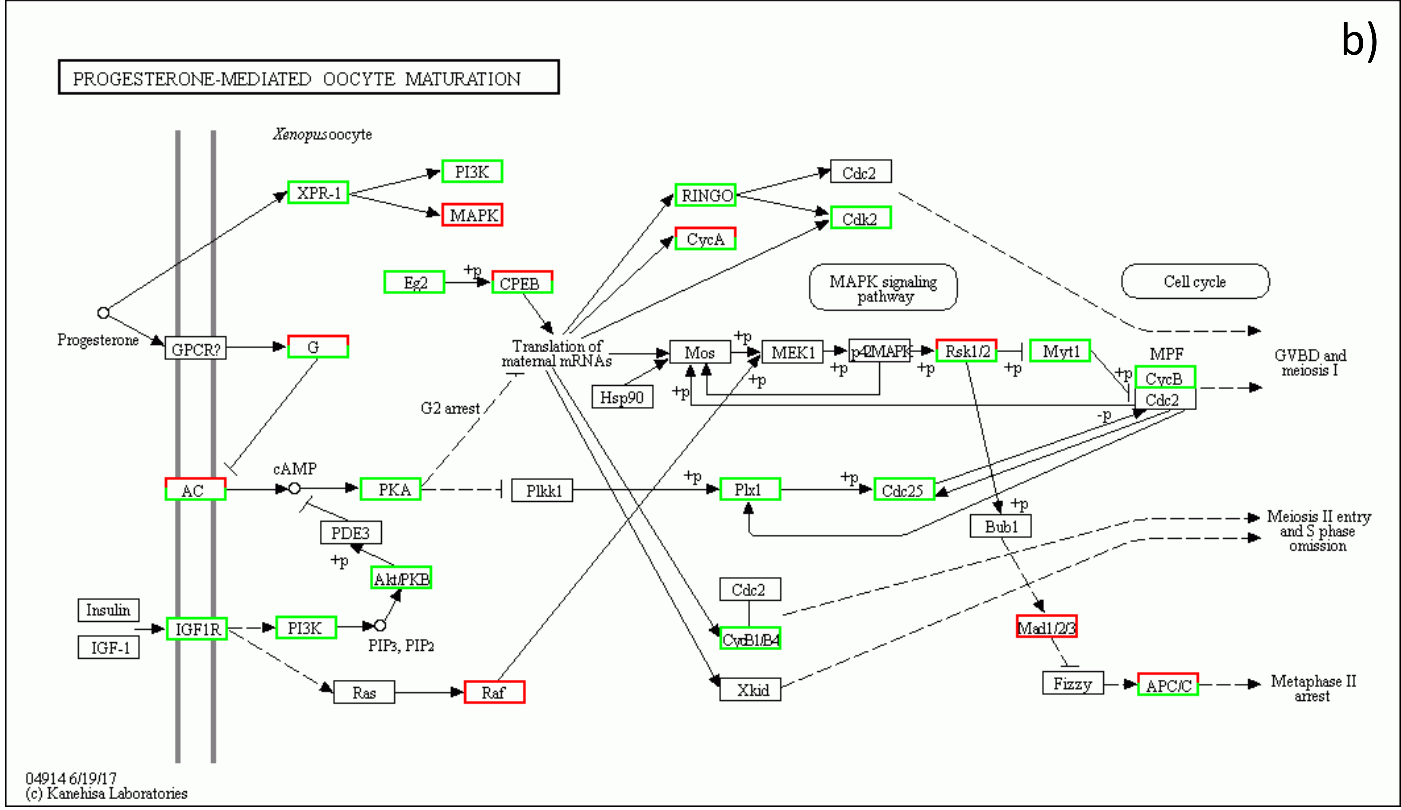


MIF+PRG treatment

Hormone signaling pathways

**Table S1**

| Application | Antigen | Gene | Clone | Secondary | Vendor | Cat # |
|---|---|---|---|---|---|---|
| IF/WB | CCM1 | Krit1 | E-8 | (anti-mouse) | Santa Cruz | sc-514371 |
| IF/WB | CCM1-AF488 | Krit1 | E-8 | (anti-mouse) | Santa Cruz | sc-514371 |
| IF/WB | CCM1 | Krit1 | 8-RY2 | (anti-mouse) | Santa Cruz | sc-134376 |
| IF/WB | CCM1 | Krit1 | | (anti-rabbit) | OriGene | AP26021PU-L |
| IHC/DAB/WB | CCM2 | MGC4607 | | (anti-rabbit) | Novus | NBP1-86730 |
| IHC/DAB/WB | CCM2 | MGC4607 | | (anti-rabbit) | Novus | NBP215761 |
| IHC/DAB/WB | CCM2 | MGC4607 | | (anti-goat) | Sigma | SAB2500214 |
| IF/WB | CCM3 | PDCD10 | C-8 | (anti-mouse) | Santa Cruz | sc-365586 |
| IF/WB | CCM3-AF647 | PDCD10 | C-8 | (anti-mouse) | Santa Cruz | sc-365586 |
| IF/WB | CCM3 | PDCD10 | F-12 | (anti-mouse) | Santa Cruz | sc-365587 |
| WB | PAQR5 | mPRγ/PAQR5 | | (Anti-Rabbit) | Aviva | OASG04642 |
| WB | PAQR6 | mPRδ/PAQR6 | | (Anti-Rabbit) | Aviva | ARP49900 |
| WB | PAQR6 | mPRδ/PAQR6 | | (Anti-Rabbit) | Aviva | ARP49901 |
| WB | PAQR5/6 | mPRδ/γ | B-8 | (anti-mouse) | Santa Cruz | sc-514273 |
| WB | PAQR7 | mPRα/PAQR7 | | (Anti-Rabbit) | Aviva | OASG04641 |
| WB | PAQR7 | mPRα/PAQR7 | | (Anti-Rabbit) | Aviva | ARP67727 |
| WB | PAQR8 | mPRβ/PAQR8 | | (Anti-Rabbit) | Aviva | ARP66903 |
| WB | PAQR9 | mPRε/PAQR9 | | (Anti-Rabbit) | Aviva | ARP62890 |
| WB | PGRMC1 | | C-4 | (anti-mouse) | Santa Cruz | sc-393015 |
| WB | PR | | AB52 | (anti-mouse) | Santa Cruz | sc-810 |
| WB | PR | | F-4 | (anti-mouse) | Santa Cruz | sc-166169 |
| WB | AR | | 441 | (anti-mouse) | Santa Cruz | sc-7305 |
| WB | GR | | FiGR | (anti-mouse) | Santa Cruz | sc-12763 |
| WB | GR | | 3D5 | (anti-mouse) | Santa Cruz | sc-56851 |
| WB | β-actin | | C-2 | (anti-mouse) | Santa Cruz | sc-8432 |
| WB | α-actinin | | H-2 | (anti-mouse) | Santa Cruz | sc-17829 |
| WB | α-actinin-AF488 | | H-2 | | Santa Cruz | sc-17829 |
| WB | Rabbit-HRP | | | | Santa Cruz | sc-2357 |
| WB | Mouse-HRP | | m-IgGκ BP | | Santa Cruz | sc-516102 |
| WB | m-IgGκ BP IgG-CFL 488 | | m-IgGκ BP | | Santa Cruz | sc-516176 |
| WB | Rabbit IgG-CFL 488 | | | | Santa Cruz | sc-516248 |
| IHC/IF | DAPI | | | | Santa Cruz | sc-3598 |
| IHC/IF | phalloidin-AF555 | | | | Santa Cruz | sc-2363794 |

**Table S2**

| Application | siRNAs | Target | Vendor | Cat # |
|---|---|---|---|---|
| RNAi | ON-TARGETplus Human KRIT1 siRNA - SMARTpool | Krit1 | Dharmacon | L-003825-00-0010 |
| RNAi | ON-TARGETplus Human CCM2 siRNA - SMARTpool | MGC4607 | Dharmacon | L-014728-01-0010 |
| RNAi | ON-TARGETplus Human CCM2l siRNA - SMARTpool | MGC4607l | Dharmacon | L-018652-02-0010 |
| RNAi | ON-TARGETplus Human PDCD10 siRNA - SMARTpool | PDCD10 | Dharmacon | L-004436-00-0010 |
| RNAi | KRIT1 (Human) - 3 unique 27mer siRNA duplexes | Krit1 | OriGene | SR313185 |
| RNAi | CCM2 (Human) - 3 unique 27mer siRNA duplexes | MGC4607 | OriGene | SAB2500214 |
| RNAi | PDCD10 (Human) - 3 unique 27mer siRNA duplexes | PDCD10 | OriGene | sc-365586 |
| RNAi | Ccm2 Rat siRNA Oligo Duplex (Locus ID 305505) | MGC4607 | OriGene | SR509691 |
| RNAi | ON-TARGETplus Human PAQR7 (164091) siRNA - SMARTpool | PAQR7 | Dharmacon | L-008033-00- 0005 |
| RNAi | ON-TARGETplus Human PAQR8 (85315) siRNA - SMARTpool | PAQR8 | Dharmacon | L-007820-00- 0005 |
| RNAi | ON-TARGETplus Human PAQR5 (54852) siRNA - SMARTpool | PAQR5 | Dharmacon | L-008034-00- 0005 |
| RNAi | ON-TARGETplus Human PAQR6 (79957) siRNA - SMARTpool | PAQR6 | Dharmacon | L-008054-020005 |
| RNAi | ON-TARGETplus Human PGRMC1 (10857) siRNA - SMARTpool | PAQR9 | Dharmacon | L-010642-000005 |
| RNAi | ON-TARGETplus Human PGR (5241) siRNA - SMARTpool | PR1/2 | Dharmacon | L-003433-000005 |
| RNAi | ON-TARGETplus Human AR (367) siRNA - SMARTpool | AR | Dharmacon | L-003400-000005 |
| RNAi | ON-TARGETplus Human NR3C1 (2908) siRNA - SMARTpool | GR | Dharmacon | L-003424-000005 |

**Table S3**

| Target Gene | Primer | Sequence | Product Size (bp) | Anneling Temp (°C) |
|---|---|---|---|---|
| **PR1/2** | PR-A/B-F1 | CGCGCTCTACCCTGCACTC | 121 | 65 |
| | PR-A/B-R1 | TGAATCCGGCCTCAGGTAGTT | | |
| **mPRα/PAQR7** | PAQR7-F1 | cgctcttctggaagccgtacatctatg | 122 | 65 |
| | PAQR7-R1 | cagcaggtgggtccagacattcac | | |
| **mPRβ/PAQR8** | PAQR8-F1 | agcctcctacata gatgctgccc | 194 | 65 |
| | PAQR8-R1 | ggtgcctggttcacatgttcttca | | |
| **mPRγ/PAQR5** | PAQR5-F1 | cagctgtttcacgtgtgtgtgatcctg | 144 | 65 |
| | PAQR5-R1 | gcacagaag tatggctccagctatctgag | | |
| **mPRδ/PAQR6** | PAQR6-F2 | gttgacccaccagcttagga | 176 | 65 |
| | PAQR6-R2 | atgccatcttcccagaacac | | |
| **mPRε/PAQR9** | PAQR9-F1 | TGCTACAAAGGGATCCCAAC | 202 | 65 |
| | PAQR9-R1 | TGGCACAGATGATTGGAAAA | | |
| **PGRMC1** | PGRMC1-F1 | CTGCATGATTTCTGTTTTATCTACCTCTA | 86 | 65 |
| | PGRMC1-R1 | TGTTACTGGACAGCGCTTAATCC | | |
| **AR** | AR-F1 | cct ggc ttc cgc aac tta cac | 168 | 65 |
| | AR-R1 | gga ctt gtg cat gcg gta ctc a | | |
| **GR/GCR** | GR-F1 | tc aaa aga gca gtg gaa gg | 260 | 65 |
| | GR-R1 | ggt agg ggt gag ttg tgg taa cg | | |

**Table S4**

All information of Table S4 is in Table 4 legend.

**Table S5, S6, S7 contain a large amount Omic raw and analytical data.**

**They were submitted in the compressed forms.**
